# Multi-omics prediction of oat agronomic and seed nutritional traits across environments and in distantly related populations

**DOI:** 10.1101/2021.05.03.442386

**Authors:** Haixiao Hu, Malachy T. Campbell, Trevor H. Yeats, Xuying Zheng, Daniel E. Runcie, Giovanny Covarrubias-Pazaran, Corey Broeckling, Linxing Yao, Melanie Caffe-Treml, Lucía Gutiérrez, Kevin P. Smith, James Tanaka, Owen A. Hoekenga, Mark E. Sorrells, Michael A. Gore, Jean-Luc Jannink

**Author notes:** Correspondence: Michael A. Gore, Jean-Luc Jannink.

## Abstract

Multi-omics prediction has been shown to be superior to genomic prediction with genome-wide DNA-based genetic markers (G) for predicting phenotypes. However, most of the existing studies were based on historical datasets from one environment; therefore, they were unable to evaluate the efficiency of multi-omics prediction in multi-environment trials and distantly-related populations. To fill those gaps, we designed a systematic experiment to collect omics data and evaluate 17 traits in two oat breeding populations planted in single and multiple environments. In the single-environment trial, transcriptomic BLUP (T), metabolomic BLUP (M), G+T, G+M and G+T+M models showed greater prediction accuracy than GBLUP for 5, 10, 11, 17 and 17 traits, respectively, and metabolites generally performed better than transcripts when combined with SNPs. In the multi-environment trial, multi-trait models with omics data outperformed both counterpart multi-trait GBLUP models and single-environment omics models, and the highest prediction accuracy was achieved when modeling genetic covariance as an unstructured covariance model. We also demonstrated that omics data can be used to prioritize loci from one population with omics data to improve genomic prediction in a distantly-related population using a two-kernel linear model that accommodated both likely casual loci with large-effect and loci that explain little or no phenotypic variance. We propose that the two-kernel linear model is superior to most genomic prediction models that assume each variant is equally likely to affect the trait and can be used to improve prediction accuracy for any trait with prior knowledge of genetic architecture.

## INTRODUCTION

Oat (Avena sativa L.) ranks sixth in world cereal production and has increasingly been consumed as a human food (USDA, 2019). Oat has a high content of health-promoting compounds such as unsaturated fatty acids, dietary fiber, antioxidants and vitamins, which makes it an interesting target for metabolomics studies from a human health and nutrition perspective (IMARC Group, 2019). In addition, high-density genetic markers have been developed in oat (Bekele et al., 2018), a draft genome sequence has been released (PepsiCo, 2020) and a high-quality and comprehensive seed transcriptome has been characterized (Hu et al., 2020). Furthermore, recent advances in high throughput sequencing and metabolite profiling technologies enable quantification of gene expression and metabolite abundance for hundreds of samples with high precision and reasonable cost (Alseekh & Fernie, 2018; Moll et al., 2014). All these advances in technology provides an opportunity to integrate different omics data and improve predictions for phenotypes of interest.

Several multi-omics prediction studies have been reported in cereal species (Guo et al., 2016; Riedelsheimer et al., 2012; Schrag et al., 2018; Wang et al., 2019; Westhues et al., 2017; Y. Xu et al., 2017; Yang Xu et al., 2021). These studies have shed light on the merits of multi-omics prediction over traditional genomic prediction and discussed useful statistical methods for integrating omics data. For instance, Y. Xu et al. (2017) and Wang et al. (2019) suggested that best linear unbiased prediction was the most efficient method compared to other commonly used genomic prediction and non-linear machine learning methods. However, most of those studies were based on historical datasets with a limited number of metabolite features and each level of omics data was collected from different environments. Therefore, they were unable to evaluate the efficiency of multi-omics prediction in multi-environment trials and genetically distant populations. However, in plant breeding, multi-environment trials are important for assessing the performance of genotypes across environments and identifying well-adapted genotypes for a specific region (Burgueño et al., 2012; Mathew et al., 2018). In addition, prediction of breeding values of distantly-related individuals are needed in many and perhaps the most promising applications of genomic selection in both plant and animal breeding programs (Lorenz & Smith, 2015; Meuwissen, 2009; Moghaddar et al., 2019).

To fill the knowledge gaps of multi-omics prediction in plant breeding, we designed a systematic experiment to collect omics data and evaluate eight agronomic and nine fatty acid traits (Supplemental Table 1) in a core set of a worldwide oat collection (termed Diversity panel) planted in one environment and advanced breeding lines adapted to the upper Midwest region in the U.S. (termed Elite panel) planted in three environments. Our efforts included (i) comparing the accuracy of mufti-omics prediction against genomic prediction in a single-environment trial; (ii) evaluating the efficiency of multi-omics prediction in multi-environment trials; and (iii) exploring the potential of using multi-omics data to predict distantly-related individuals.

## RESULTS AND DISCUSSION

After filtering out lines with low-quality genetic markers, the Diversity and Elite panels consisted of 368 and 232 lines (Supplemental Table 2), respectively, with 32 lines in common. A reconstructed phylogenetic tree revealed that the two panels were separated from each other in general, although some Diversity panel members were clustered to the Elite panel branches; and both panels showed population structure (Figure 1). This is consistent with our prior knowledge about origins of the two panels (Campbell et al., 2021).

**Figure 1.**
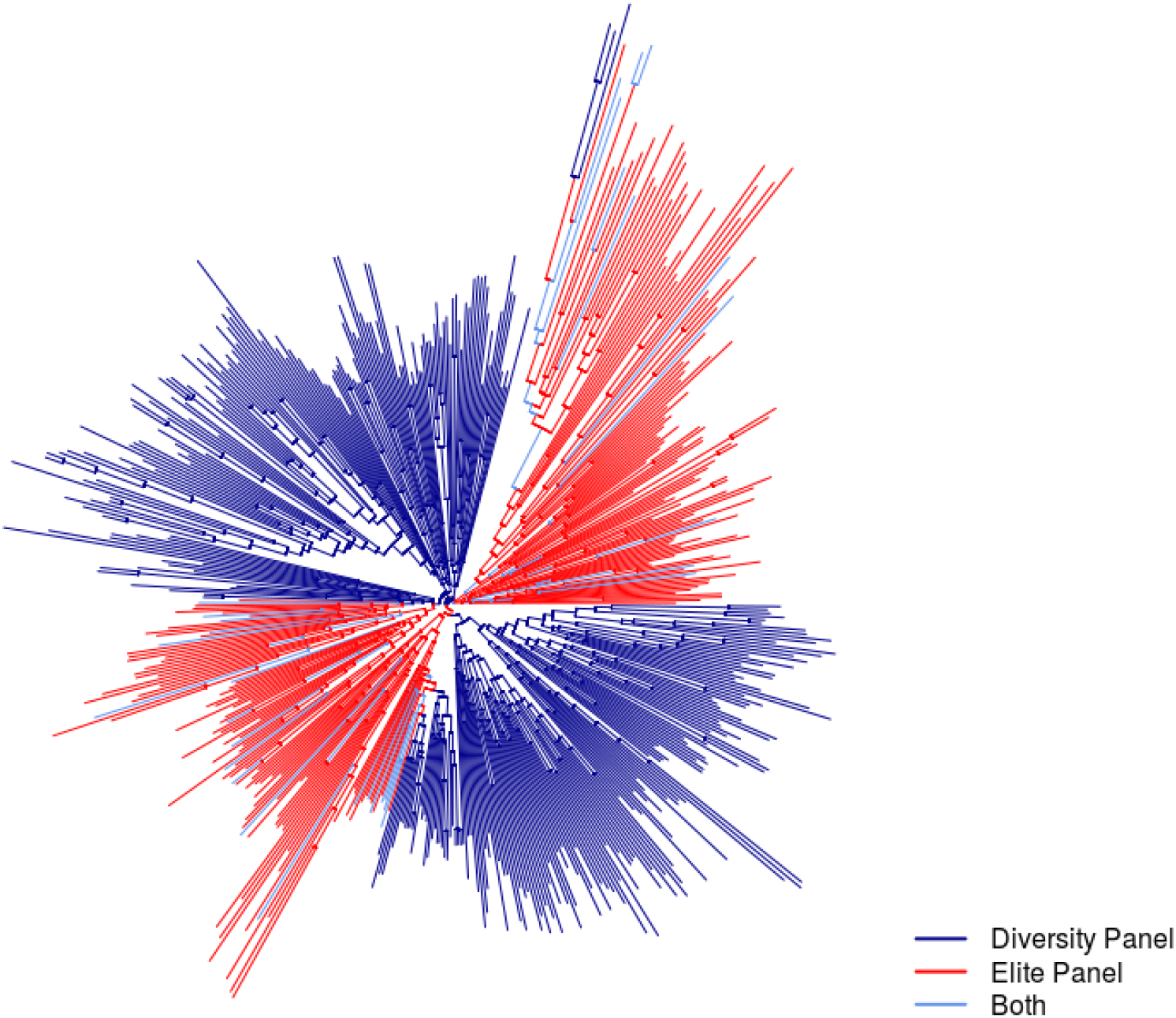
Neighbor-joining tree of 568 oat lines in the Diversity and Elite panels.

### Single-environment prediction in the Diversity panel

Using GBLUP (G) as a baseline, there were 5, 10, 11, 17 and 17 traits out of the 17 total traits with improved prediction accuracy from transcriptomic BLUP (T), metabolomic BLUP (M), G+T, G+M and G+T+M models, respectively (Figure 2, Supplemental Table 3). Percent change in prediction accuracy over GBLUP ranged from 0.1% (Days to Heading, G+T model) to 70.3% (C18:0, G+M model) with a median of 21.5%. Because GBLUP does not allow for large-effect or zero-effect genetic markers, we also compared BayesB with the multi-omics models, and found BayesB showed similar results to GBLUP (Supplemental Figure 1).

**Figure 2.**
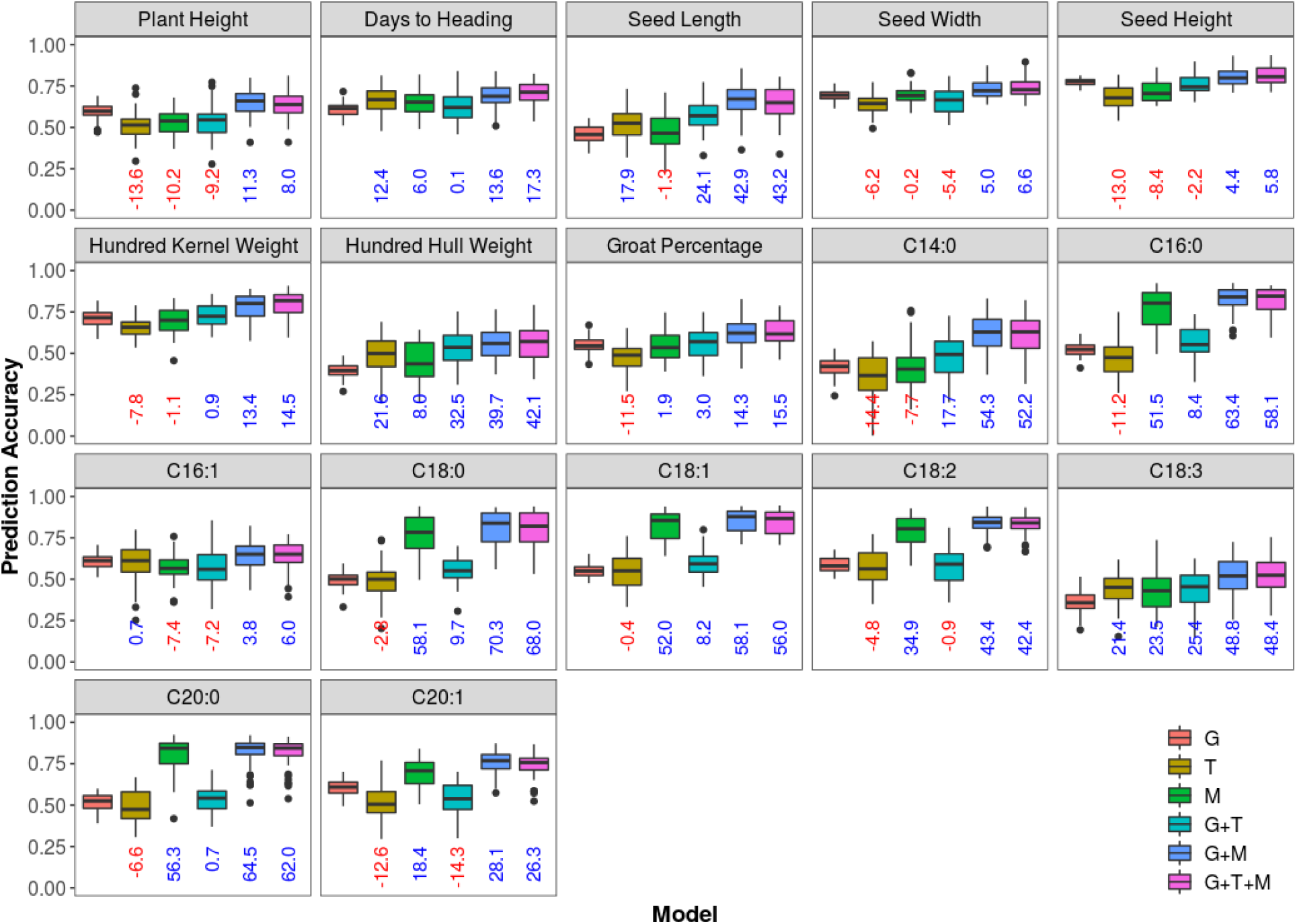
Distribution of prediction accuracy of the 17 phenotypic traits in the Diversity panel across 50 re-sampling runs. For each trait, boxplots with different colors represent prediction models, which are G, T, M, G+T, G+M and G+T+M from left to right. Medians of percent change in prediction accuracy of models relative to GBLUP are indicated below each box in blue if positive and in red if negative. G = genomic BLUP, T = transcriptomic BLUP, M = metabolomic BLUP.

To evaluate whether transcriptomic and metabolomic features equally contribute to improved prediction accuracy or if one is more important than the other, we compared multi-omics prediction models with T and M kernels added in different orders. By adding kernels in their order along the central dogma of molecular biology, median prediction accuracy changes from G to G+T models and from G+T to G+T+M models across all traits ranged from −11.6% to 35.8% (median=3.2%) and 6.5% to 55.6% (median=16.3%), respectively (Supplemental Figure 2). In contrast, when adding the M kernel first (G+M model) then followed by the T kernel (G+T+M model), percent changes in prediction accuracy ranged from 2.5% to 67.3% (median=41.7%) and −3.3% to 3.5% (median=−0.03%), respectively (Supplemental Figure 3). These results indicated that seed metabolites generally contributed more than transcripts to improving prediction accuracy of both agronomic and seed nutritional traits when combined with SNPs. Other researchers (Westhues et al., 2017; Y. Xu et al., 2017) reported that prediction abilities of transcripts were lower than GBLUP. The poor predictive performance of transcripts in existing studies might be explained by the fact that they were collected from a single time point and subject to dynamic changes in later unsampled developmental stages or by that transcripts and SNPs tend to capture similar genetic signals for predicted traits (Guo et al., 2016).

**Figure 3.**
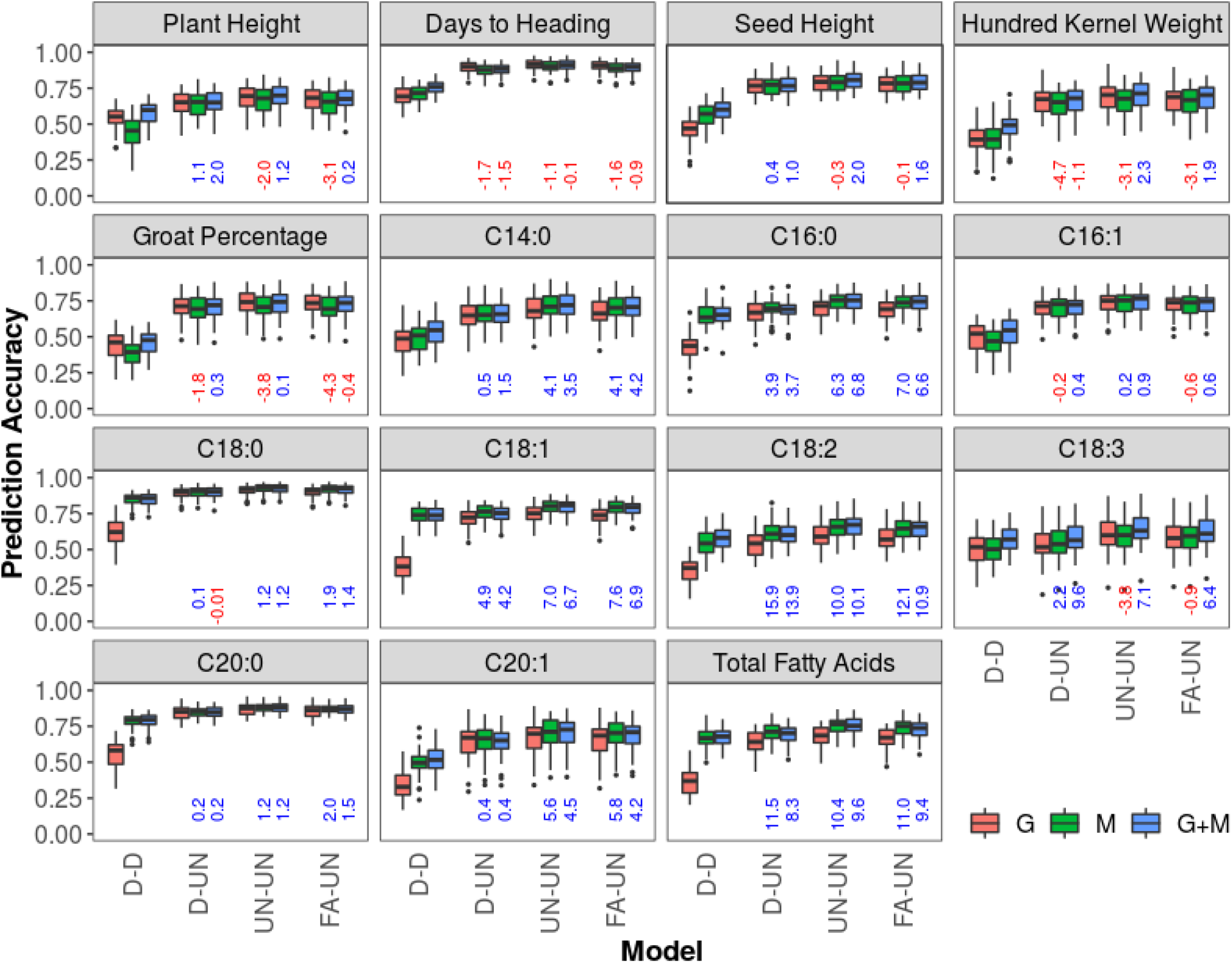
Distribution of prediction accuracy of the 15 phenotypic traits in the Elite panel across 50 re-sampling runs estimated by multi-trait models for multi-environment prediction. The 15 phenotypic traits in the Elite panel were evaluated at three environments. For each trait, boxplots with different colors represent models. Medians of percent change in prediction accuracy of M and G+M models relative to the G model are indicated below each box in blue if positive and in red if negative. For each model, the uppercase letters before and after the hyphen represent genetic and residual covariance structures: D=diagonal, UN=unstructured, FA=factor-analytic.

Although metabolites played important roles when combined with other kernels in improving prediction accuracy, we found that metabolites alone from mature seeds (M model) showed mixed results for predicting agronomic traits (Figure 2), while they greatly outperformed SNPs in predicting fatty acids. The relatively low performance of mature seed compounds in predicting agronomic traits might be explained by the fact that development of the agronomic traits and accumulation of compounds in mature seeds occurred either at different times or in different tissues. To further understand why metabolites are better predictors for fatty acid traits, we used the Weighted Gene Co-expression Network Analysis (WGCNA, Zhang & Horvath, 2005) that accommodated both annotated and unannotated compounds and used metabolites annotations (Supplemental Table 4) to elucidate their biological functions. We found 26 network modules and eight of them were enriched with lipids and lipid-like molecules (Supplemental Table 5), which included 33.0% of identified seed metabolite compounds. Those compounds directly or indirectly connected with fatty acids through biochemical pathways and different pathways relevant to lipids were likely influenced by overlapping gene sets. Therefore, they should be able to capture more genetic co-variation (including both additive and non-additive covariation) with fatty acids than SNPs fitted in an additive model. This hypothesis was partially supported by our results that combining G model and M model (G+M model) significantly improved prediction accuracies than using either model alone for all the 17 traits (Figure 2, Supplemental Table 6) and by findings of Guo et al. (2016) that adding metabolites to saturated SNP densities still led to significant increases in predictive abilities.

### Multi-environment prediction in the Elite panel

Beyond single-environment prediction, omics data might also have merit in predicting multi-environment trials, which has not yet been investigated to our knowledge. Here we used SNPs and metabolites for analyzing the multi-environment trials in the Elite panel, because transcript profiling from a single developmental time point showed limited value for improving prediction accuracy in addition to being very labor-intensive. We focused on prediction of lines that have been evaluated in some but not in target environments (CV2, Burgueño et al., 2012). To this aim, we applied a single environment cross validation method (Mathew et al., 2018) (Supplemental Figure 4). Briefly, to predict a phenotype in the first environment, we masked 20% of lines for cross validation and used metabolites from the other two environments to construct metabolomic relationship matrices to minimize the influence of non-genetic effects on prediction accuracy. We then used multi-trait models treating phenotypes from all three environments as separate traits for model training but using only the phenotype data of the masked lines from the first environment as the testing data. This procedure was repeated for the second and third environments and prediction accuracies were averaged across the three environments for each run.

Multi-environment predictions were performed using six multi-trait models (Supplemental Table 7) on three different kernels/combinations (G, M, G+M) with various genetic and residual covariance structures (Figure 3, Supplemental Figure 5). The diagonal heterogeneous covariance structure (D-D) corresponds to a single-environment model without borrowing information from other environments. The question that we explored was whether multi-omics models (M and G+M) could improve prediction accuracy compared to corresponding multi-trait models based on SNPs alone (G model). To answer this question, within each of the five multi-trait models (the D-D model was excluded), we compared percent change in prediction accuracy of M and G+M models relative to the G model. We found the M model outperformed the G model for all seed fatty acid traits except C16:1 and C18:3, with an increase in prediction accuracy ranging from 0.1 to 15.9%. However, the G+M model outperformed the G model for all traits except days to heading, with an increase in prediction accuracy over the G model ranging from 0.1 to 13.9%. These results confirmed the value of using multi-omics data in multi-environment prediction.

We then used the prediction accuracy from GBLUP in the single-environment model (D-D) as a baseline to compare the performance of different multi-trait models. We found that all multi-trait models outperformed their counterpart single-environment models (Figure 3, Supplemental Figures 6-8). The multi-trait models generally performed better when modeling the genetic covariance as unstructured (UN) or as factor-analytic (FA) than modeling genetic covariance as a diagonal structure (D). The highest prediction accuracy was achieved by either UN-D (UN and D represent genetic and residual covariance structures, respectively) or UN-UN models, although FA-D and FA-UN models provided very similar results. This indicated that the genetic covariance between environments played an important role in the multi-omics prediction models. These findings agree with recent genomic prediction studies (Malosetti et al., 2016; Montesinos-López et al., 2016) that UN covariance structure improved prediction accuracy compared to the models with diagonal homogeneous or heterogeneous covariances. Overall, we concluded that considering genetic and non-genetic covariances is useful to improve prediction accuracy of multi-environment models using multi-omics data.

### Using multi-omics data to improve genomic prediction in distantly-related populations

Although multi-omics data showed superiority over SNPs to predict phenotypes in both single and multi-environment trials, currently transcript and metabolite profiling is more expensive than SNP genotyping, which would limit their applications in plant breeding. Here we hypothesized that omics data from well characterized populations can be used to prioritize likely causal loci and improve performance of genomic prediction models in distantly-related populations. Seed fatty acid concentrations were used as target traits to test the hypothesis because their genetic architectures have been well characterized (Carlson et al., 2019) and lipid biosynthetic pathways are known to be highly conserved in higher plants (de Abreu e Lima et al., 2018).

To explore this scientific question, we first attempted to prioritize likely causal loci from the Diversity panel based on the eight network modules enriched with lipids and lipid-like molecules (Supplemental Table 5). Among the eight network modules, only one (darkred) strongly correlated with fatty acids (Supplemental Figure 9). We then performed hierarchical clustering and GWAS on eigenvectors of all the 26 network modules and PC1 of fatty acids. The eigenvector of the darkred module was clustered together with PC1 of fatty acids (Supplemental Figure 10) and had significant GWAS hits on chromosome 6A (Supplemental Figure 11), which co-located with the fatty acids major-effect QTL (*QTL-6A*, Supplemental Figure 12). However, the *QTL-6A* was not detected from other network modules. We further prioritized 140 markers including significant markers and the markers in LD with them based on the darkred module GWAS hits on chromosome 6A.

The primary use of locus prioritization is to split markers in the test population into two sets for a multi-kernel model prediction, in which the two genomic relationship kernels were constructed from the two marker sets. We termed our method multi-kernel network-based prediction (MK-Network) and found it improved prediction accuracy over GBLUP and BayesB for all fatty acid traits (Figure 4) except C14:0 and C18:3, because they had different genetic architectures from other fatty acids and no significant markers from GWAS (Supplemental Figure 12). The percent change of mean prediction accuracy over 50 cross-validation runs ranged from 4.0% to 32.0% with a mean of 14.5%.

**Figure 4.**
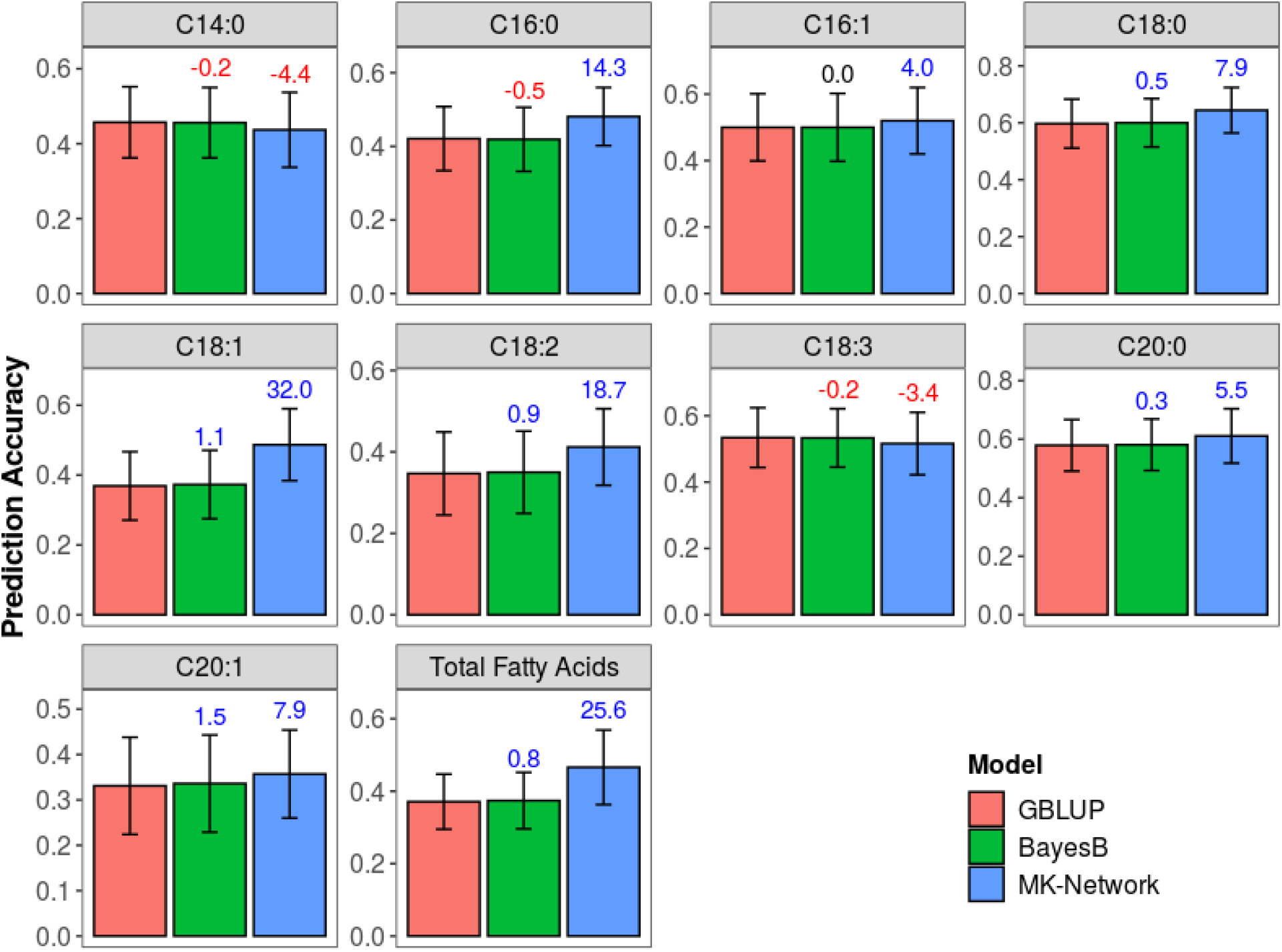
Prediction accuracy of the 10 fatty acid traits in the Elite panel estimated by GBLUP, BayesB and two-kernel BLUP models across 50 re-sampling runs. For each trait, barplots with different colors represent models. Means of percent change in prediction accuracy of all other models relative to GBLUP are indicated above each bar (in blue if positive, in red if negative, and in black if zero). MK-Network=network-based multiple-kernel prediction.

The universal QTL of fatty acids (*QTL-6A*, Supplemental Figures 12-13) and similar LD relationships (Supplemental Figure 14) with the surrounding loci between the Diversity and Elite panels promoted the success of our likely causal loci prioritization. The network-based prioritization strategy takes advantages of pleiotropy, in which one or a few genes influence both target traits and other metabolites from related network modules. In the darkred module, 23 of 32 metabolites showed clear peaks at the *QTL-6A*, although only five of them were significant at FDR<0.05 (Supplemental Figure 15). This indicated that *QTL-6A* was likely a causal locus and influenced both fatty acids and the darkred module. The relationships between fatty acids and the darkred module are expected to be conserved between populations. However, we were unable to test this because there is currently no robust method to map all untargeted metabolites from one panel to another and quantify them precisely.

Most genomic prediction methods assume that each variant is equally likely to affect the trait (MacLeod et al., 2016). There are certain loci that explain more phenotypic variance and they should be placed in different kernels than loci that explain little or no variance. However, the other kernel is still needed because we may unintentionally exclude important loci based on prior biological knowledge alone, for example, a prior GWAS might not identify all possible causal loci. There are many loci that have small effects, through whatever pathway, whether it is through trans effects as hypothesized in the omnigenic model (Liu et al., 2019) or through much more indirect effects like competition for photosynthates or impact on fitness (Price et al., 2018). Li et al. (2018) found that excluding those small-effect loci could not further improve prediction accuracy compared to GBLUP with all SNPs. Therefore, a two-kernel linear model that accommodates both likely casual loci and loci with minimal to no effect should be used to improve prediction accuracy for any traits with prior knowledge of genetic architecture.

## METHODS

### The plant materials and experimental designs

The Diversity and Elite panels consisted of 378 and 252 lines (Supplemental Table 2), respectively. The Diversity panel originally included 500 lines described by Carlson et al. (2019) that was a core set of worldwide collection of oat germpalsm, and we further selected for lines with visible anther extrusion. The Diversity panel was planted at Ithaca, NY, and the Elite panel was planted at Madison, WI, Crookston, MN, and Brookings, SD, respectively. An augmented incomplete design was used for both panels. The Diversity panel included 18 blocks of 23 plots each, one common check across all blocks and six secondary checks replicated in three blocks each. The Elite panel included 12 blocks of 25 plots each, one common check across all blocks and two secondary checks replicated in six blocks each.

### Phenotype evaluation and analysis

Plant height was evaluated for five randomly selected plants in each plot after anthesis. Days to heading was defined by the days from seeding to heading in >50% of total plants. 100 randomly selected seeds from each plot were dehulled with a hand dehuller for evaluation of hundred kernel weight, hundred hull weight and groat percentage. After dehulling, 50 randomly selected seeds were delivered to the Proteomics and Metabolomics Facility at Colorado State University for metabolite analysis, and the other 50 seeds were used for measuring seed length, width and height with an electronic micrometer. Fatty acids were identified and quantified with targeted GC-MS, then normalized to concentration (mg/g of oats) against the internal standard (C17:0) (details were described in the Supplemental Methods).

### Genotype analysis

Genotypic data of the two panels were downloaded from T3/oat (https://triticeaetoolbox.org/oat/). Marker quality control followed Calson et al. (2019) and there were 73,014 markers and 568 lines (368 for the diversity panel, 232 for the elite panel, 32 in common) left after filtering. Subsequently, missing genotypes were imputed using the linear regression method glmnet described by Chan et al. (2016). The imputed genotypic data was used for constructing a neighbor-joining tree based on Rogers’ distance using the ape package (Paradis et al., 2004).

### Transcript profiling

RNAseq was based on developing seeds at 23 days after anthesis (DAA). The 23 DAA was chosen based on our pilot study (Hu et al., 2020) that showed 23 DAA had slightly higher correlation between transcript and metabolite abundance than other sampled developmental time points. Seed sample collection, RNA extraction, library construction procedures were described in details by Hu et al. (2020). Pooled libraries were sequenced using Illumina NextSeq500 with a 150 nt single-end run. The RNAseq reads quality trimming, transcript abundance quantification, and library size normalization following Hu et al.(2020).

### Metabolite profiling

Metabolite analysis was based on physiologically mature seeds because they have the highest level of health-promoting compounds and those compounds are stable at room temperature until germination. GC-MS non-targeted analysis and LC-MS Phenyl-Hexyl analysis were done at the Proteomics and Metabolomics Facility at Colorado State University. Details of chemical analysis, raw mass spectrometry data processing, metabolite annotation and normalization were described in the Supplemental Methods. The normalized metabolomics data was used for network analysis with WGCNA (Zhang & Horvath, 2005).

### Analysis of phenotypic traits, transcriptomic and metabolic features

Phenotypic traits, transcriptomic and metabolic features were analyzed following a standard linear mixed model of an augmented design accounting for effects of check genotypes and blocks. For metabolites analysis, batch effect was also included in the model. All statistical models were described in the Supplemental Methods and fitted using the sommer package (Covarrubias-Pazaran, 2016).

### Single-environment prediction

The additive genomic relationship matrix was made with the rrBLUP package (Endelman, 2011), and relationship matrices for transcriptomics and metabolomics data were made following Westhues et al. (2017). GBLUP, Transcriptomic BLUP (T), metabolomic BLUP (M), G+T, G+M and G+T+M models were fitted with the BGLR package (Pérez & De Los Campos, 2014). In the Diversity panel, transcriptomics and metabolomics data were collected on the same plots as the phenotypic data and therefore non-genetic (i.e., microenvironmental) factors that affected both omics features and phenotypic traits may induce non-genetic correlations among traits. Therefore, we estimated prediction accuracy as 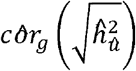 described by Runcie and Cheng (2019), and used a 50:50 training:testing split of the data to ensure that *côr* _*g*_ could be estimated accurately in the testing partition. This cross-validation procedure was repeated for 50 times with different random partitions.

### Multi-environment prediction

The metabolomics data were collected on the same plots as the phenotypic data in the Elite panel, which would bias prediction accuracy if directly using metabolites to predict target phenotypes in the same environment. Therefore, when predicting target phenotypes from one environment, we used metabolites from other two environments to make metabolomic relationship matrix. For each trait, we fitted six multi-trait mixed models on G, M and G+M kernels with different genetic and residual covariance structures (Supplemental Table 7). We applied a single environment cross validation method for genomic prediction described by (Mathew et al., 2018). Procedure of the single environment cross validation was illustrated in Supplemental Figure 4 and described in detail in the Supplemental Methods.

### Prediction of distantly related individuals

Prediction of distantly related individuals included two steps: likely causal loci prioritization and multiple-kernel prediction. We first performed likely causal locus prioritization from the Diversity panel based on the eight network modules enriched with lipids and lipid-like molecules, then utilized the prioritized markers and all rest markers to construct two genomic relationship kernels for a multiple-kernel prediction in the Elite panel. The details of related analyses were described in the Supplemental Methods.

## AUTHOR CONTRIBUTIONS

J.J., M.A.G and M.E.S designed the research. H.H. analyzed the data. H.H., M.T.C, M.A.G and J.J. wrote the manuscript. D.E.R, G.C., O.A.H and M.E.S advised H.H. on data analysis. H.H, T.H.Y, X.Z., M.C., L.C., K.P.S. J.T. performed experiments. C.B. and L.Y. performed metabolite analysis. All co-authors were involved in editing the manuscript.

## ACKNOWLEDGMENTS

We thank Joshua Wood and Robin Buell for helping with oat seed RNA extraction; David Benscher, Amy Tamara Fox and Nicholas Kaczmar for help with planting and harvesting field trials and sample collection; Yujie Meng for phenotype evaluation; Jing Wu and Peter Schweitzer for library preparation and RNA sequencing.

