## Supplemental Information for "Multi-omics prediction of oat agronomic and seed nutritional traits across environments and in distantly related populations"

The Supplemental Information includes:

- Supplemental Methods
- Supplemental Figures
- Supplemental Tables

### Supplemental Methods

#### Metabolic profiling

#### Analysis of phenotypic traits, transcriptomic and metabolic features

#### Multi-environment prediction

#### Prediction of distantly related individuals

#### Metabolic profiling

**Extraction**. Samples were homogenized and extracted using a biphasic extraction method to separate polar and non-polar compounds. Samples were homogenized using the Next Advance Bullet Blender system (Troy, NY) in which the intact oats were first crushed in 5-mL polypropylene tubes before the addition of stainless steel beads followed by repetitive concussive forces that grind the oats into a fine powder. The homogenized tissue (100 mg) was weighed into 2-mL glass vials for automated biphasic extraction using a GERSTEL Dual-head MPS fitted with 2.5 mL and 100 µL syringes (GERSTEL; Linthicum, MD).

The extraction solvent methanol/methyl-tert-butyl-ether (MTBE) (1.155 mL, 2:1, v/v) was added to each sample followed by vortexing for 30 min at 800 rpm on MPS. To induce biphasic separation, 640 µL of water was added to each vial followed by an additional 15 minutes of vortexing at 800 rpm. The samples were then centrifuged at 3,000 *g* for 15 min at 4°C and put back onto the MPS system. 60 µL of the upper organic layer from each sample was pooled to create a QC mix. 600 µL of the upper organic layer was transferred to clean 2-mL glass vials. To remove the remaining organic layer in the extraction vial that could not be selectively removed by the syringe needle, 600 µL of MTBE was added back to the original vial slowly as to not disturb the biphasic boundary and then 600 µL was subsequently removed and pooled with the rest of the organic layer. This addition and removal processes was repeated three times to ensure all organic phase was recovered. 600 µL of methanol/acetonitrile (1:1, v/v) was added to the remaining aqueous layer in the extraction vial and vortexed for 3 min to precipitate proteins and excess glycans followed by centrifugation at 3,000 *g* for 15 min at 4°C. 60 µL of aqueous extract was then removed and pooled to create an aqueous layer QC mix before 1.7 mL was transferred to new 2 mL glass vials. Both the organic and aqueous extracts were dried under nitrogen gas overnight. Aqueous fractions were re-suspended in 1.5 mL of 50% methanol in water (v/v) for GC-MS and LC-MS untargeted analysis. Organic fractions were re-suspended in 0.7 mL of methanol/toluene (1:1, v/v) for FAME analysis.

**LC-MS Phenyl-Hexyl Analysis**. 5 µL of aqueous extract was injected onto a Waters Acquity UPLC system with a pooled QC injection after every 5 sample injections. The metabolites were separated using a Waters Acquity UPLC CSH Phenyl Hexyl column (1.7 µm, 1.0 x 100 mm), using a gradient from solvent A (2mM ammonium formate with 0.1% formic acid) to solvent B (acetonitrile with 0.1% formic acid). The gradient started with 100% A, held at 100% A for 1 min, ramped to 98% B over 12 min, held at 98% B for 3 min, and then returned to starting conditions over 0.05 min and allowed to re-equilibrate for 3.95 min, with a 200 µL/min constant flow rate. The column and samples were held at 65°C and 6°C, respectively. The column eluent was analyzed with a Waters Xevo G2 TOF-MS with an electrospray source in positive mode (cone voltage =30 V), scanning 50-2000 m/z at 0.2 s per scan, alternating between MS (6 V collision energy) and MSE mode (15-30 V ramp). Calibration was performed using sodium iodide with 1 ppm mass accuracy. The capillary voltage was held at 2200 V, source temp at 150°C, and nitrogen desolvation temperature at 350°C with a flow rate of 800 L/hr.

**GC-MS Non-targeted Analysis.** 40 µL of the aqueous extract was dried under nitrogen, re-suspended in 50 µL of pyridine containing 25 mg/mL of methoxyamine hydrochloride, incubated at 60°C for 1 h, sonicated for 10 min, and incubated for an additional 1 h at 60°C. Next, 50 µL of N- methyl-N-trimethylsilyltrifluoroacetamide with 1% trimethylchlorosilane (MSTFA + 1% TMCS, Thermo Scientific) was added and samples were incubated at 60°C for 45 min, briefly centrifuged, cooled to room temperature, and 100 µL of the supernatant was transferred to a 150-µL glass insert in a GC-MS autosampler vial. Metabolites were detected using a Trace 1310 GC coupled to a Thermo ISQ mass spectrometer (Thermo Scientific). Samples were injected in a 1:10 split ratio. Separation occurred using a 30 m TG-5MS column (Thermo Scientific, 0.25 mm i.d., 0.25 µm film thickness) with a 1.2 mL/min helium gas flow rate, and the program consisted of 80˚C for 30 s, a ramp of 15°C/min to 330°C, and an 8 min hold. Masses between 50-650 m/z were scanned at 5 scans/s after electron impact ionization.

**FAME preparation.** Samples were re-suspended in 0.7 mL of methanol/toluene (1:1, v/v) and 70 µL was transferred to a 2-mL glass vial and solvent was completely removed by nitrogen evaporation at ambient temperature. To the dry sample, 100 µL of toluene containing 2.5 mg/mL of internal standard, glyceryl triheptadecanoate, and 200 µL of 3 N methanolic HCl were added. The mixture was incubated at 60 °C for 1 hour. Then 0.5 mL of hexane and 0.7 mL of water were added to the cooled sample. After a brief vortex, the sample was centrifuged at 3000 *g* for 5 min at 4°C. After centrifugation, the upper hexane layer was diluted twice with 100% hexane and 100 µL transferred to glass inserts for analysis.

**FAME GCMS.** Upper hexane layer containing FAME (1 µL) was injected into a TG-WAXMS column (30 m x 0.25 mm x 0.25 µm, Thermo) on a Trace1310 GC (Thermo) coupled to a Thermo ISQ-LT MS. The injector temperature was 260 °C, and split ratio was 15:1. A constant flow rate of the carrier gas (Helium) was controlled at 1.2 mL/min. The initial oven temperature was 200°C and held for 1 min, then increased to 260 °C at 10 °C/min and held for 3 min. Detection was completed under electron impact mode, with a scan range of 50-650 m/z and scan rate 5 scans/s. Transfer line and source temperature were both at 250°C. Data processing was completed with Chromeleon 7 software (ThermoFisher). QC sample were injected after every 6 samples. Standard curves of C14:0, C16:1, C16:0, C18:0, C18:1, C18:2, C18:3, C20:0, and C20:1 were acquired.

**Data Processing**. For each sample, raw data files were converted to .cdf format, and matrix of molecular features as defined by retention time and mass (m/z) was generated using XCMS software in R (Smith et al., 2006) for feature detection and alignment. The matchedFilter algorithm was used for GC-MS data, and the centWave algorithm for LC-MS data. Features were grouped using RAMClustR (Broeckling et al., 2014), with normalization set to ‘none’. GC-MS spectra were annotated by matching unknown spectra to the GOLM metabolome retention indexed spectral library (Kopka et al., 2005), using retention times plotted vs the GOLM retention index to increase confidence in the spectral match. Searching was accomplished using the RAMSearch program (Broeckling et al., 2016).

LC-MS data were first annotated by searching against an in-house spectra and retention time database using RAMSearch. RAMClustR was used to call the findMain (Jaeger et al., 2017) function from the interpretMSSpectrum package to infer the molecular weight of each LC-MS compound and annotate the mass signals. The complete MS spectrum and a trunctated MSE spectrum were written to a .mat format for import to MSFinder (Tsugawa et al., 2016). The MSE spectrum was truncated to only include masses with values less than the inferred M plus its isotopes, and the .mat file precursor ion is set to the M+H ion for the findMain inferred M value. These .mat spectra were analyzed to determine the most probable molecular formula and structure. MSFinder was also used to perform a spectral search against the MassBank database. All results were imported into R and a collective annotation is derived with prioritization of RAMSearch > MSFinder mssearch > MSFinder structure > MSFinder formula > findMain M. Annotation confidence is reported as described (Sumner et al., 2007). All R work was performed using R 3.3.1 (R Core Team, 2017).

**Normalization.** The original GC-MS and LC-MS spectra abundances were normalized using the Log Transformation method as described by Li et al. (2016). Then, to further control batch effect, the log transformed spectra abundances were fitting in a linear mixed model including batch number as a covariate (described on the next page).

#### Analysis of phenotypic traits, transcriptomic and metabolic features

Linear mixed model for analyzing agronomic traits in a single-environment trial:

y = μ + Check + Block + New:Entry + e (Eq.1)

Linear mixed model for analyzing metabolites in a single-environment trial:

y = μ + Check + Block + Batch + New:Entry + e (Eq.2)

Linear mixed model for analyzing metabolites in multi-environment trials:

y = μ + Check + Loc + Loc:Block + Batch + New:Entry + Loc:New:Entry+ e (Eq.3)

where Check is a fixed effect for check varieties; Loc is a random location effect; Block and Batch are random effects to account for field blocks and injection batch for GCMS/LCMS; Loc:Block is Block effect nested within location; New is an indicator variable where 0 indicates a check variety and 1 indicates a test entry, and is nested within entry; Loc:New:Entry is random effect of location by entry interaction. The terms μ and e represent the overall mean and the vector of residuals, respectively. The above models were fitted using the sommer package in R (Covarrubias-Pazaran, 2016). Best linear unbiased predictors (BLUPs) were calculated for each variety for downstream analysis.

#### Multi-environment prediction

The metabolomics data were collected on the same plots as the phenotypic data in the Elite panel, which would bias prediction accuracy if directly using metabolites to predict target phenotypes of the same environment. Therefore, when predicting target phenotypes from one environment, we used metabolites from other two environments to make metabolomic relationship matrix. For each trait, we fitted six multi-trait mixed models on G, M and G+M kernels with different genetic and residual covariance structures (Supplemental Table 7). We applied a single environment cross validation method for genomic prediction described by (Mathew et al., 2018) and extended it to multi-kernel omic prediction (illustrated in Supplemental Figure 4). To predict a phenotype in the first environment, we masked 20% of lines for cross validation and used metabolites from the other two environments to construct metabolomic relationship matrices. We then used multi-trait models treating phenotypes from all three environments as separate traits for model training but using only the phenotyped data of the masked lines from the first environment as the testing data. We further estimated prediction accuracy of the first environment as $r\,(\hat{y}, y)\Big/\sqrt{h^2}$ (Riedelsheimer et al., 2012), where r(ŷ,y) is the Pearson correlation between the observed (y) and predicted (ŷ) phenotypic values and $h^2$ is the heritability of the target trait. To predict the phenotype in the second and third environments, we masked 20% of lines (the same genotypes as those in the first environment) from the second and third environments, respectively, and calculated their prediction accuracies following the same procedure as that applied to the first environment.

Finally, we averaged the three prediction accuracies across environments to represent the prediction accuracy of a single run. This procedure was repeated for 50 times with different random partitions.

#### Prediction of distantly related individuals

Prediction of distantly related individuals included two steps: likely causal loci prioritization in the Diversity panel and multiple-kernel prediction in the Elite panel.

We first performed WGCNA on all metabolite features in the Diversity panel, and performed module-trait correlation analysis between eigenvectors of the network modules and fatty acids traits. Based on the identified network modules and metabolites annotation, we further performed Fisher's exact test to identify modules enriched with lipids and lipid-like molecules. We then performed hierarchical clustering and GWAS on eigenvectors of the twenty-six network modules and PC1 of fatty acids. The darkred module was found enriched with lipids and lipid-like molecules, clustered together with PC1 of fatty acids and its eigenvector had a QTL co-located with fatty acids major-effect QTL on chromosome 6A. We further prioritized 140 markers including significant markers and the markers in LD with them based on the darkred module GWAS hits on chromosome 6A. A LD threshold of $r^2$=0.1 was used as it is frequently recommended for SNP pruning (Kawakami et al., 2014).

The prioritized markers and all rest markers were used to construct two genomic relationship kernels in the Elite panel and perform a multiple kernel prediction. Genomic predictions with GBLUP and BayesB models were used as references to compare with the two-kernel linear model. The five-fold cross-validation was ued to estimate prediction accuracies for all models and the prediction accuracy was estimated as $r\,(\hat{y}, y)\Big/\sqrt{h^2}$ (Riedelsheimer et al., 2012). This cross-validation procedure was repeated for 50 times with different random partitions.

#### Supplemental Figures

**Supplemental Figure 1.** Distribution of prediction accuracy of the 17 phenotypic traits in the Diversity panel across 50 re-sampling runs estimated by multi-omics and BayesB models.

**Supplemental Figure 2.** Prediction accuracy changes from G to G+T models and from G+T to G+T+M models of the 17 phenotypic traits in the Diversity panel across 50 re-sampling runs.

**Supplemental Figure 3.** Prediction accuracy changes from G to G+M models and from G+M to G+T+M models of the 17 phenotypic traits in the Diversity panel across 50 re-sampling runs.

**Supplemental Figure 4.** a scheme of single environment cross validation using metabolites (M model) or SNPs and metabolites (G+M model) for multi-environment prediction.

**Supplemental Figure 5.** Distribution of prediction accuracy of the 15 phenotypic traits in the Elite panel across 50 re-sampling runs estimated by multi-trait models of D-D, UN-D and FA-D.

**Supplemental Figure 6.** Percentage changes in prediction accuracy estimated from multi-environment GBLUP models over single-environmental GBLUP for the 15 phenotypic traits in the Elite panel

**Supplemental Figure 7.** Percentage changes in prediction accuracy estimated from multi-environment metabolite BLUP (M) models over single-environmental GBLUP for the 15 phenotypic traits in the Elite panel

**Supplemental Figure 8.** Percentage changes in prediction accuracy estimated from multi-environment genomic and metabolomic BLUP (G+M) models over single-environmental GBLUP for the 15 phenotypic traits in the Elite panel.

**Supplemental Figure 9.** Correlation between eigenvector of network modules and fatty acid traits.

**Supplemental Figure 10.** Hierarchical clustering dendrogram of the network eigenvectors, PC1 and PC2 of the nine fatty acids.

**Supplemental Figure 11.** Manhattan plots of eigenvectors of twenty-six network modules identified by WGCNA and PC1 of fatty acids in the Diversity panel.

**Supplemental Figure 12.** Manhattan plots of fatty acids traits in the Diversity panel.

**Supplemental Figure 13.** Manhattan plots of fatty acids traits in the Elite panel.

**Supplemental Figure 14.** LD relationships of locus QTL-6A with the surrounding loci in the Diversity and Elite panels.

**Supplemental Figure 15.** Manhattan plots of metabolites in the darkred module identified by WGCNA in the Diversity panel.

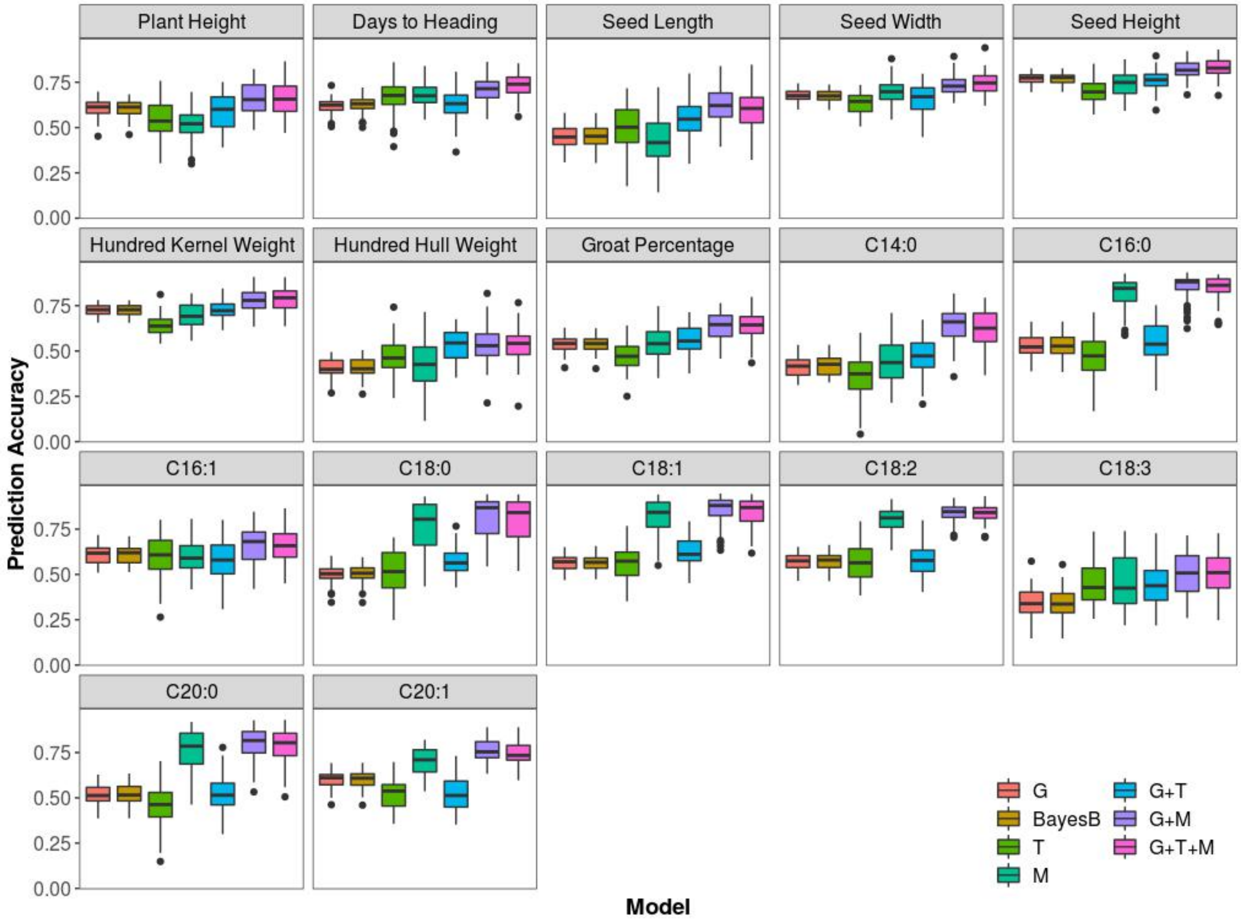


**Supplemental Figure 1.** Distribution of prediction accuracy of the 17 phenotypic traits in the Diversity panel across 50 re-sampling runs estimated by multi-omics and BayesB models. For each trait, boxplots with different colors represented prediction results estimated by different models. G = genomic BLUP, T = transcriptomic BLUP, M = metabolomic BLUP.

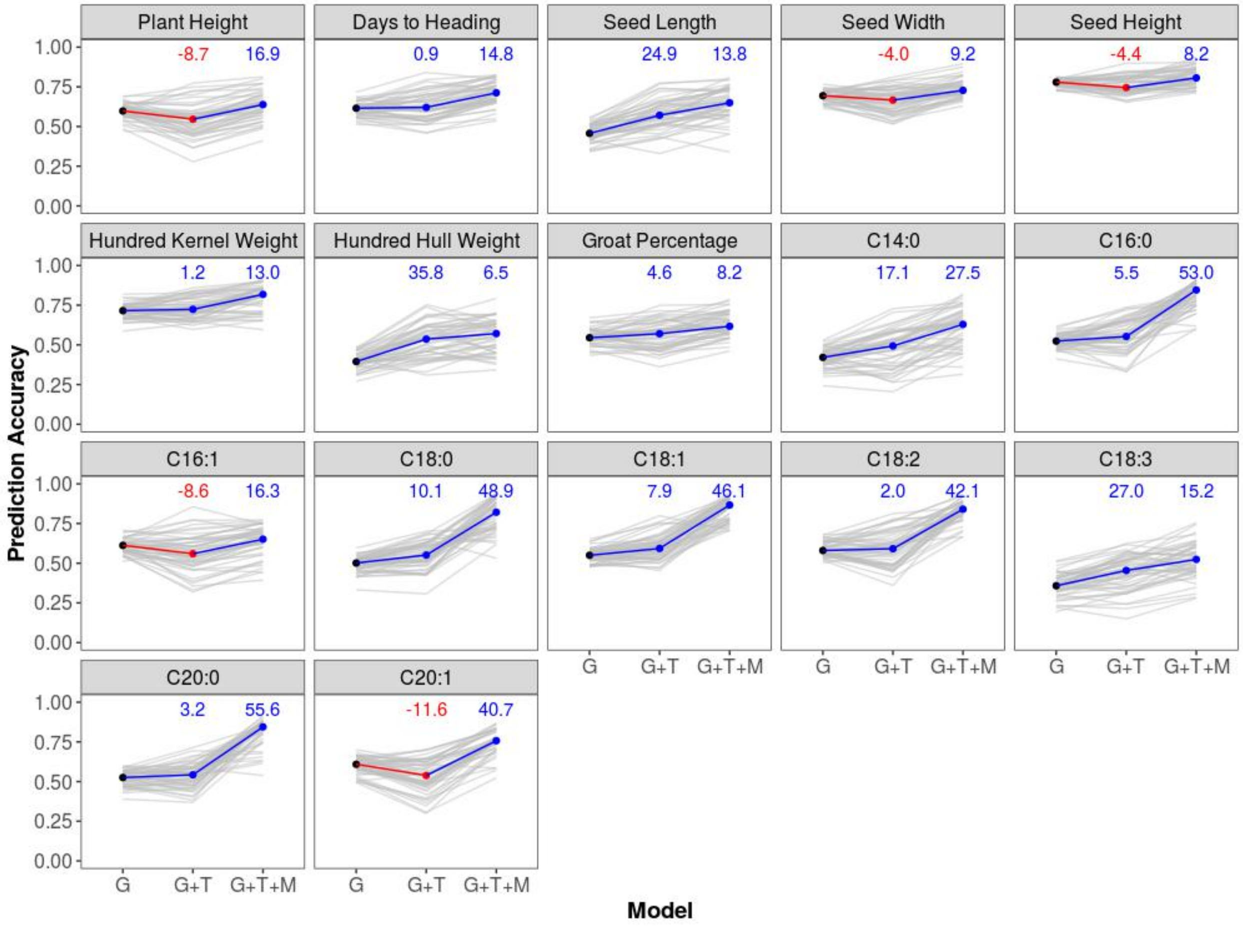


**Supplemental Figure 2.** Prediction accuracy changes from G to G+T models and from G+T to G+T+M models of the 17 phenotypic traits in the Diversity panel across 50 re-sampling runs. Each gray line represents a re-sampling run, and colored lines represent median prediction accuracy across the 50 re-sampling runs. Medians of percent change in prediction accuracy of models relative to a previous reduced model are indicated in blue if positive and in red if negative on top of each box. G = genomic BLUP, T = transcriptomic BLUP, M = metabolomic BLUP.

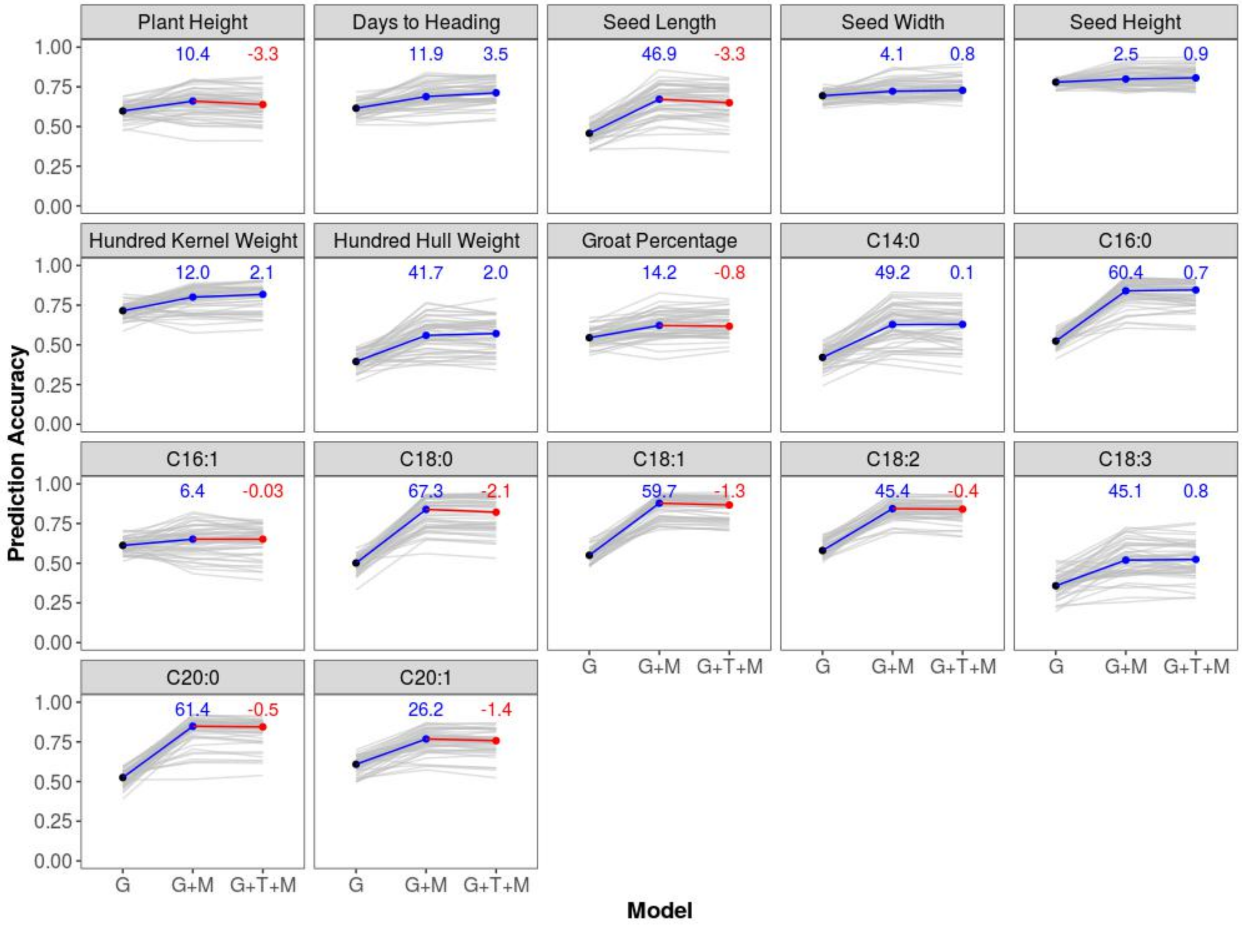


**Supplemental Figure 3.** Prediction accuracy changes from G to G+M models and from G+M to G+T+M models of the 17 phenotypic traits in the Diversity panel across 50 re-sampling runs. Each gray line represents a re-sampling run, and colored lines represent median prediction accuracy across the 50 re-sampling runs. Medians of percent change in prediction accuracy of models relative to a previous reduced model are indicated in blue if positive and in red if negative on top of each box. G = genomic BLUP, T = transcriptomic BLUP, M = metabolomic BLUP.

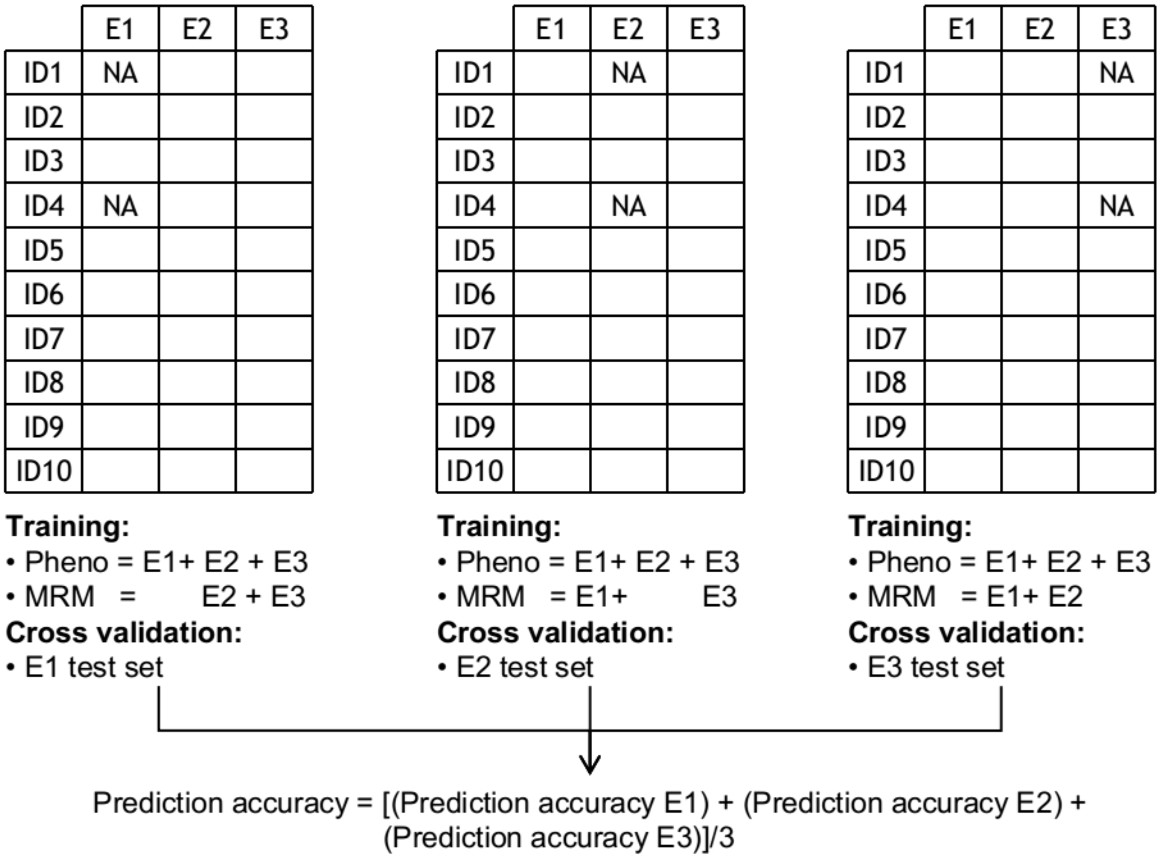


**Supplemental Figure 4.** a scheme of single environment cross validation using metabolites (M model) or SNPs and metabolites (G+M model) for multi-environment prediction. E1= Environment 1, E2=Environment 2, E3=Environment 3, Pheno=Phenotype, MRM=metabolomic relationship matrix.

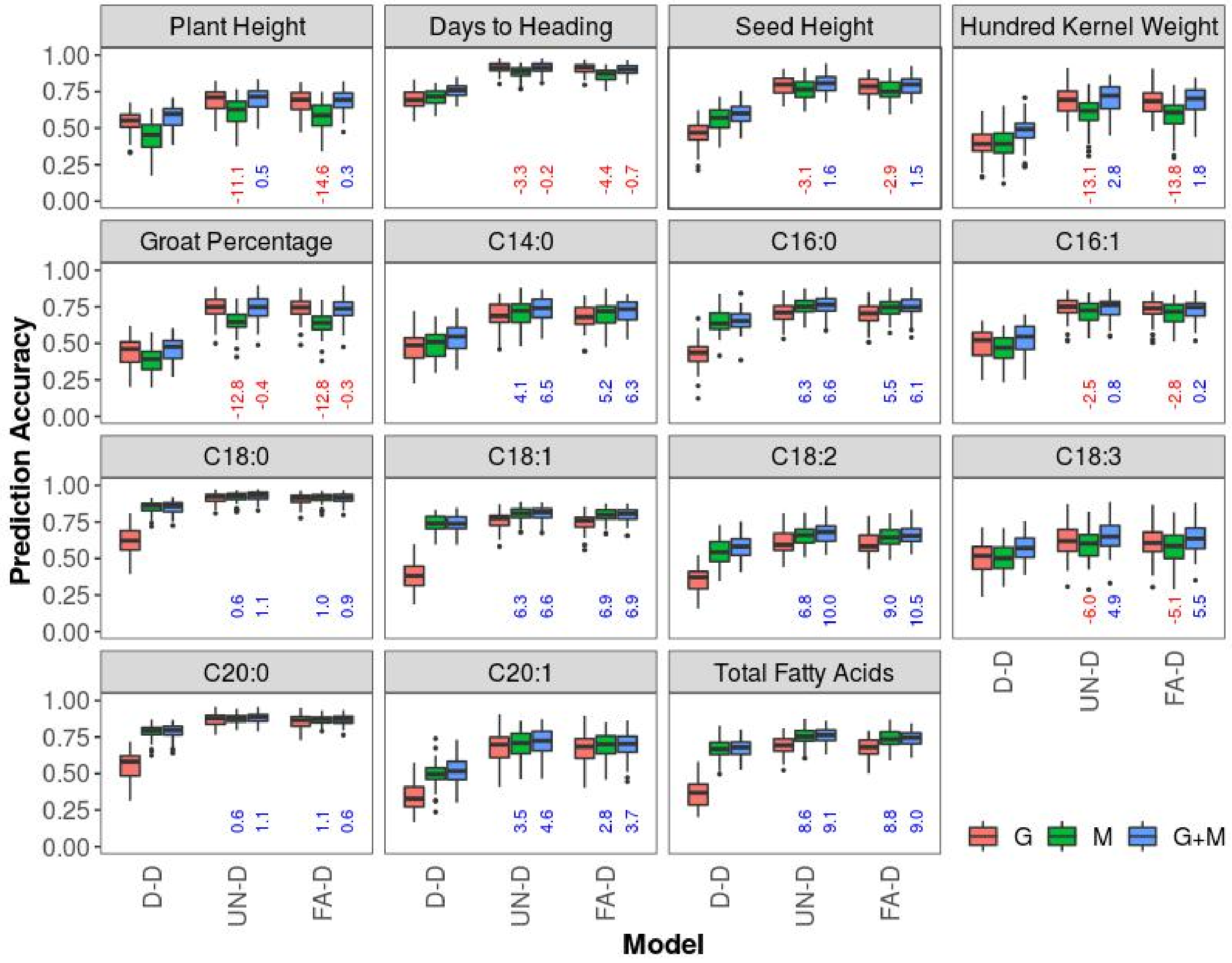


**Supplemental Figure 5.** Distribution of prediction accuracy of the 15 phenotypic traits in the Elite panel across 50 re-sampling runs estimated by multi-trait models of D-D, UN-D and FA-D. For each trait, boxplots with different colors represent models. Medians of percent change in prediction accuracy of M and G+M models relative to the G model are indicated below each box in blue if positive and in red if negative. For each model, the uppercase letters before and after the hyphen represent genetic and residual covariance structures: D=diagonal, UN=unstructured, FA=factor-analytic.

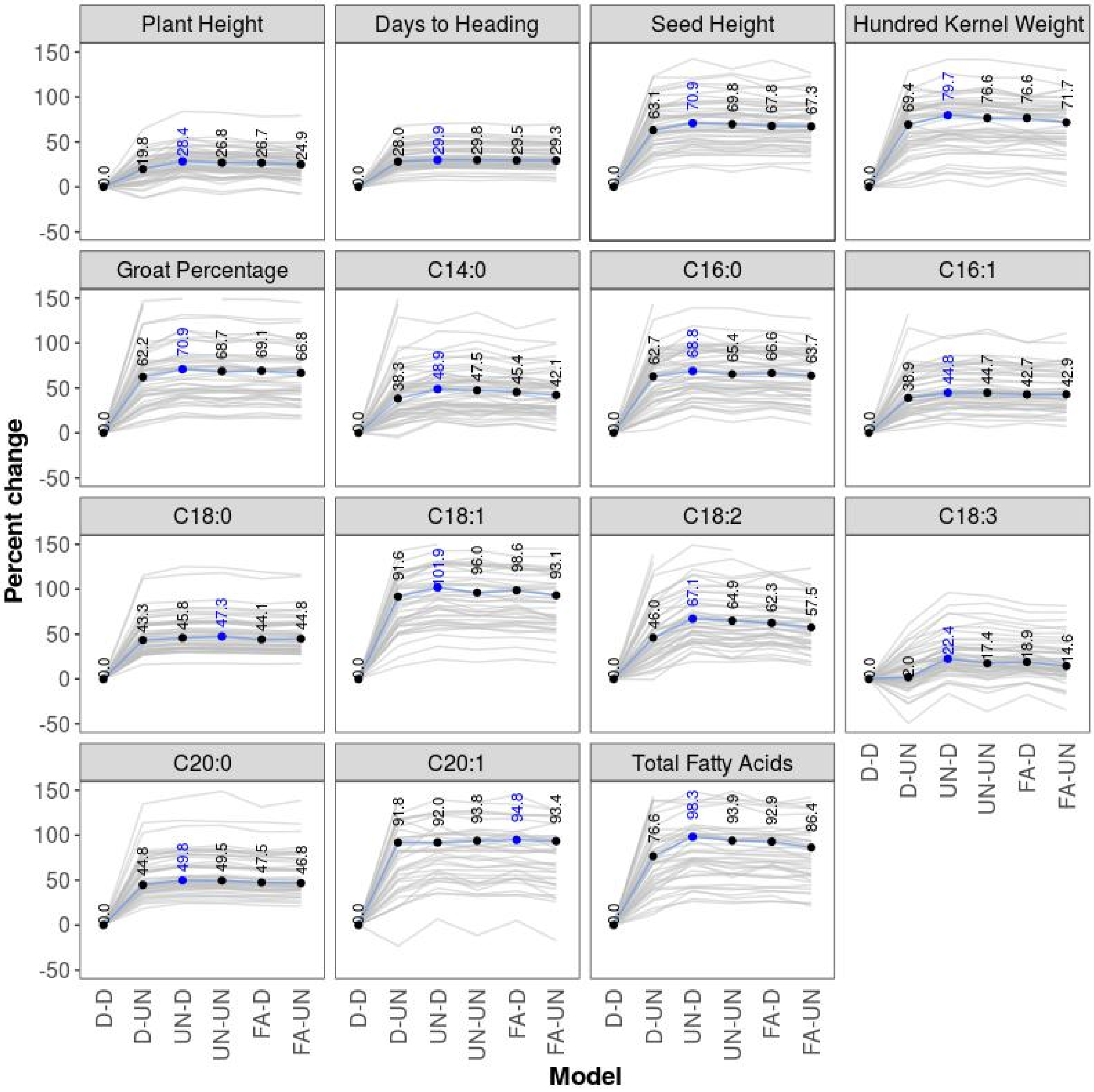


**Supplemental Figure 6.** Percentage changes in prediction accuracy estimated from multi-environment GBLUP models over single-environmental GBLUP for the 15 phenotypic traits in the Elite panel. Each gray line represents a re-sampling run, and the blue line represents median values across 50 re-sampling runs. The number in blue is the approach that showed the most improvement. For each model, the uppercase letters before and after the hyphen represent genetic and residual covariance structures: D=diagonal, UN=unstructured, FA=factor-analytic.

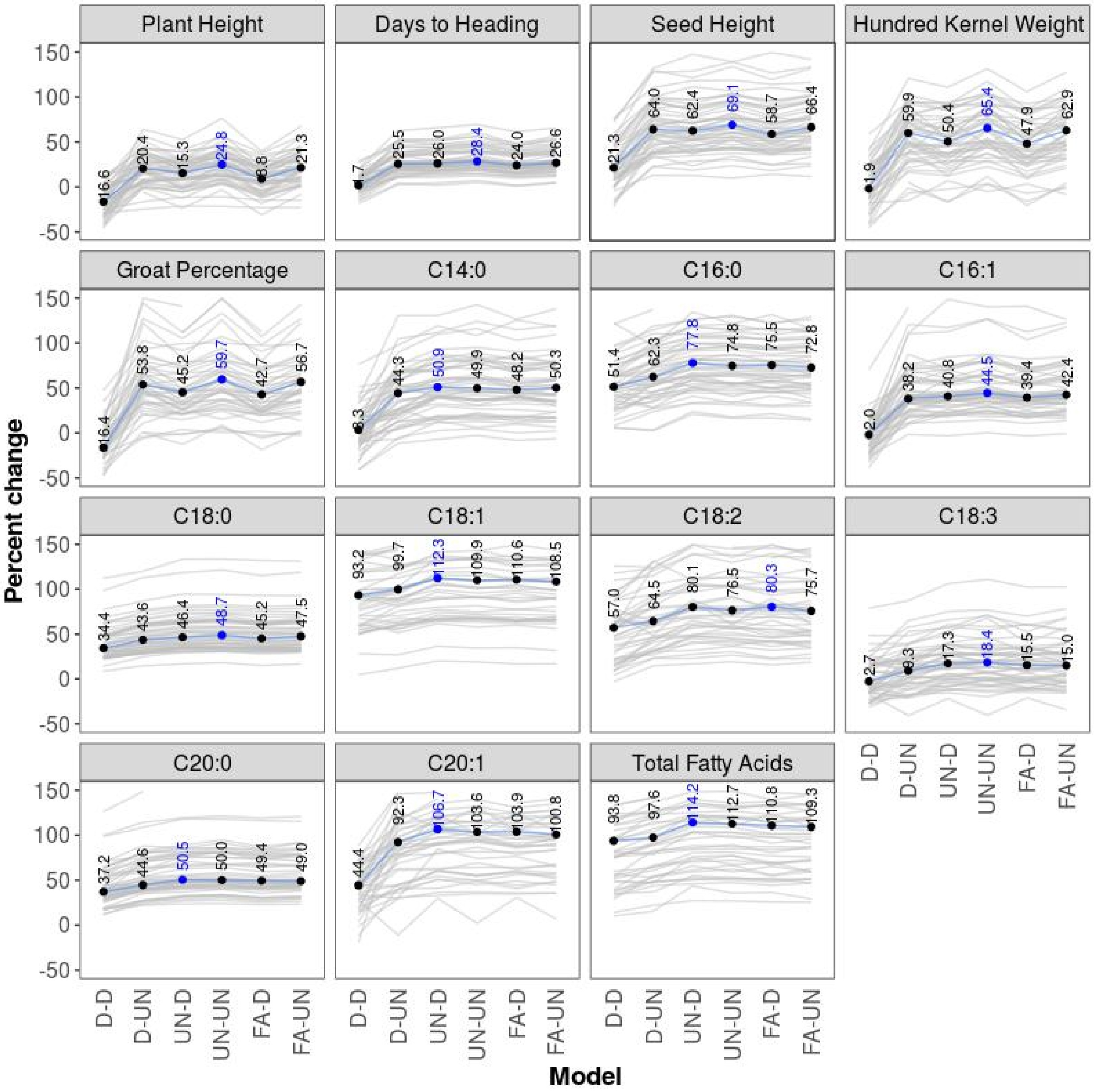


**Supplemental Figure 7.** Percentage changes in prediction accuracy estimated from multi-environment metabolite BLUP (M) models over single-environmental GBLUP for the 15 phenotypic traits in the Elite panel. Each gray line represents a re-sampling run, and the blue line represents median values across 50 re-sampling runs. The number in blue is the approach that showed the most improvement. For each model, the uppercase letters before and after the hyphen represent genetic and residual covariance structures: D=diagonal, UN=unstructured, FA=factor-analytic.

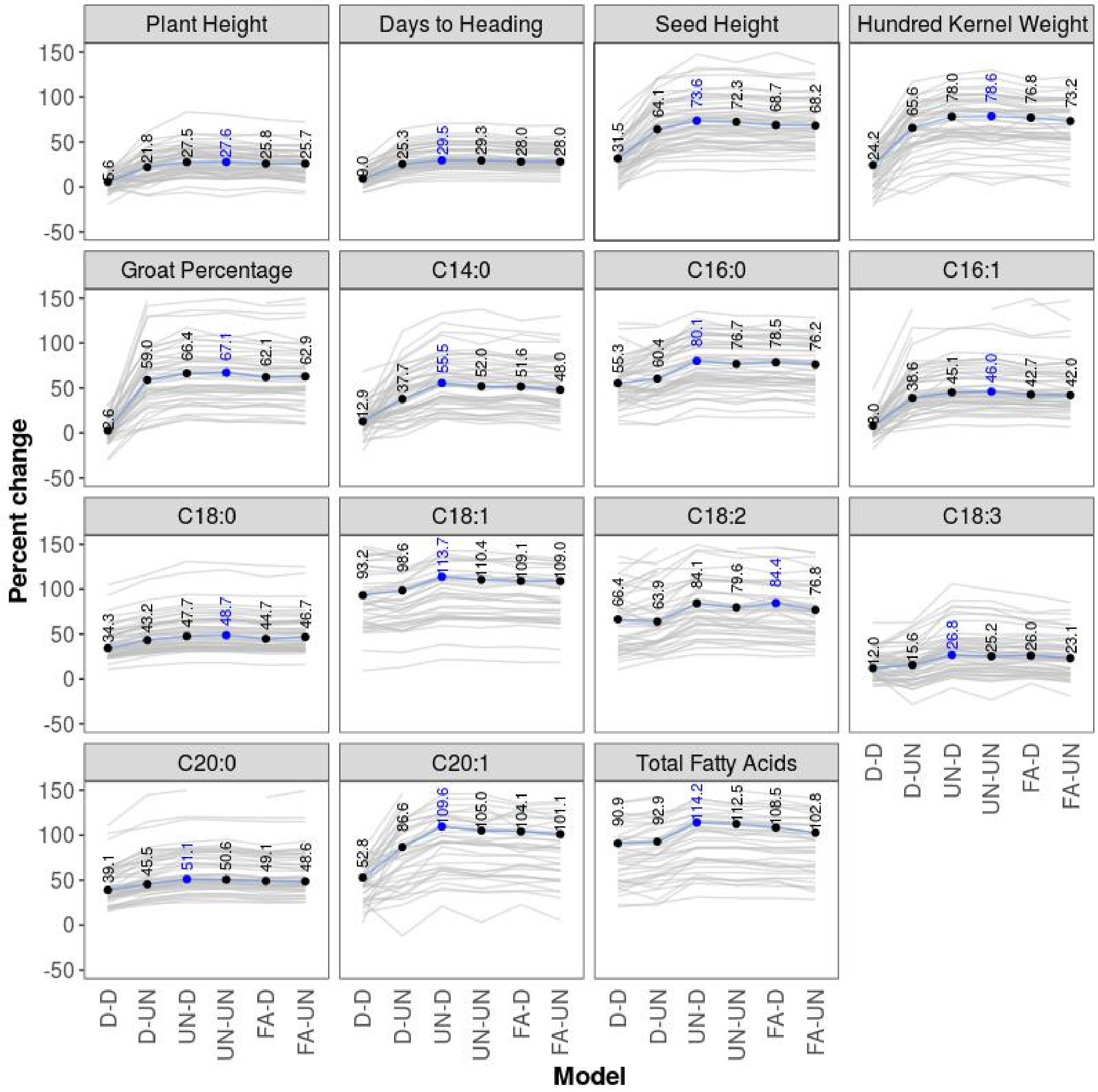


**Supplemental Figure 8.** Percentage changes in prediction accuracy estimated from multi-environment genomic and metabolomic BLUP (G+M) models over single-environmental GBLUP for the 15 phenotypic traits in the Elite panel. Each gray line represents a re-sampling run, and the blue line represents median values across 50 re-sampling runs. The number in blue is the approach that showed the most improvement. For each model, the uppercase letters before and after the hyphen represent genetic and residual covariance structures: D=diagonal, UN=unstructured, FA=factor-analytic.

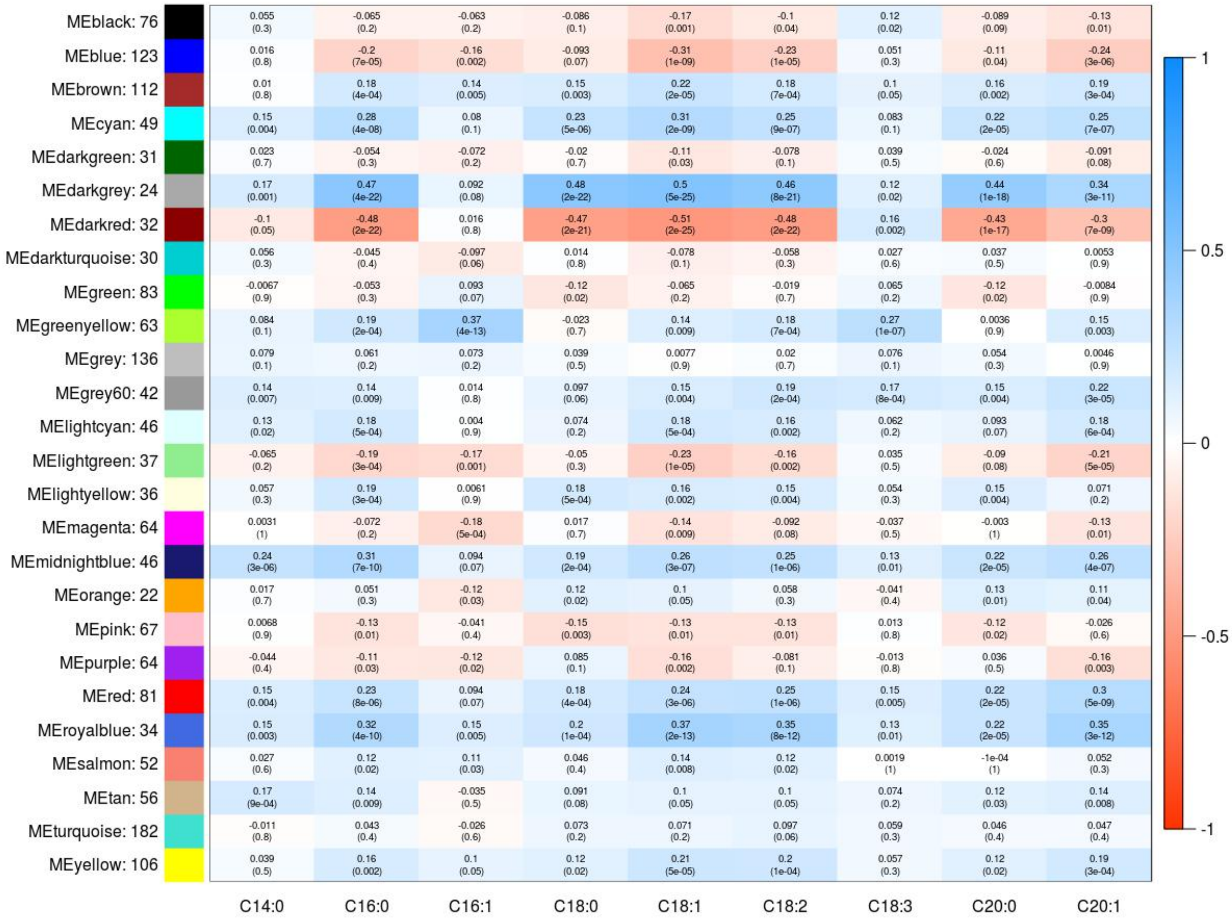


**Supplemental Figure 9.** Correlation between eigenvector of network modules and fatty acid traits. Each row corresponds to a module eigengene, column to a fatty acid trait. Each cell contains the corresponding correlation and p-value. The table is color-coded by correlation according to the color legend.

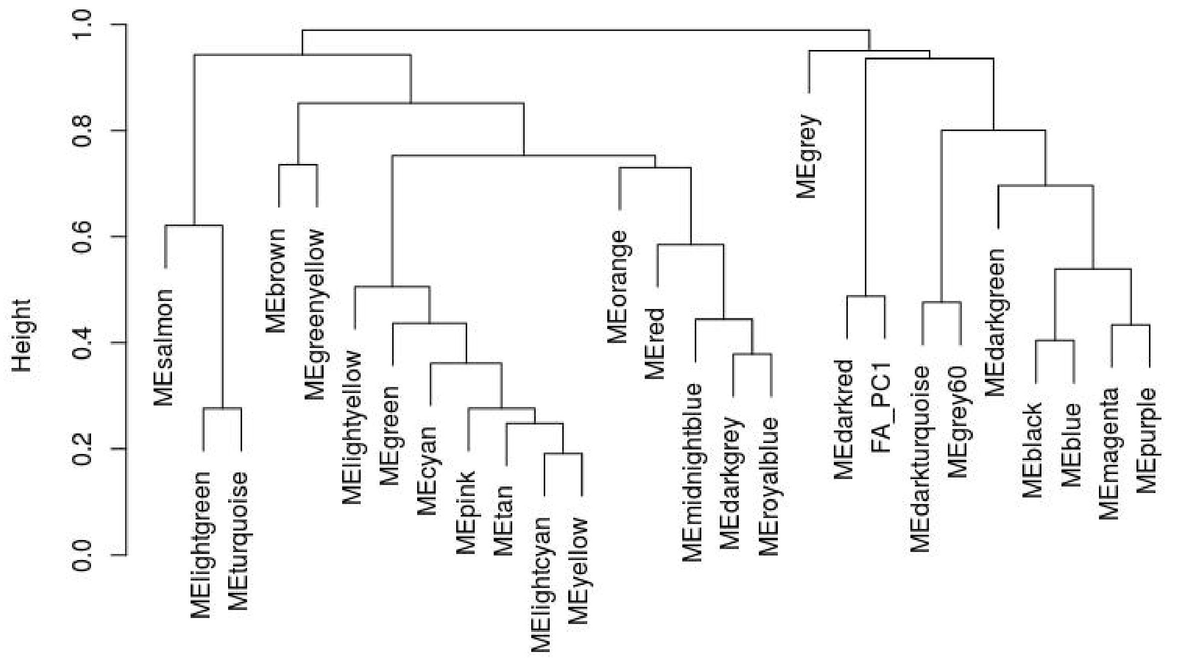


**Supplemental Figure 10.** Hierarchical clustering dendrogram of the network eigenvectors, PC1 and PC2 of the nine fatty acids.

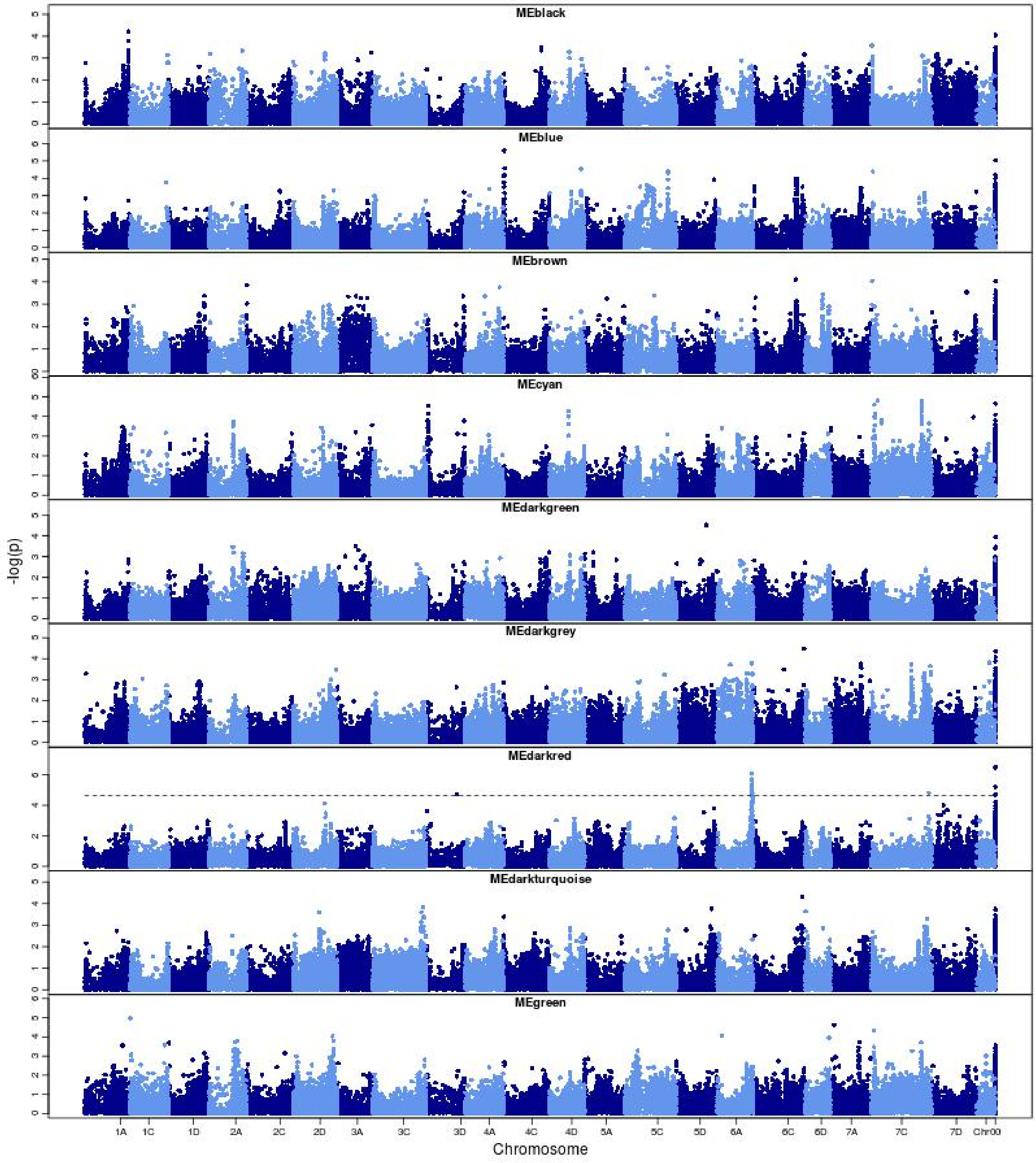


**Supplemental Figure 11.** Manhattan plots of eigenvectors of twenty-six network modules identified by WGCNA and PC1 of fatty acids in the Diversity panel. The dashed line corresponds to an FDR rate of 0.05 (continued on the next page).

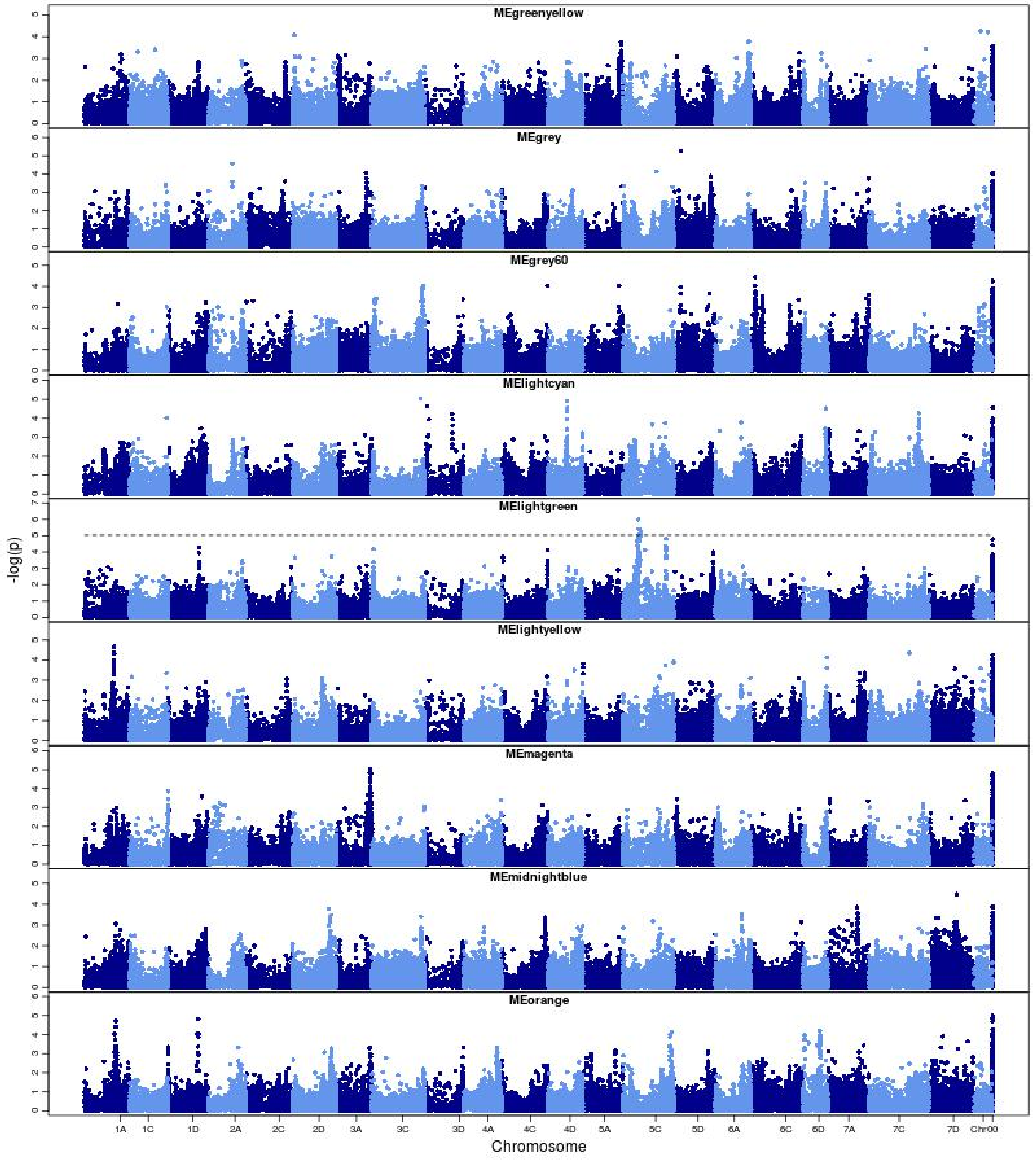


**Supplemental Figure 11.** Manhattan plots of eigenvectors of twenty-six network modules identified by WGCNA and PC1 of fatty acids in the Diversity panel. The dashed line corresponds to an FDR rate of 0.05 (continued on the next page).

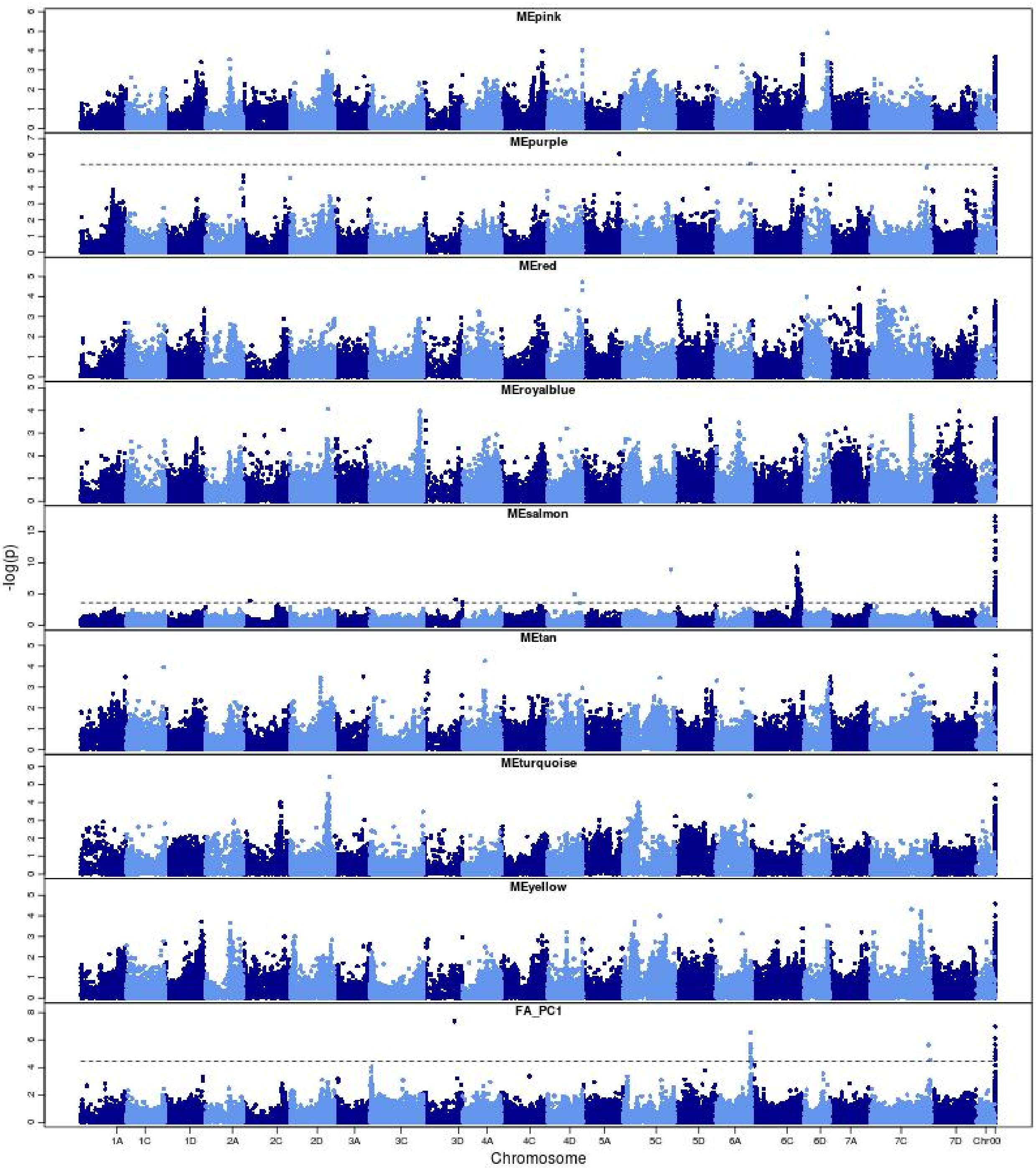


**Supplemental Figure 11.** Manhattan plots of eigenvectors of twenty-six network modules identified by WGCNA and PC1 of fatty acids in the Diversity panel. The dashed line corresponds to an FDR rate of 0.05.

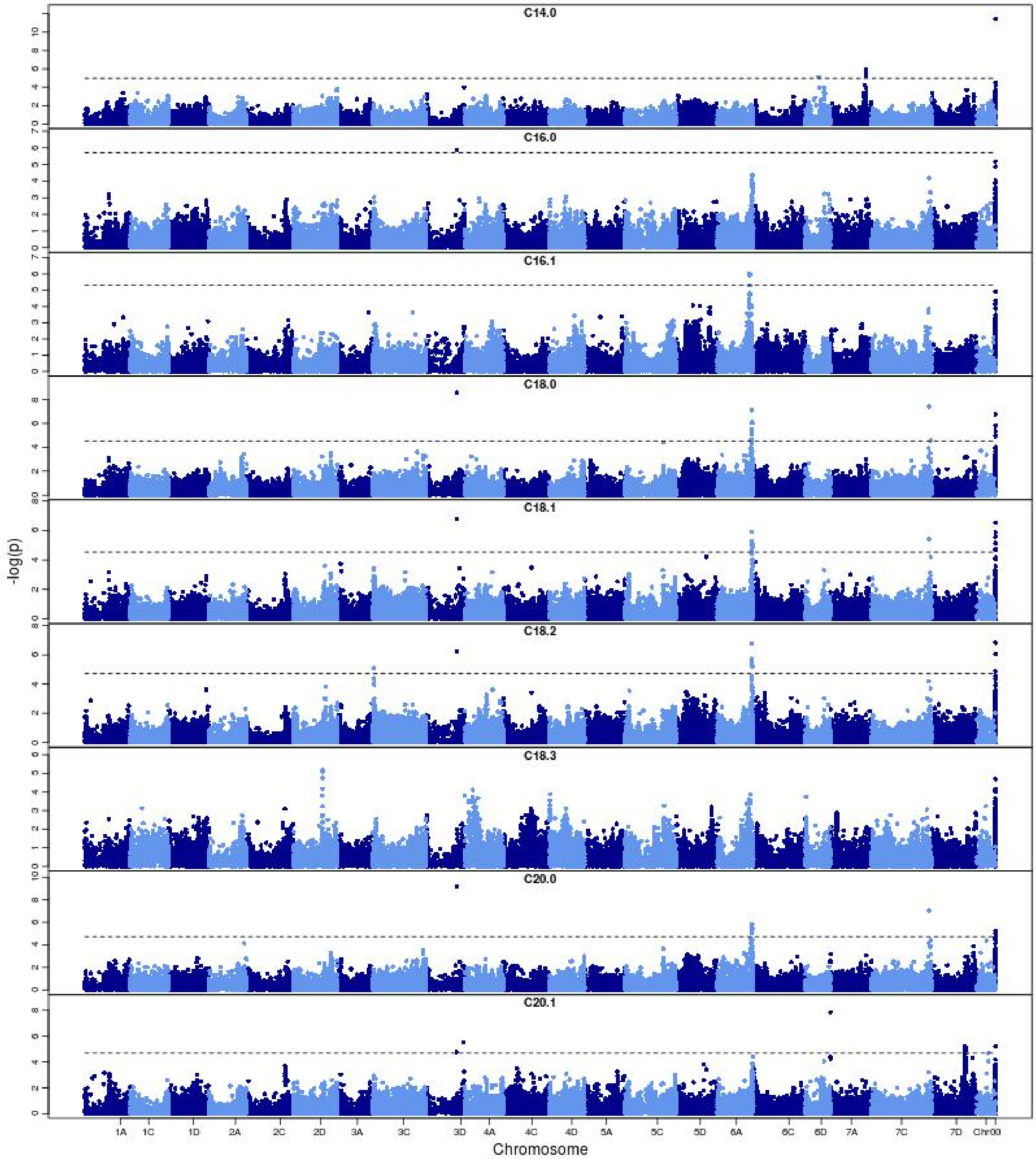


**Supplemental Figure 12.** Manhattan plots of fatty acids traits in the Diversity panel. The dashed line corresponds to an FDR rate of 0.05.

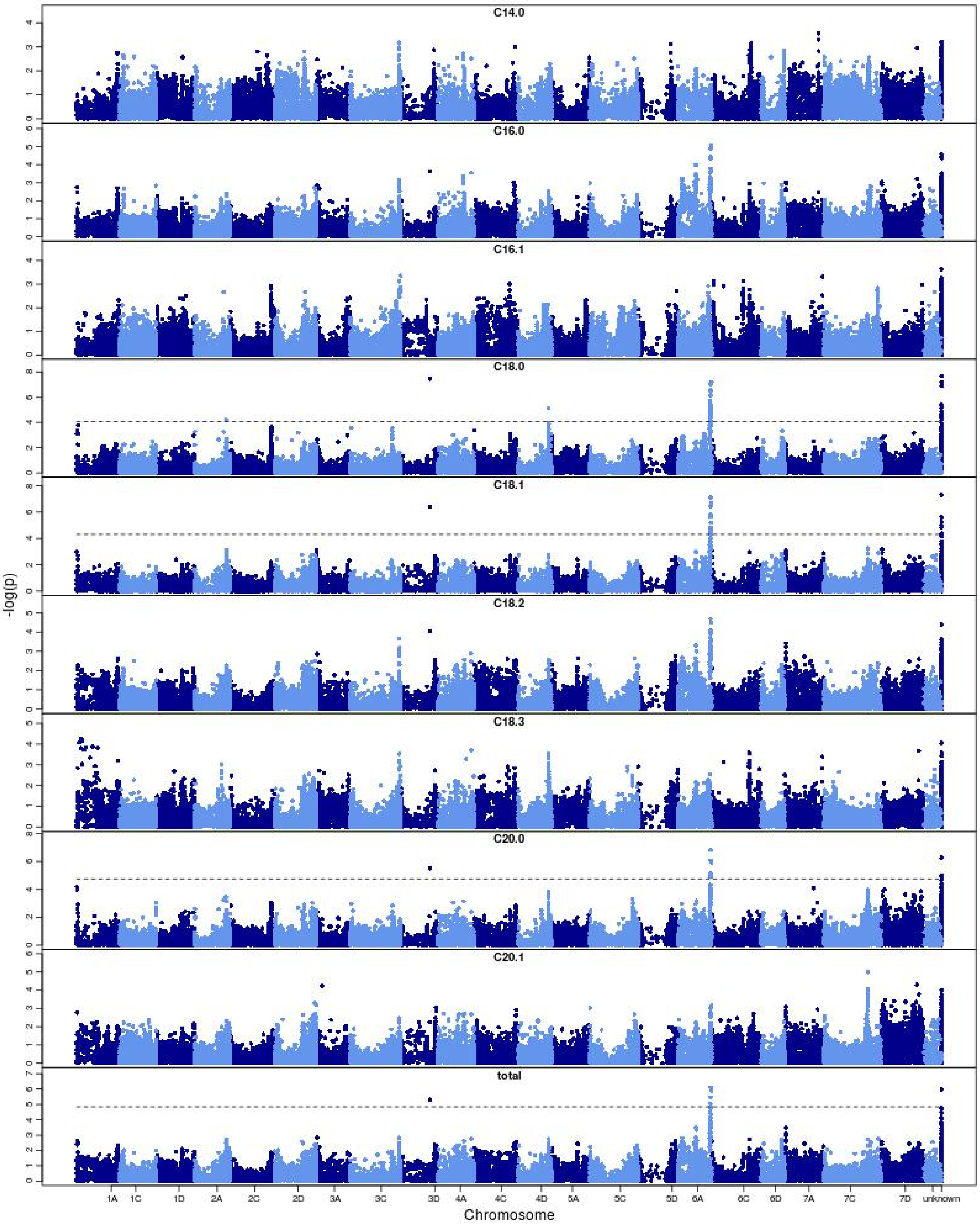


**Supplemental Figure 13.** Manhattan plots of fatty acids traits in the Elite panel. The fatty acids data was collected from filed trial located at Crookston, MN. The dashed line corresponds to an FDR rate of 0.05 (continued on the next page).

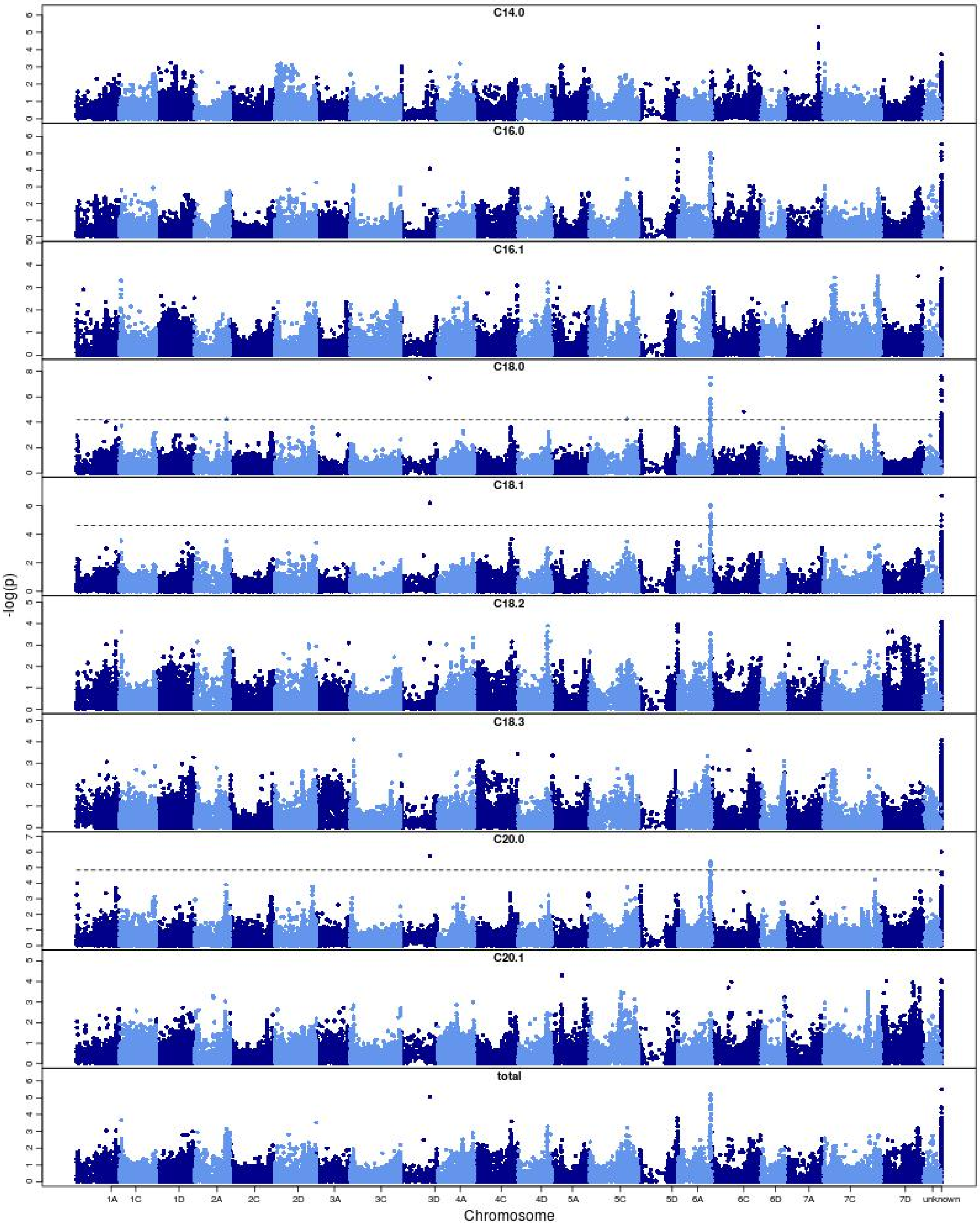


**Supplemental Figure 13.** Manhattan plots of fatty acids traits in the Elite panel. The fatty acids data was collected from filed trial located at Brookings, SD. The dashed line corresponds to an FDR rate of 0.05 (continued on the next page).

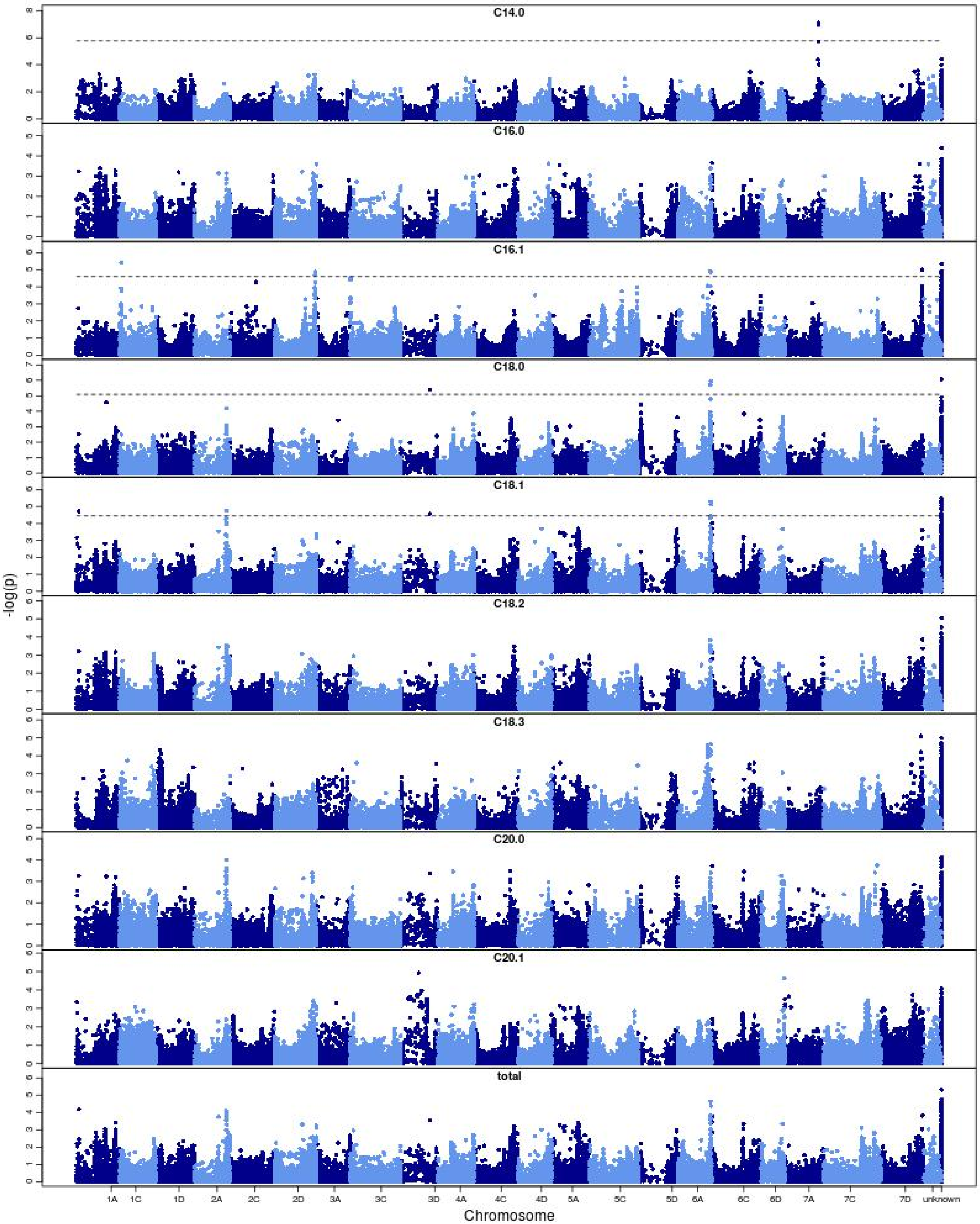


**Supplemental Figure 13.** Manhattan plots of fatty acids traits in the Elite panel. The fatty acids data was collected from  filed trial located at Madison, WI. The dashed line corresponds to an FDR rate of 0.05.

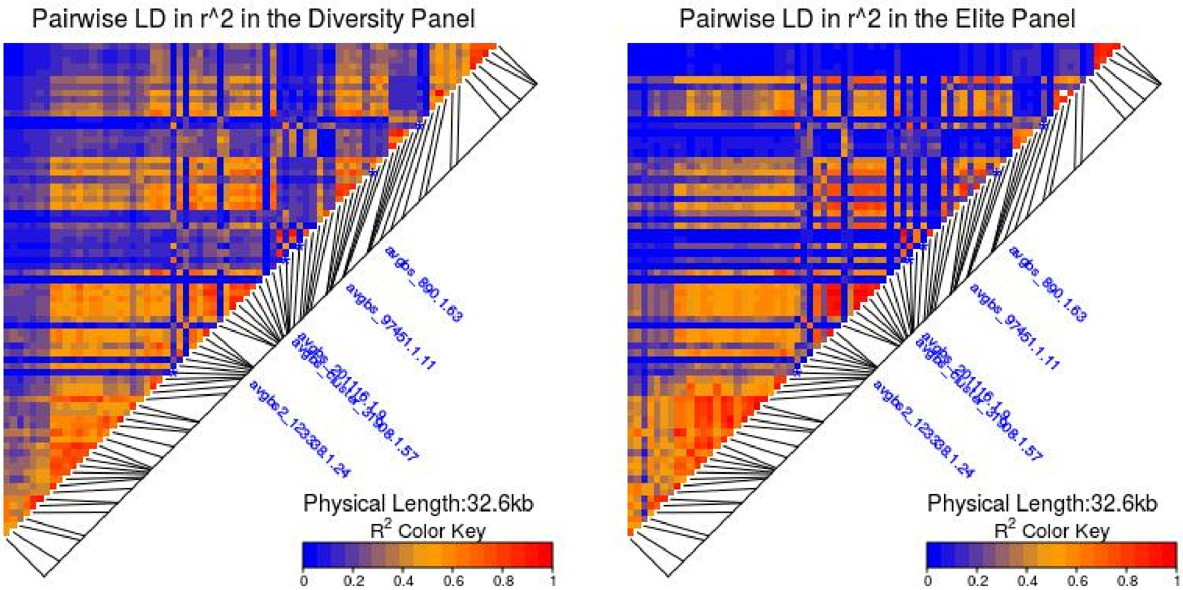


**Supplemental Figure 14.** LD heatmaps of significant markers of *QTL-6A* and the markers in LD with them in the Diversity and Elite panels. Only 75 SNPs on chromosome 6A with putative physical positions were used for the LD plot.The top 5 SNPs with highest marker score (-$\log_{10}$Pvalue) were labeled.

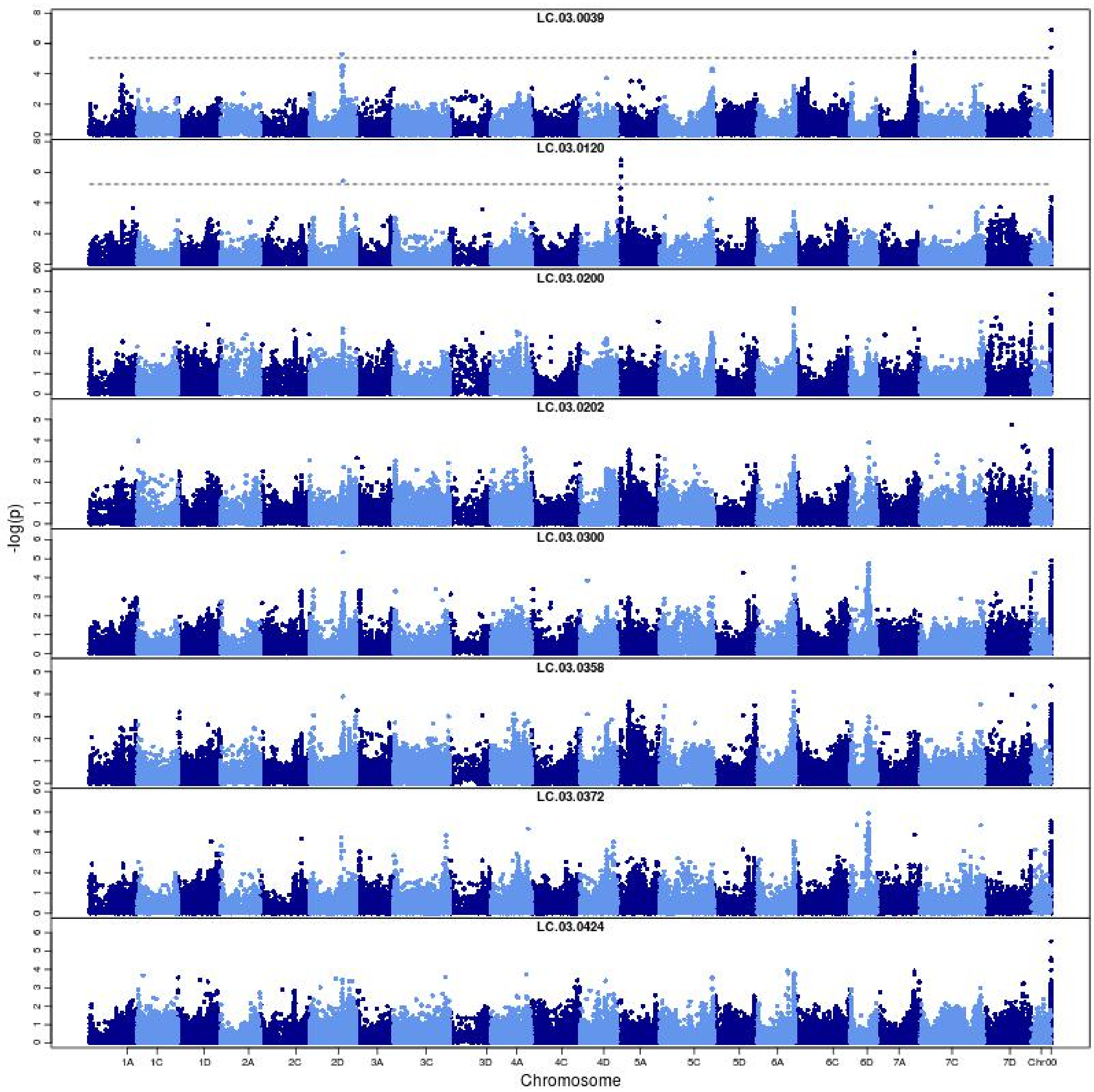


**Supplemental Figure 15.** Manhattan plots of metabolites in the darkred module identified by WGCNA in the Diversity panel. The dashed line corresponds to an FDR rate of 0.05 (continued on the next page).

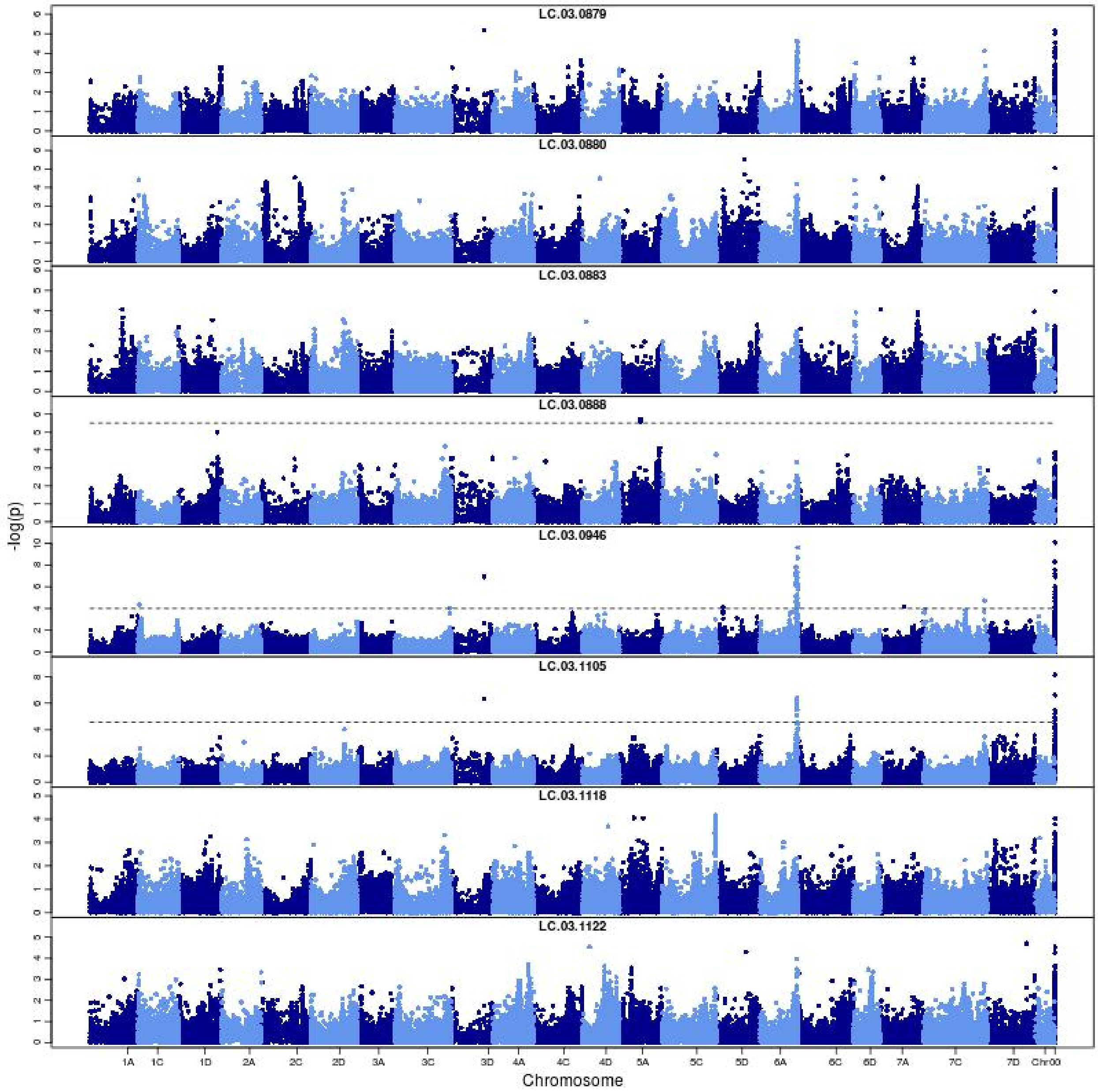


**Supplemental Figure 15.** Manhattan plots of metabolites in the darkred module identified by WGCNA in the Diversity panel. The dashed line corresponds to an FDR rate of 0.05 (continued on the next page).

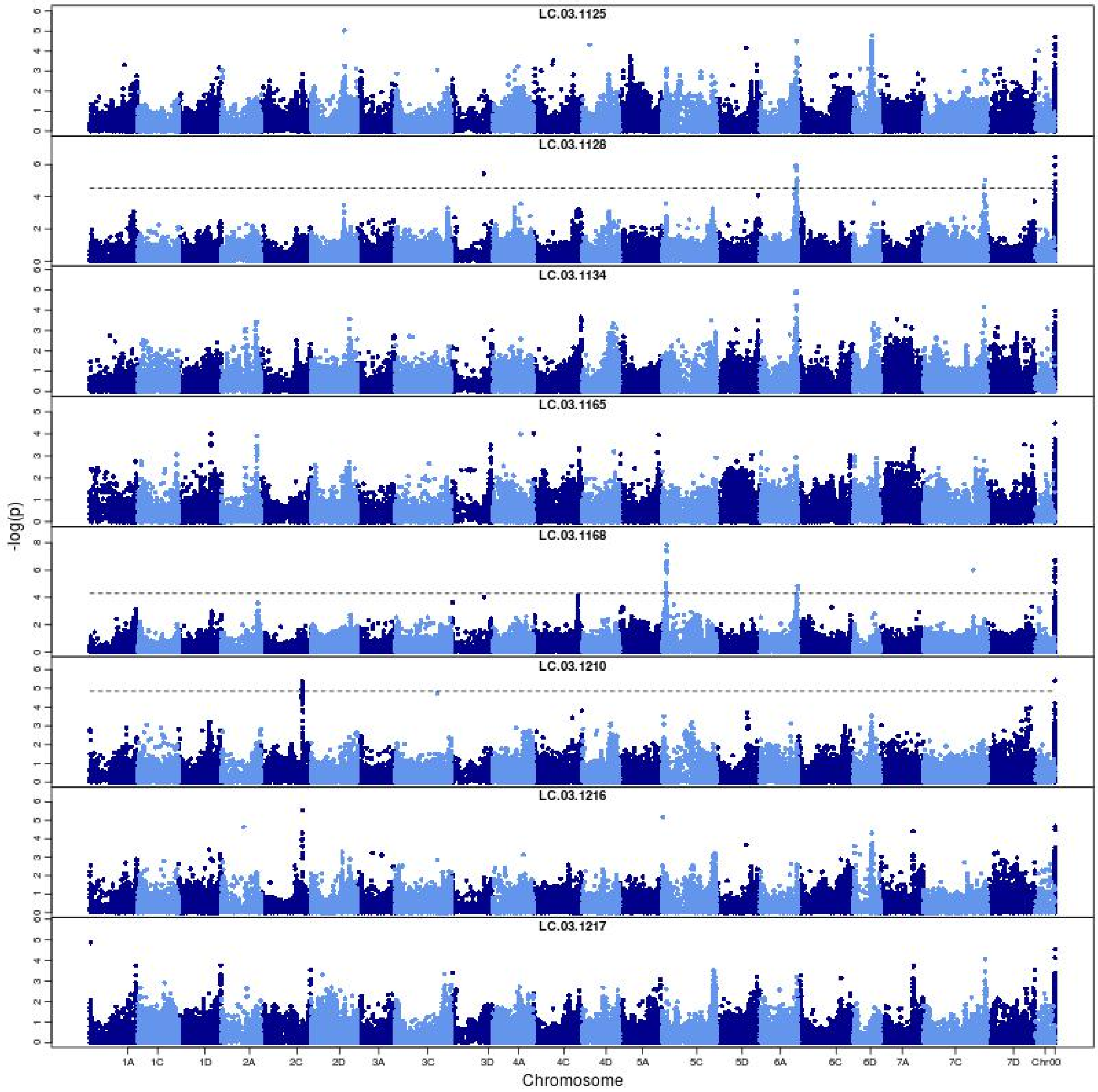


**Supplemental Figure 15.** Manhattan plots of metabolites in the darkred module identified by WGCNA in the Diversity panel. The dashed line corresponds to an FDR rate of 0.05 (continued on the next page).

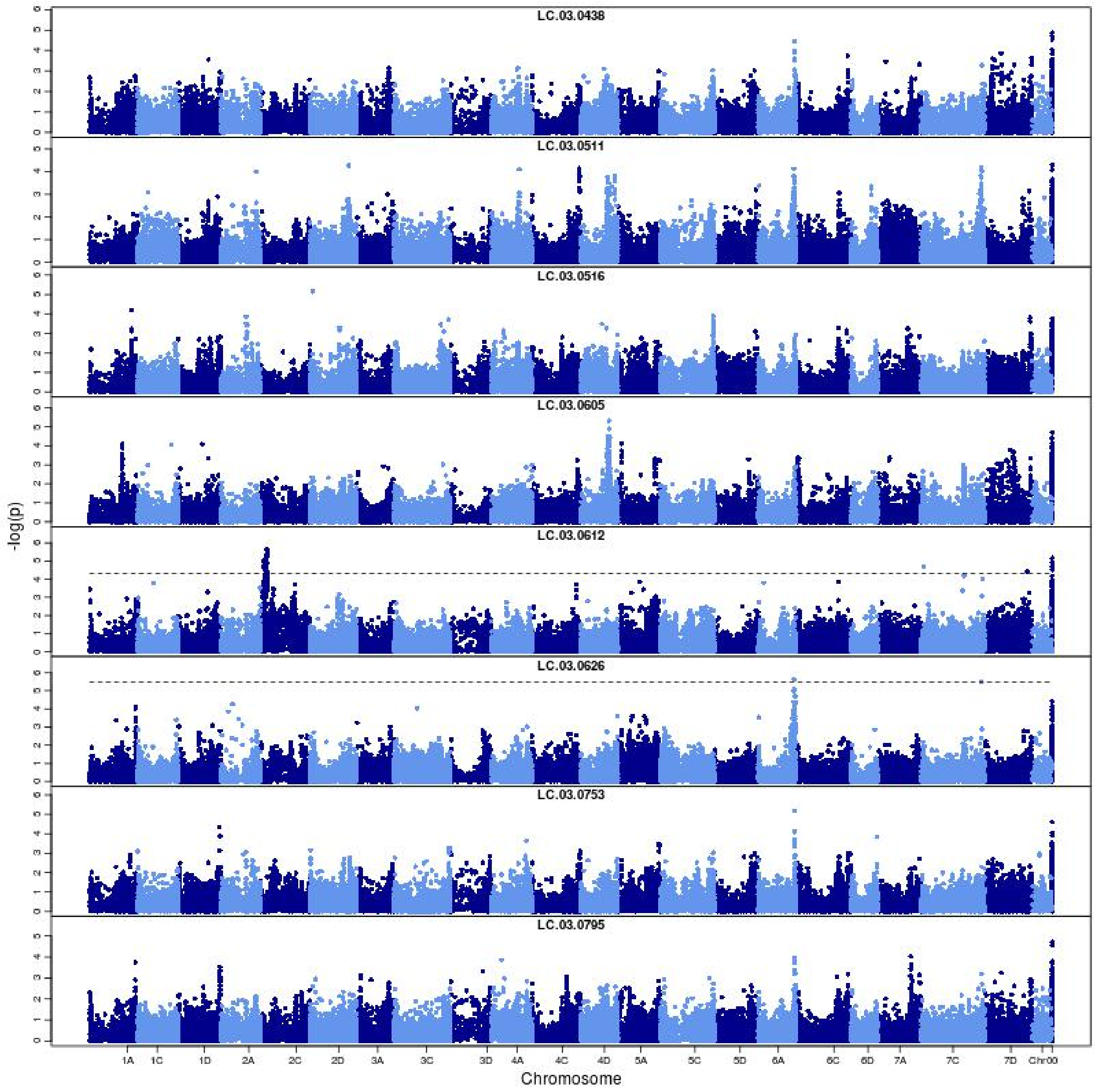


**Supplemental Figure 15.** Manhattan plots of metabolites in the darkred module identified by WGCNA in the Diversity panel. The dashed line corresponds to an FDR rate of 0.05.

#### Supplemental Tables

**Supplemental Table 1** Phenotypic traits evaluated in the Diversity panel and the Elite panel and their heritabilities

**Supplemental Table 2** A list of the 368 and 232 lines included in the Diversity panel and Elite panel

**Supplemental Table 3.** Percent changes in prediction accuracy over GBLUP of 5, 10, 11, 17 and 17 traits from transcriptomic BLUP (T), metabolomic BLUP (M), G+T, G+M and G+T+M models in the Diversity panel.

**Supplemental Table 4** Annotation of Metabolites identified in the Diversity Panel

**Supplemental Table 5** Lipids and lipid-like molecules enrichment test in 26 network modules identified by WGCNA based on metabolite annotation

**Supplemental Table 6.** Percent changes in prediction accuracy of G+M over GBLUP(G) and metabolomic BLUP (M) models for the 17 traits in the Diversity panel.

**Supplemental Table 7** Description of the multi-trait models

**Supplemental Table 1** Phenotypic traits evaluated in the Diversity panel and the Elite panel and their heritabilities

| | | Diversity Panel | Elite Panel | | | |
|---|---|---|---|---|---|---|
| Category | Trait Unknownme | ITH | Across-Site | MN | SD | WI |
| Agronomic | Plant Height | 0.82 | 0.76 | 0.63 | 0.50 | 0.52 |
| | Days to Heading | 0.80 | 0.89 | 0.97 | 0.60 | 0.71 |
| | Seed Length | 0.79 | 0.81 | 0.81 | – | 0.62 |
| | Seed Width | 0.86 | 0.80 | 0.87 | – | 0.73 |
| | Seed Height | 0.89 | 0.78 | 0.76 | 0.82 | 0.69 |
| | Hundred Kernel Weight | 0.89 | 0.67 | 0.64 | 0.65 | 0.56 |
| | Hundred Hull Weight | 0.82 | 0.69 | 0.51 | 0.41 | – |
| | Groat Percentage | 0.88 | 0.77 | 0.84 | 0.90 | 0.46 |
| Fatty Acids | C14:0 | 0.80 | 0.60 | 0.67 | 0.82 | 0.75 |
| | C16:0 | 0.80 | 0.71 | 0.62 | 0.80 | 0.76 |
| | C16:1 | 0.75 | 0.78 | 0.67 | 0.80 | 0.72 |
| | C18:0 | 0.92 | 0.89 | 0.89 | 0.93 | 0.92 |
| | C18:1 | 0.89 | 0.80 | 0.66 | 0.81 | 0.82 |
| | C18:2 | 0.78 | 0.62 | 0.54 | 0.74 | 0.71 |
| | C18:3 | 0.49 | 0.58 | 0.67 | 0.68 | 0.73 |
| | C20:0 | 0.87 | 0.89 | 0.77 | 0.85 | 0.89 |
| | C20:1 | 0.74 | 0.71 | 0.55 | 0.58 | 0.76 |
| | Total Fatty Acids | – | 0.70 | 0.58 | 0.78 | 0.76 |

ITH=Ithaca, MN=Minnesota, SD=South Dakota, WI=Wisconsin

**Supplemental Table 2** A list of the 368 and 232 lines included in the Diversity panel and Elite panel

| Line Name | Population |
|---|---|
| IL09-5508 | Elite |
| AAC_ROSKENS | Both |
| ND102000 | Elite |
| WIX8787-3 | Elite |
| IL06-3761 | Elite |
| AAC_NICOLAS | Elite |
| IL11-5748 | Elite |
| P973A38-9-3-2-29 | Elite |
| OTEE | Elite |
| RON | Elite |
| MN04242 | Elite |
| CLINTFORD | Both |
| SD120129 | Elite |
| MN09115 | Elite |
| SD050834 | Elite |
| SD120640 | Elite |
| BETAGENE | Both |
| SD110304 | Elite |
| IA02130-2-2 | Elite |
| SD110765 | Elite |
| SD041445-119 | Elite |
| P0541A1-1 | Elite |
| SD080348 | Elite |
| PI344841 | Both |
| P0714A1-29-2 | Elite |
| WIX9500-6 | Elite |
| SD091510 | Elite |
| SD120258 | Elite |
| SD120524 | Elite |
| MN08243 | Both |
| SD120266 | Elite |
| IL09-6937 | Elite |

| Line Name | Population |
|---|---|
| SD110808 | Elite |
| SD120456 | Elite |
| IL04-7077 | Both |
| CORRAL | Both |
| ND060182 | Both |
| SD020883 | Elite |
| WIX9878-3 | Elite |
| MN06203 | Elite |
| SD081644 | Elite |
| MN08252 | Both |
| BADGER | Elite |
| LEGGETT | Elite |
| P0528A1-6-1 | Elite |
| MN10209 | Elite |
| AAC_OAKLIN | Elite |
| SD110605 | Elite |
| P021A1-66-2 | Elite |
| P0528A1-1-1 | Elite |
| WIX9528-1 | Elite |
| ND030349 | Elite |
| SD081107 | Elite |
| SD100184 | Elite |
| SD090880 | Elite |
| KAME | Elite |
| FL0238BSB-22 | Both |
| MN08160 | Both |
| SD090893 | Elite |
| MN07208 | Elite |
| IL11-2353 | Elite |
| IL08-2010 | Elite |
| WPAT04P03-PY3B | Elite |
| IA111003 | Both |
| AAC_ALMONTE | Elite |
| IL00-1030 | Elite |

| Line Name | Population |
|---|---|
| OT2083 | Elite |
| SD050938 | Elite |
| P0216A1-1 | Elite |
| ND050490 | Elite |
| SD041405 | Elite |
| P021A1-25 | Elite |
| HORSEPOWER | Elite |
| IA02130-2-3 | Elite |
| WIX8787-4 | Elite |
| MNBT1021-1 | Elite |
| AC_ASSINIBOIA | Elite |
| MN02234 | Elite |
| SD050608 | Elite |
| CLINTLAND64 | Elite |
| OA1342-2 | Elite |
| SD041016 | Elite |
| OA1256-1 | Elite |
| SD120553 | Elite |
| SD090965 | Elite |
| MN04136 | Elite |
| TX02U7479 | Both |
| IA02168-1-1 | Elite |
| SD60130 | Elite |
| IL02-8011 | Both |
| LAO-1012-040 | Elite |
| MN10130 | Elite |
| ND090868 | Elite |
| WIX9287-2 | Elite |
| MN04120 | Elite |
| COLT | Elite |
| ND051306 | Elite |
| IL08-9201 | Elite |
| QUALITY\|PI289587 | Both |
| MN06120 | Elite |

| Line Name | Population |
|---|---|
| SABER | Elite |
| WIX10088-6 | Elite |
| MN03114 | Elite |
| ND021612 | Elite |
| MN10253 | Elite |
| X9221-8 | Both |
| OA1341-1 | Elite |
| HA07-02X22-3 | Elite |
| P075A1-7 | Elite |
| OA1357-2 | Elite |
| MN08138 | Elite |
| SD111922 | Elite |
| SD041451 | Elite |
| SD100719 | Elite |
| EXCEL | Both |
| IL05-7133 | Elite |
| SD050945 | Elite |
| SD090510 | Elite |
| SD090552 | Elite |
| ND111357 | Elite |
| WIX9449-1 | Elite |
| P9741A1-4-6-86 | Elite |
| IL05-9931 | Elite |
| SD030888 | Elite |
| ND060342 | Both |
| MN09255 | Elite |
| ND070388 | Elite |
| SD090780 | Elite |
| SD081949 | Elite |
| MN05237 | Elite |
| SD081577 | Elite |
| IA02010-2-1 | Elite |
| WIX10045-9 | Elite |
| SD100198 | Elite |

| Line Name | Population |
|---|---|
| SD082192 | Elite |
| MN09230 | Elite |
| SD081085 | Elite |
| ND040492 | Elite |
| MN11221 | Elite |
| OT2074 | Elite |
| SD100602 | Elite |
| MN11110 | Elite |
| OA1331-6 | Elite |
| WIX9487-1 | Elite |
| ND040250 | Elite |
| SD031128 | Elite |
| OA1271-3 | Elite |
| UnknownTTY | Elite |
| WIX10045-12 | Elite |
| MN05119 | Elite |
| LAO-1104-028C1 | Elite |
| SD080788 | Elite |
| 95AB12770 | Both |
| SD61081 | Elite |
| ND040196 | Elite |
| MN09223 | Elite |
| SD050716 | Elite |
| HA08-03X15-1 | Elite |
| IL05-3337 | Elite |
| SD081563 | Elite |
| MN06213 | Elite |
| P075A1-3-2 | Elite |
| WIX9645-1 | Elite |
| MN08260 | Both |
| WIX9414-1 | Elite |
| P021A1-78-1 | Elite |
| 04P07A-BS5C | Elite |
| SD110640 | Elite |

| Line Name | Population |
|---|---|
| ND090807 | Elite |
| GOPHER | Elite |
| SD100015 | Elite |
| MN05157 | Elite |
| SUMO | Elite |
| SD120316 | Elite |
| WIX9562-5 | Elite |
| MN05205 | Elite |
| WIX9897-5 | Elite |
| 06P04A-E07B3 | Elite |
| P0216A1-1-45 | Elite |
| MN09105 | Elite |
| SD100454 | Elite |
| ND021052 | Elite |
| SD110470 | Elite |
| IL06-8153 | Elite |
| MN10121 | Elite |
| TX07CS1402 | Both |
| SD050616 | Elite |
| IA01005-2 | Elite |
| ND100362 | Elite |
| SD081038 | Elite |
| MN09103 | Elite |
| DON | Elite |
| MN09256 | Elite |
| IL09-5239 | Elite |
| MN07204 | Elite |
| IL09-5745 | Elite |
| OMSKIJ19260 | Elite |
| SHELBY427 | Elite |
| SD61433 | Elite |
| SD051005 | Elite |
| SD101027 | Elite |
| SD080611 | Elite |

| Line Name | Population |
|---|---|
| IL06-5465 | Elite |
| OPTIMUM | Both |
| SD100644 | Elite |
| WIX9150-1 | Elite |
| ND101473 | Elite |
| HAYDEN | Elite |
| IL02-8663 | Elite |
| 00P28-AN01B1 | Elite |
| CIAV5218 | Both |
| SD090240 | Elite |
| MORTON | Elite |
| MN02231 | Elite |
| WIX8791-1 | Elite |
| 04P07B-GN1C | Elite |
| ND051312 | Both |
| WIX9082-1 | Both |
| SD60980 | Elite |
| MN08211 | Elite |
| SD111736 | Elite |
| MN04232 | Elite |
| ND070182 | Elite |
| MN08130 | Both |
| SHERWOOD | Both |
| IL05-8515 | Both |
| PERDEBERG | Both |
| ANDREW | Both |
| X8826-1 | Both |
| WOODBURN | Both |
| DEON | Elite |
| OGLE | Elite |
| 9876C1-2-1-5-2-4-1 | Diversity |
| 0219A1-84-4-4-4-4 | Diversity |
| IL05-10069 | Diversity |
| WHITE_TARTARIAN\|CIAV800 | Diversity |

| Line Name | Population |
|---|---|
| IA111188 | Diversity |
| X9410-2 | Diversity |
| GERE | Diversity |
| 02HO-139 | Diversity |
| OA1232-2 | Diversity |
| MN861900 | Diversity |
| LANG-DOERFLERS_WEIHENSTEPHANER_WEISSHAFER\|PI180932 | Diversity |
| RODGERS | Diversity |
| RANCH | Diversity |
| ND060223 | Diversity |
| X9287-2 | Diversity |
| CIAV5389 | Diversity |
| X9507-1 | Diversity |
| IL05-1705 | Diversity |
| IA111041 | Diversity |
| 027A1-87-8-1 | Diversity |
| OA1250-2 | Diversity |
| PI577862 | Diversity |
| OA1226-4 | Diversity |
| LAO-1135-015 | Diversity |
| IA111006 | Diversity |
| IA111186 | Diversity |
| IA111153 | Diversity |
| DRUMMOND | Diversity |
| SA070906 | Diversity |
| 04P07B-GY5E | Diversity |
| OA1248-1 | Diversity |
| NIAGARA | Diversity |
| IA111235 | Diversity |
| OA1197-1 | Diversity |
| PI159180 | Diversity |
| CEUnknownD88\|PI361889 | Diversity |
| IA111281 | Diversity |
| ABERDEEN_SELECTION1939-2005 | Diversity |

| Line Name | Population |
|---|---|
| TIOGA | Diversity |
| IA111199 | Diversity |
| ENDRESS | Diversity |
| FLORIDA500 | Diversity |
| ONOHOJSKIJ_A-547 | Diversity |
| SA070781 | Diversity |
| X8903-2 | Diversity |
| IA111036 | Diversity |
| RANSOM | Diversity |
| IA111180 | Diversity |
| MN08268 | Diversity |
| CIAV6209 | Diversity |
| CIAV5220 | Diversity |
| ND060111 | Diversity |
| IA111136 | Diversity |
| IA111045 | Diversity |
| CDC_BIG_BROWN | Diversity |
| SA071616 | Diversity |
| IA111027 | Diversity |
| ND061813 | Diversity |
| MN06125 | Diversity |
| 00P06-HD1D | Diversity |
| IA111119 | Diversity |
| ND061614 | Diversity |
| PI168094 | Diversity |
| 059A1-2-2-4 | Diversity |
| HA05AB20-1 | Diversity |
| FLAMINGSKOMET | Diversity |
| 02G31-NL7A | Diversity |
| CD3774 | Diversity |
| II-30-39 | Diversity |
| SA061148 | Diversity |
| AVOINE_NUE-NUE_NOISE\|PI401772 | Diversity |
| CIAV5222 | Diversity |

| Line Name | Population |
| --- | --- |
| SA071760 | Diversity |
| IA111071 | Diversity |
| LAO-1136-014 | Diversity |
| HAZEL | Diversity |
| KHARKOVSKIJ596\|PI158223 | Diversity |
| BANKUT_GALBEN | Diversity |
| TX07CS1039 | Diversity |
| PI577985 | Diversity |
| OA1263-2 | Diversity |
| KOLBU | Diversity |
| TX07CS2140 | Diversity |
| MN861218 | Diversity |
| PENNCOMP38 | Diversity |
| HORIZON270 | Diversity |
| SYLVA | Diversity |
| GRANE | Diversity |
| OA1234-1 | Diversity |
| FLORILAND | Diversity |
| CIAV5666 | Diversity |
| TX05CS542 | Diversity |
| MN08234 | Diversity |
| IA111228 | Diversity |
| HOHENHEIMER_V\|PI180928 | Diversity |
| 97AB7767 | Diversity |
| IA111158 | Diversity |
| ND051236 | Diversity |
| IA111178 | Diversity |
| IA111123 | Diversity |
| SA070655 | Diversity |
| KRYMSKIJ90\|PI296174 | Diversity |
| 26-35-B69 | Diversity |
| Y-90 | Diversity |
| JERRY | Diversity |
| CDC_WEAVER | Diversity |

| Line Name | Population |
|---|---|
| HORIZON201 | Diversity |
| IL05-9948 | Diversity |
| TYLER | Diversity |
| ORBIT | Diversity |
| IA111018 | Diversity |
| CIAV5928 | Diversity |
| SELECTION3841-1 | Diversity |
| GOLDEN_RUSTPROOF\|CIAV1751 | Diversity |
| CDC_SOL-FI | Diversity |
| X345-1-B4-20-1 | Diversity |
| SEVERIANIN\|PI326231 | Diversity |
| IA111218 | Diversity |
| CILLA | Diversity |
| IA111044 | Diversity |
| OA1266-1 | Diversity |
| TIFT | Diversity |
| FL0047-J9 | Diversity |
| X9396-1 | Diversity |
| BIRI | Diversity |
| CHLUMECKY | Diversity |
| IA111282 | Diversity |
| STANTON\|PI412928 | Diversity |
| 00P01-A11A4 | Diversity |
| CDC_MINSTREL | Diversity |
| CIAV6130 | Diversity |
| IA111046 | Diversity |
| X9195-2 | Diversity |
| IA00020-12-3 | Diversity |
| TENNESSEE_SELECTION090 | Diversity |
| IA111185 | Diversity |
| IL03-7936 | Diversity |
| BRANCH | Diversity |
| LA02012-S-B-139-S2-B-S2-B-S2 | Diversity |
| PENNLINE6571 | Diversity |

| Line Name | Population |
|---|---|
| MN08270 | Diversity |
| IL06-3751 | Diversity |
| LA604 | Diversity |
| IA111161 | Diversity |
| IL06-3258 | Diversity |
| SA070469 | Diversity |
| 151C-2 | Diversity |
| OT380 | Diversity |
| IA111004 | Diversity |
| MN08230 | Diversity |
| WIR4301 | Diversity |
| LUTZ | Diversity |
| HUSAR | Diversity |
| MN08238 | Diversity |
| DAWSON | Diversity |
| PA7967-11759 | Diversity |
| X9421-3 | Diversity |
| IA111200 | Diversity |
| BIHARIA | Diversity |
| IA111177 | Diversity |
| IA111232 | Diversity |
| IL04-2727 | Diversity |
| MN08132 | Diversity |
| IL05-9330 | Diversity |
| IL86-4189 | Diversity |
| IL05-3806 | Diversity |
| WIR4672 | Diversity |
| MARION_QC | Diversity |
| IL75-5743 | Diversity |
| SA060605 | Diversity |
| MAIDA | Diversity |
| LAO-882-036 | Diversity |
| SA070592 | Diversity |
| CIAV6218 | Diversity |

| Line Name | Population |
|---|---|
| SA071405 | Diversity |
| IA111208 | Diversity |
| IA111244 | Diversity |
| Y-498 | Diversity |
| MOHAWK | Diversity |
| 04P06A-CY5D | Diversity |
| KAPP | Diversity |
| SA060830 | Diversity |
| X9290-2 | Diversity |
| BLANCHE_DE_HONGRIE | Diversity |
| IA111072 | Diversity |
| LENROC | Diversity |
| VI56 | Diversity |
| SA070972 | Diversity |
| PA7733-1268 | Diversity |
| IA111149 | Diversity |
| X397-1-B5-2\|PI605547 | Diversity |
| GN04399 | Diversity |
| IA111221 | Diversity |
| LANG | Diversity |
| HA05AB38-22 | Diversity |
| ND080724 | Diversity |
| WIR4071 | Diversity |
| OSAGE | Diversity |
| X9384-2 | Diversity |
| IA111165 | Diversity |
| 57A-3 | Diversity |
| OA1180-5 | Diversity |
| MN08251 | Diversity |
| IA111215 | Diversity |
| IA111222 | Diversity |
| LONDRIUnknown | Diversity |
| MN08129 | Diversity |
| IA111002 | Diversity |

| Line Name | Population |
|---|---|
| HA05AB29-17 | Diversity |
| TIPPECANOE | Diversity |
| FL03001BSB-S7 | Diversity |
| LISCHOWER_FRUHHAFER\|CIAV3799 | Diversity |
| SA060716 | Diversity |
| OT399 | Diversity |
| REEVES | Diversity |
| IL00-7070 | Diversity |
| IA111162 | Diversity |
| CORNELLIAN | Diversity |
| SKOROSPELKS | Diversity |
| IL75-5665 | Diversity |
| MN08254 | Diversity |
| PI159172 | Diversity |
| CIAV5925 | Diversity |
| IA111070 | Diversity |
| BENDERY878A\|PI258703 | Diversity |
| CIAV5033 | Diversity |
| IA111137 | Diversity |
| OA1268-3 | Diversity |
| FLAEMINGSNOVA | Diversity |
| FL03129-AB7 | Diversity |
| CDC_ORRIN | Diversity |
| IA111109 | Diversity |
| DUPPAWSKI | Diversity |
| IL98-10145 | Diversity |
| SELECTION3863-8 | Diversity |
| TX02U7605 | Diversity |
| AMES_SELECTION4103 | Diversity |
| M1-7 | Diversity |
| FREDDY | Diversity |
| ND050506 | Diversity |
| CLINTLAND60 | Diversity |
| IA111077 | Diversity |

| Line Name | Population |
|---|---|
| MN08225 | Diversity |
| ND071521 | Diversity |
| IA111012 | Diversity |
| 053B1-95 | Diversity |
| 74C-1 | Diversity |
| ROBUST | Diversity |
| MN08139 | Diversity |
| IA111238 | Diversity |
| IA111155 | Diversity |
| CIAV6227 | Diversity |
| BELINDA | Diversity |
| ABERDEEN_SELECTION1939-3927 | Diversity |
| IL05-3928 | Diversity |
| NEWDAK | Diversity |
| 2A-3 | Diversity |
| ND060464 | Diversity |
| X9258-5 | Diversity |
| 77NZ_AA322 | Diversity |
| IA111121 | Diversity |
| ND051467 | Diversity |
| X9285-1 | Diversity |
| MN08253 | Diversity |
| IA111144 | Diversity |
| IL05-6223 | Diversity |
| SA070452 | Diversity |
| ND051069 | Diversity |
| X9270-4 | Diversity |
| TERRY | Diversity |
| OT3028 | Diversity |
| NES | Diversity |
| ABEGWEIT | Diversity |
| ND051037 | Diversity |
| VALLEY | Diversity |
| HY174-OA | Diversity |

| Line Name | Population |
|---|---|
| CDC_SEABISCUIT | Diversity |
| IL03-2658 | Diversity |
| CIAV5019 | Diversity |
| LAO-1134-022 | Diversity |
| IA111171 | Diversity |
| SA070860 | Diversity |
| IA111138 | Diversity |
| ROUnknownLD | Diversity |
| HA05AB41-38 | Diversity |
| CHERNIGOVSKIJ27B | Diversity |
| VI2 | Diversity |
| P1-9 | Diversity |
| IL2294-8 | Diversity |
| ND050578 | Diversity |
| X9396-4 | Diversity |
| TX02U7047 | Diversity |
| SD751187 | Diversity |
| ODAL | Diversity |
| DELAIR | Diversity |
| IA10033 | Diversity |
| G2-8 | Diversity |
| IA111066 | Diversity |
| PI193957 | Diversity |
| RED_TEXAS\|CIAV1914 | Diversity |
| MISSOURI04103 | Diversity |
| 02G31-NU6D | Diversity |
| CD3708 | Diversity |
| BARAGAN114 | Diversity |
| ND060652 | Diversity |
| KINVARRA_NO_A-8 | Diversity |
| X9410-1 | Diversity |
| OA1130-1 | Diversity |
| MN06108 | Diversity |
| IA111279 | Diversity |

| Line Name | Population |
|---|---|
| IA111236 | Diversity |
| ALLEN | Diversity |
| 02G31-NN4B | Diversity |
| PI436071 | Diversity |
| IA111192 | Diversity |
| FULGRAIN_SELECTION\|CIAV4565 | Diversity |
| 001A1-24-2-4-1-3 | Diversity |
| FL03167BSB-147 | Diversity |
| COLBERSON | Diversity |
| LA03046SBS7-B-S1 | Diversity |
| IA111146 | Diversity |
| ARIANE\|PI361884 | Diversity |
| LEUnknown | Diversity |
| SA070845 | Diversity |
| ND072258 | Diversity |
| SEVERNYJ209 | Diversity |
| TX07CS1268 | Diversity |
| SA01223-02 | Diversity |
| IA111069 | Diversity |
| CAUCAZ4275 | Diversity |
| TX07CS2201 | Diversity |
| TX02U7097 | Diversity |
| X8995-4 | Diversity |
| CIAV4143 | Diversity |
| SECRETARIAT_LA495 | Diversity |
| CD3737 | Diversity |

**Supplemental Table 3.** Percent changes in prediction accuracy over GBLUP of 5, 10, 11, 17 and 17 traits from transcriptomic BLUP (T), metabolomic BLUP (M), G+T, G+M and G+T+M models in the Diversity panel.

| Model | TraitName | Percent Change |
|---|---|---|
| T | Days to Heading | 12.4 |
| T | Seed Length | 17.9 |
| T | Hundred Hull Weight | 21.6 |
| T | C16:1 | 0.7 |
| T | C18:3 | 21.4 |
| M | Days to Heading | 6.0 |
| M | Hundred Hull Weight | 8.0 |
| M | Groat Percentage | 1.9 |
| M | C16:0 | 51.5 |
| M | C18:0 | 58.1 |
| M | C18:1 | 52.0 |
| M | C18:2 | 34.9 |
| M | C18:3 | 23.5 |
| M | C20:0 | 56.3 |
| M | C20:1 | 18.4 |
| G+T | Days to Heading | 0.1 |
| G+T | Seed Length | 24.1 |
| G+T | Hundred Kernel Weight | 0.9 |
| G+T | Hundred Hull Weight | 32.5 |
| G+T | Groat Percentage | 3.0 |
| G+T | C14:0 | 17.7 |
| G+T | C16:0 | 8.4 |
| G+T | C18:0 | 9.7 |
| G+T | C18:1 | 8.2 |
| G+T | C18:3 | 25.4 |
| G+T | C20:0 | 0.7 |
| G+M | Plant Height | 11.3 |
| G+M | Days to Heading | 13.6 |
| G+M | Seed Length | 42.9 |

| Model | TraitName | Percent Change |
| --- | --- | --- |
| G+M | Seed Width | 5.0 |
| G+M | Seed Height | 4.4 |
| G+M | Hundred Kernel Weight | 13.4 |
| G+M | Hundred Hull Weight | 39.7 |
| G+M | Groat Percentage | 14.3 |
| G+M | C14:0 | 54.3 |
| G+M | C16:0 | 63.4 |
| G+M | C16:1 | 3.8 |
| G+M | C18:0 | 70.3 |
| G+M | C18:1 | 58.1 |
| G+M | C18:2 | 43.4 |
| G+M | C18:3 | 48.8 |
| G+M | C20:0 | 64.5 |
| G+M | C20:1 | 28.1 |
| G+T+M | Plant Height | 8.0 |
| G+T+M | Days to Heading | 17.3 |
| G+T+M | Seed Length | 43.2 |
| G+T+M | Seed Width | 6.6 |
| G+T+M | Seed Height | 5.8 |
| G+T+M | Hundred Kernel Weight | 14.5 |
| G+T+M | Hundred Hull Weight | 42.1 |
| G+T+M | Groat Percentage | 15.5 |
| G+T+M | C14:0 | 52.2 |
| G+T+M | C16:0 | 58.1 |
| G+T+M | C16:1 | 6.0 |
| G+T+M | C18:0 | 68.0 |
| G+T+M | C18:1 | 56.0 |
| G+T+M | C18:2 | 42.4 |
| G+T+M | C18:3 | 48.4 |
| G+T+M | C20:0 | 62.0 |
| G+T+M | C20:1 | 26.3 |

**Supplemental Table 4** Annotation of Metabolites identified in the Diversity Panel

| Compound | Retention Time | Super class |
|---|---|---|
| GC.03.0001 | 905.342419 | Organic oxygen compounds |
| GC.03.0002 | 873.6860748 | Organic oxygen compounds |
| GC.03.0003 | 913.280496 | Organic oxygen compounds |
| GC.03.0004 | 1079.375838 | Organic oxygen compounds |
| GC.03.0005 | 608.4465354 | Organic oxygen compounds |
| GC.03.0006 | 615.0757784 | Organic oxygen compounds |
| GC.03.0007 | 622.9920508 | Organic oxygen compounds |
| GC.03.0008 | 562.0521686 | NA |
| GC.03.0009 | 604.3603099 | Organic oxygen compounds |
| GC.03.0010 | 689.5111293 | Organic oxygen compounds |
| GC.03.0011 | 905.3270492 | NA |
| GC.03.0012 | 618.4773554 | NA |
| GC.03.0013 | 973.4837304 | Organic oxygen compounds |
| GC.03.0014 | 426.7172661 | Organic acids and derivatives |
| GC.03.0015 | 554.5475926 | Lipids and lipid-like molecules |
| GC.03.0016 | 582.2242427 | Organic oxygen compounds |
| GC.03.0017 | 913.19679 | NA |
| GC.03.0018 | 235.2794138 | Homogeneous non-metal compounds |
| GC.03.0019 | 512.4669012 | Organic acids and derivatives |
| GC.03.0020 | 526.9852785 | NA |
| GC.03.0021 | 416.0852179 | NA |
| GC.03.0022 | 720.6606712 | Lipids and lipid-like molecules |
| GC.03.0023 | 317.2797887 | Homogeneous non-metal compounds |
| GC.03.0024 | 578.4344783 | Organic acids and derivatives |
| GC.03.0025 | 873.5896212 | NA |
| GC.03.0026 | 488.1969242 | Organic acids and derivatives |
| GC.03.0027 | 785.2558571 | NA |
| GC.03.0029 | 265.8842459 | Organic acids and derivatives |
| GC.03.0030 | 615.115 | Organic oxygen compounds |
| GC.03.0031 | 1079.578789 | NA |
| GC.03.0032 | 873.6081964 | NA |
| GC.03.0033 | 913.213625 | NA |
| GC.03.0034 | 722.9986364 | Organic acids and derivatives |

| Compound | Retention Time | Super class |
|---|---|---|
| GC.03.0035 | 973.4794375 | NA |
| GC.03.0036 | 317.1070217 | Organic oxygen compounds |
| GC.03.0037 | 613.7480435 | Organic oxygen compounds |
| GC.03.0038 | 905.3105455 | NA |
| GC.03.0039 | 615.0139545 | NA |
| GC.03.0040 | 517.92075 | NA |
| GC.03.0041 | 317.229119 | Organic oxygen compounds |
| GC.03.0042 | 731.4316829 | NA |
| GC.03.0043 | 628.381475 | Organic oxygen compounds |
| GC.03.0044 | 543.4971 | NA |
| GC.03.0046 | 893.4089487 | NA |
| GC.03.0047 | 604.6133421 | NA |
| GC.03.0048 | 604.368 | Organic acids and derivatives |
| GC.03.0050 | 689.4509459 | NA |
| GC.03.0051 | 803.7083514 | Organic oxygen compounds |
| GC.03.0052 | 659.4322222 | Lipids and lipid-like molecules |
| GC.03.0053 | 263.9096765 | NA |
| GC.03.0054 | 905.4049091 | NA |
| GC.03.0055 | 442.8898438 | Organic acids and derivatives |
| GC.03.0056 | 622.9176129 | NA |
| GC.03.0057 | 607.9156774 | Organoheterocyclic compounds |
| GC.03.0058 | 412.8580323 | NA |
| GC.03.0059 | 704.0695484 | NA |
| GC.03.0060 | 659.3991 | NA |
| GC.03.0061 | 350.137 | NA |
| GC.03.0062 | 761.0765333 | NA |
| GC.03.0063 | 628.8161724 | NA |
| GC.03.0064 | 392.5553448 | NA |
| GC.03.0066 | 338.6081111 | NA |
| GC.03.0067 | 641.2526667 | NA |
| GC.03.0069 | 873.5831923 | NA |
| GC.03.0070 | 647.3682692 | NA |
| GC.03.0071 | 618.4015385 | NA |
| GC.03.0072 | 628.4966923 | NA |

| Compound | Retention Time | Super class |
|---|---|---|
| GC.03.0073 | 1079.40664 | NA |
| GC.03.0075 | 189.03824 | NA |
| GC.03.0076 | 623.17728 | NA |
| GC.03.0077 | 467.44088 | NA |
| GC.03.0078 | 731.15488 | Organoheterocyclic compounds |
| GC.03.0079 | 1079.38425 | NA |
| GC.03.0080 | 608.1451667 | NA |
| GC.03.0081 | 531.9253333 | NA |
| GC.03.0082 | 562.0827083 | NA |
| GC.03.0083 | 740.2407083 | NA |
| GC.03.0084 | 792.018625 | NA |
| GC.03.0085 | 1079.578957 | NA |
| GC.03.0086 | 631.8909565 | Organic acids and derivatives |
| GC.03.0087 | 785.402 | NA |
| GC.03.0088 | 209.7632273 | NA |
| GC.03.0089 | 631.0757273 | NA |
| GC.03.0090 | 427.4815 | NA |
| GC.03.0091 | 721.4842381 | Lipids and lipid-like molecules |
| GC.03.0092 | 873.5687 | NA |
| GC.03.0094 | 513.1085 | Organic oxygen compounds |
| GC.03.0095 | 233.8148947 | NA |
| GC.03.0096 | 235.2594211 | Homogeneous non-metal compounds |
| GC.03.0097 | 606.3486842 | NA |
| GC.03.0098 | 722.9605789 | NA |
| GC.03.0099 | 898.6029444 | NA |
| GC.03.0100 | 246.4493889 | NA |
| GC.03.0101 | 246.5315556 | NA |
| GC.03.0102 | 664.4071667 | NA |
| GC.03.0103 | 615.0996111 | NA |
| GC.03.0104 | 432.1025556 | Organic acids and derivatives |
| GC.03.0105 | 572.2538333 | NA |
| GC.03.0106 | 864.1787778 | NA |
| GC.03.0107 | 206.4792941 | NA |
| GC.03.0108 | 188.7794118 | NA |

| Compound | Retention Time | Super class |
|---|---|---|
| GC.03.0109 | 308.4041765 | NA |
| GC.03.0110 | 284.8552353 | NA |
| GC.03.0111 | 596.7862941 | NA |
| GC.03.0112 | 381.0474118 | NA |
| GC.03.0113 | 496.5770588 | Organic acids and derivatives |
| GC.03.0114 | 460.4919412 | NA |
| GC.03.0115 | 561.9962353 | NA |
| GC.03.0117 | 595.261625 | NA |
| GC.03.0118 | 485.7451875 | NA |
| GC.03.0119 | 582.1916875 | NA |
| GC.03.0120 | 832.465 | NA |
| GC.03.0121 | 203.3900667 | NA |
| GC.03.0123 | 240.4784 | NA |
| GC.03.0124 | 335.339 | NA |
| GC.03.0125 | 263.3005333 | NA |
| GC.03.0126 | 656.6228667 | NA |
| GC.03.0127 | 618.1748667 | Organoheterocyclic compounds |
| GC.03.0128 | 630.5423333 | NA |
| GC.03.0129 | 634.2838667 | Organic oxygen compounds |
| GC.03.0131 | 444.79 | NA |
| GC.03.0132 | 582.3156667 | NA |
| GC.03.0133 | 553.9890667 | Organic oxygen compounds |
| GC.03.0134 | 942.3575 | NA |
| GC.03.0135 | 892.7530714 | NA |
| GC.03.0136 | 335.219 | Organic acids and derivatives |
| GC.03.0137 | 679.1025 | NA |
| GC.03.0138 | 618.2345 | Organic acids and derivatives |
| GC.03.0139 | 604.5229286 | NA |
| GC.03.0140 | 377.0225 | Organic acids and derivatives |
| GC.03.0141 | 413.0057143 | NA |
| GC.03.0143 | 819.23 | NA |
| GC.03.0144 | 1101.444846 | NA |
| GC.03.0145 | 972.3684615 | NA |
| GC.03.0146 | 873.6438462 | NA |

| Compound | Retention Time | Super class |
|---|---|---|
| GC.03.0147 | 879.5807692 | NA |
| GC.03.0148 | 940.225 | NA |
| GC.03.0149 | 194.9468462 | Organoheterocyclic compounds |
| GC.03.0150 | 246.4186923 | NA |
| GC.03.0152 | 301.2323846 | NA |
| GC.03.0153 | 293.6822308 | NA |
| GC.03.0154 | 628.313 | NA |
| GC.03.0156 | 427.171 | NA |
| GC.03.0157 | 443.808 | Organic acids and derivatives |
| GC.03.0158 | 476.6702308 | Organic acids and derivatives |
| GC.03.0161 | 968.5646667 | NA |
| GC.03.0162 | 972.1901667 | NA |
| GC.03.0163 | 917.8673333 | NA |
| GC.03.0165 | 227.0151667 | NA |
| GC.03.0166 | 284.8361667 | NA |
| GC.03.0167 | 614.381 | NA |
| GC.03.0168 | 618.1998333 | NA |
| GC.03.0169 | 604.8261667 | NA |
| GC.03.0170 | 598.7845 | NA |
| GC.03.0171 | 476.1353333 | Organic acids and derivatives |
| GC.03.0172 | 472.0796667 | NA |
| GC.03.0173 | 689.3560833 | Organoheterocyclic compounds |
| GC.03.0174 | 710.2935 | NA |
| GC.03.0175 | 767.903 | NA |
| GC.03.0176 | 807.1899167 | NA |
| GC.03.0178 | 1092.359727 | NA |
| GC.03.0179 | 181.0630909 | NA |
| GC.03.0180 | 673.1062727 | NA |
| GC.03.0181 | 604.7054545 | NA |
| GC.03.0182 | 609.2276364 | NA |
| GC.03.0183 | 608.609 | NA |
| GC.03.0184 | 598.5461818 | NA |
| GC.03.0185 | 731.5755455 | NA |
| GC.03.0186 | 760.5769091 | NA |

| Compound | Retention Time | Super class |
|---|---|---|
| GC.03.0187 | 1004.9993 | NA |
| GC.03.0189 | 205.5885 | NA |
| GC.03.0190 | 313.7486 | Organic nitrogen compounds |
| GC.03.0191 | 284.7833 | NA |
| GC.03.0192 | 295.9964 | NA |
| GC.03.0193 | 670.4067 | NA |
| GC.03.0194 | 675.2756 | NA |
| GC.03.0195 | 655.9534 | NA |
| GC.03.0196 | 625.1102 | Organic acids and derivatives |
| GC.03.0197 | 635.9493 | NA |
| GC.03.0198 | 426.6229 | NA |
| GC.03.0200 | 446.7036 | Organic acids and derivatives |
| GC.03.0201 | 450.1565 | NA |
| GC.03.0202 | 461.6541 | NA |
| GC.03.0203 | 456.0132 | NA |
| GC.03.0204 | 522.9084 | NA |
| GC.03.0205 | 814.038 | NA |
| GC.03.0206 | 819.2587 | NA |
| GC.03.0207 | 1078.298 | NA |
| GC.03.0208 | 1100.992333 | NA |
| GC.03.0209 | 1068.961222 | NA |
| GC.03.0210 | 973.8846667 | NA |
| GC.03.0211 | 943.3602222 | NA |
| GC.03.0212 | 909.169 | NA |
| GC.03.0213 | 326.7267778 | Organoheterocyclic compounds |
| GC.03.0215 | 360.925 | NA |
| GC.03.0216 | 488.6055556 | NA |
| GC.03.0217 | 479.5762222 | NA |
| GC.03.0218 | 460.9891111 | NA |
| GC.03.0220 | 730.1925556 | Lipids and lipid-like molecules |
| GC.03.0221 | 756.8393333 | NA |
| GC.03.0222 | 788.3192222 | NA |
| GC.03.0223 | 1084.893875 | NA |
| GC.03.0224 | 1004.480125 | NA |

| Compound | Retention Time | Super class |
|---|---|---|
| GC.03.0225 | 973.410375 | NA |
| GC.03.0226 | 877.066375 | NA |
| GC.03.0227 | 935.778 | NA |
| GC.03.0228 | 194.964875 | NA |
| GC.03.0229 | 249.973125 | NA |
| GC.03.0230 | 314.811125 | NA |
| GC.03.0231 | 312.413875 | NA |
| GC.03.0232 | 622.81 | NA |
| GC.03.0233 | 622.87675 | NA |
| GC.03.0234 | 426.713625 | NA |
| GC.03.0235 | 441.438625 | NA |
| GC.03.0237 | 577.382 | NA |
| GC.03.0238 | 695.023625 | NA |
| GC.03.0239 | 725.275125 | NA |
| GC.03.0241 | 826.794375 | NA |
| GC.03.0242 | 1051.95 | NA |
| GC.03.0243 | 1005.120571 | NA |
| GC.03.0244 | 1025.348 | NA |
| GC.03.0245 | 996.541 | NA |
| GC.03.0247 | 228.9065714 | NA |
| GC.03.0248 | 218.4891429 | NA |
| GC.03.0249 | 327.7115714 | NA |
| GC.03.0250 | 673.1548571 | NA |
| GC.03.0251 | 623.7868571 | NA |
| GC.03.0252 | 622.707 | NA |
| GC.03.0253 | 604.2615714 | NA |
| GC.03.0254 | 363.3385714 | Organic acids and derivatives |
| GC.03.0255 | 490.4192857 | NA |
| GC.03.0256 | 442.2804286 | NA |
| GC.03.0257 | 444.5221429 | NA |
| GC.03.0258 | 445.0444286 | NA |
| GC.03.0259 | 477.2072857 | NA |
| GC.03.0260 | 466.036 | Organic acids and derivatives |
| GC.03.0261 | 523.5861429 | NA |

| Compound | Retention Time | Super class |
|---|---|---|
| GC.03.0262 | 565.2338571 | NA |
| GC.03.0263 | 686.191 | NA |
| GC.03.0264 | 714.9767143 | NA |
| GC.03.0265 | 1094.098333 | NA |
| GC.03.0266 | 1100.736667 | NA |
| GC.03.0267 | 1280.6235 | NA |
| GC.03.0269 | 987.5375 | NA |
| GC.03.0270 | 873.631 | NA |
| GC.03.0271 | 878.1773333 | NA |
| GC.03.0272 | 877.0715 | NA |
| GC.03.0274 | 891.6821667 | NA |
| GC.03.0276 | 194.5426667 | NA |
| GC.03.0277 | 183.1735 | NA |
| GC.03.0278 | 247.3016667 | NA |
| GC.03.0279 | 221.6095 | NA |
| GC.03.0280 | 259.7858333 | NA |
| GC.03.0281 | 268.316 | NA |
| GC.03.0282 | 644.6096667 | NA |
| GC.03.0283 | 623.0313333 | NA |
| GC.03.0285 | 631.5475 | NA |
| GC.03.0286 | 351.7346667 | Organic acids and derivatives |
| GC.03.0287 | 402.2008333 | NA |
| GC.03.0288 | 426.8566667 | NA |
| GC.03.0289 | 443.5591667 | NA |
| GC.03.0290 | 443.99 | NA |
| GC.03.0291 | 474.8458333 | NA |
| GC.03.0292 | 450.0641667 | NA |
| GC.03.0293 | 512.752 | NA |
| GC.03.0294 | 512.724 | NA |
| GC.03.0295 | 558.9558333 | NA |
| GC.03.0296 | 733.558 | NA |
| GC.03.0297 | 730.4918333 | NA |
| GC.03.0298 | 800.3888333 | NA |
| GC.03.0299 | 834.6165 | NA |

| Compound | Retention Time | Super class |
|---|---|---|
| GC.03.0300 | 863.8686667 | NA |
| GC.03.0301 | 1079.201 | NA |
| GC.03.0302 | 1094.0334 | NA |
| GC.03.0303 | 1130.0212 | NA |
| GC.03.0304 | 1127.9542 | NA |
| GC.03.0305 | 1163.1256 | NA |
| GC.03.0306 | 1244.5964 | NA |
| GC.03.0307 | 1056.609 | NA |
| GC.03.0310 | 981.745 | NA |
| GC.03.0311 | 879.2638 | NA |
| GC.03.0312 | 893.073 | NA |
| GC.03.0313 | 195.2622 | NA |
| GC.03.0314 | 199.7622 | Organic acids and derivatives |
| GC.03.0315 | 184.9592 | NA |
| GC.03.0316 | 246.8122 | NA |
| GC.03.0317 | 246.5346 | NA |
| GC.03.0318 | 330.6656 | NA |
| GC.03.0319 | 269.8078 | NA |
| GC.03.0320 | 296.0504 | NA |
| GC.03.0321 | 662.4996 | NA |
| GC.03.0322 | 644.958 | NA |
| GC.03.0323 | 613.7746 | NA |
| GC.03.0324 | 622.7488 | NA |
| GC.03.0325 | 631.1318 | NA |
| GC.03.0326 | 608.2534 | NA |
| GC.03.0327 | 595.037 | NA |
| GC.03.0328 | 596.7488 | NA |
| GC.03.0329 | 388.4514 | NA |
| GC.03.0330 | 488.2368 | NA |
| GC.03.0331 | 493.2632 | Organic acids and derivatives |
| GC.03.0332 | 432.1688 | NA |
| GC.03.0333 | 433.127 | NA |
| GC.03.0334 | 439.1348 | NA |
| GC.03.0335 | 435.684 | NA |

| Compound | Retention Time | Super class |
|---|---|---|
| GC.03.0336 | 435.3612 | NA |
| GC.03.0337 | 459.0384 | NA |
| GC.03.0338 | 519.0626 | NA |
| GC.03.0340 | 551.7124 | NA |
| GC.03.0343 | 717.1176 | NA |
| GC.03.0344 | 715.5446 | NA |
| GC.03.0345 | 739.7466 | NA |
| GC.03.0347 | 762.1498 | NA |
| GC.03.0348 | 754.2912 | NA |
| GC.03.0350 | 863.9002 | NA |
| GC.03.0351 | 840.9386 | NA |
| GC.03.0352 | 1082.608 | NA |
| GC.03.0353 | 1124.36225 | NA |
| GC.03.0354 | 1136.354 | NA |
| GC.03.0355 | 1280.7175 | NA |
| GC.03.0356 | 1025.72525 | NA |
| GC.03.0358 | 966.374 | NA |
| GC.03.0359 | 988.6985 | NA |
| GC.03.0360 | 981.723 | NA |
| GC.03.0361 | 873.561 | NA |
| GC.03.0362 | 883.904 | NA |
| GC.03.0365 | 883.28175 | NA |
| GC.03.0366 | 886.9605 | NA |
| GC.03.0367 | 934.69775 | NA |
| GC.03.0368 | 891.4185 | NA |
| GC.03.0370 | 920.947 | NA |
| GC.03.0371 | 210.04125 | NA |
| GC.03.0372 | 191.25025 | NA |
| GC.03.0373 | 254.05275 | NA |
| GC.03.0375 | 246.93825 | NA |
| GC.03.0376 | 246.54625 | NA |
| GC.03.0377 | 249.331 | NA |
| GC.03.0378 | 223.194 | NA |
| GC.03.0379 | 314.111 | NA |

| Compound | Retention Time | Super class |
|---|---|---|
| GC.03.0380 | 302.46375 | NA |
| GC.03.0381 | 296.22475 | NA |
| GC.03.0382 | 675.47575 | NA |
| GC.03.0383 | 674.29525 | NA |
| GC.03.0384 | 665.9075 | NA |
| GC.03.0385 | 655.71375 | NA |
| GC.03.0386 | 645.0745 | Organoheterocyclic compounds |
| GC.03.0387 | 637.2755 | NA |
| GC.03.0388 | 605.3515 | NA |
| GC.03.0389 | 363.18825 | Organic acids and derivatives |
| GC.03.0390 | 365.58475 | Organoheterocyclic compounds |
| GC.03.0391 | 378.51625 | NA |
| GC.03.0393 | 389.81175 | NA |
| GC.03.0394 | 391.336 | NA |
| GC.03.0397 | 480.44975 | NA |
| GC.03.0398 | 482.176 | NA |
| GC.03.0399 | 471.09075 | Organic acids and derivatives |
| GC.03.0400 | 464.094 | NA |
| GC.03.0401 | 457.2385 | NA |
| GC.03.0402 | 536.88525 | NA |
| GC.03.0403 | 543.9925 | NA |
| GC.03.0404 | 582.9045 | NA |
| GC.03.0406 | 560.33375 | NA |
| GC.03.0408 | 707.4015 | NA |
| GC.03.0409 | 702.86125 | NA |
| GC.03.0410 | 715.51 | NA |
| GC.03.0411 | 724.2105 | NA |
| GC.03.0412 | 748.74025 | NA |
| GC.03.0413 | 736.8835 | NA |
| GC.03.0414 | 766.417 | NA |
| GC.03.0415 | 798.72375 | NA |
| GC.03.0416 | 843.2145 | NA |
| GC.03.0417 | 851.511 | NA |
| GC.03.0418 | 1081.232333 | NA |

| Compound | Retention Time | Super class |
|---|---|---|
| GC.03.0419 | 1086.226667 | NA |
| GC.03.0420 | 1083.263 | NA |
| GC.03.0421 | 1089.906333 | NA |
| GC.03.0423 | 1102.091 | NA |
| GC.03.0424 | 1123.993 | NA |
| GC.03.0428 | 1107.434333 | NA |
| GC.03.0430 | 1137.665 | NA |
| GC.03.0431 | 1156.337667 | NA |
| GC.03.0433 | 1262.054 | NA |
| GC.03.0434 | 1319.258 | NA |
| GC.03.0437 | 1060.916333 | NA |
| GC.03.0439 | 1052.597 | NA |
| GC.03.0441 | 1038.898333 | NA |
| GC.03.0442 | 1040.258667 | NA |
| GC.03.0443 | 1034.104 | NA |
| GC.03.0445 | 962.382 | NA |
| GC.03.0452 | 882.0203333 | NA |
| GC.03.0453 | 938.6636667 | NA |
| GC.03.0454 | 940.42 | NA |
| GC.03.0455 | 893.1216667 | NA |
| GC.03.0456 | 917.2693333 | NA |
| GC.03.0457 | 921.737 | NA |
| GC.03.0458 | 920.0956667 | NA |
| GC.03.0459 | 208.616 | NA |
| GC.03.0460 | 182.455 | NA |
| GC.03.0461 | 188.8603333 | NA |
| GC.03.0462 | 235.7036667 | NA |
| GC.03.0463 | 323.4 | NA |
| GC.03.0464 | 275.1286667 | NA |
| GC.03.0465 | 281.1256667 | NA |
| GC.03.0466 | 291.202 | NA |
| GC.03.0467 | 668.8576667 | NA |
| GC.03.0468 | 641.913 | NA |
| GC.03.0469 | 640.053 | NA |

| Compound | Retention Time | Super class |
|---|---|---|
| GC.03.0471 | 621.667 | Nucleosides, nucleotides, and analogues |
| GC.03.0473 | 347.1076667 | NA |
| GC.03.0474 | 355.275 | NA |
| GC.03.0475 | 372.5243333 | NA |
| GC.03.0476 | 396.2766667 | NA |
| GC.03.0478 | 420.5663333 | NA |
| GC.03.0480 | 497.2276667 | NA |
| GC.03.0481 | 443.8643333 | NA |
| GC.03.0482 | 432.495 | NA |
| GC.03.0483 | 437.1293333 | NA |
| GC.03.0484 | 470.0406667 | NA |
| GC.03.0485 | 453.0446667 | NA |
| GC.03.0486 | 534.1453333 | NA |
| GC.03.0488 | 547.064 | NA |
| GC.03.0491 | 586.896 | NA |
| GC.03.0492 | 584.338 | NA |
| GC.03.0493 | 565.8193333 | NA |
| GC.03.0494 | 554.077 | NA |
| GC.03.0495 | 555.001 | NA |
| GC.03.0496 | 556.6086667 | NA |
| GC.03.0498 | 727.9223333 | NA |
| GC.03.0500 | 806.621 | NA |
| GC.03.0501 | 803.4856667 | NA |
| GC.03.0502 | 796.653 | NA |
| GC.03.0503 | 794.8743333 | NA |
| GC.03.0505 | 1507.061 | NA |
| GC.03.0506 | 1505.7175 | NA |
| GC.03.0510 | 1447.731 | NA |
| GC.03.0511 | 1453.019 | NA |
| GC.03.0514 | 1440.88 | NA |
| GC.03.0517 | 1499.0775 | NA |
| GC.03.0518 | 1497.005 | NA |
| GC.03.0520 | 1081.8115 | NA |
| GC.03.0521 | 1080.897 | NA |

| Compound | Retention Time | Super class |
|---|---|---|
| GC.03.0522 | 1080.568 | NA |
| GC.03.0523 | 1080.5895 | NA |
| GC.03.0524 | 1079.4165 | NA |
| GC.03.0525 | 1087.061 | NA |
| GC.03.0528 | 1087.2835 | NA |
| GC.03.0531 | 1085.512 | NA |
| GC.03.0532 | 1084.3415 | NA |
| GC.03.0533 | 1090.57 | NA |
| GC.03.0535 | 1094.228 | NA |
| GC.03.0537 | 1094.451 | NA |
| GC.03.0540 | 1094.8665 | NA |
| GC.03.0543 | 1092.308 | NA |
| GC.03.0545 | 1091.455 | NA |
| GC.03.0548 | 1100.317 | NA |
| GC.03.0549 | 1099.996 | NA |
| GC.03.0550 | 1100.124 | NA |
| GC.03.0551 | 1101.866 | NA |
| GC.03.0552 | 1101.911 | NA |
| GC.03.0556 | 1101.3125 | NA |
| GC.03.0557 | 1129.5605 | NA |
| GC.03.0559 | 1122.5225 | NA |
| GC.03.0561 | 1127.4445 | NA |
| GC.03.0562 | 1125.1655 | NA |
| GC.03.0564 | 1126.174 | NA |
| GC.03.0565 | 1119.4915 | NA |
| GC.03.0567 | 1114.5935 | NA |
| GC.03.0568 | 1113.4565 | NA |
| GC.03.0569 | 1114.053 | NA |
| GC.03.0571 | 1112.0295 | NA |
| GC.03.0573 | 1108.224 | NA |
| GC.03.0574 | 1108.3875 | NA |
| GC.03.0576 | 1136.4615 | NA |
| GC.03.0577 | 1136.1075 | NA |
| GC.03.0578 | 1136.3315 | NA |

| Compound | Retention Time | Super class |
|---|---|---|
| GC.03.0579 | 1135.3305 | NA |
| GC.03.0580 | 1135.984 | NA |
| GC.03.0582 | 1138.107 | NA |
| GC.03.0583 | 1137.66 | NA |
| GC.03.0584 | 1137.0845 | NA |
| GC.03.0585 | 1136.7075 | NA |
| GC.03.0590 | 1149.0065 | NA |
| GC.03.0591 | 1148.9255 | NA |
| GC.03.0592 | 1148.008 | NA |
| GC.03.0593 | 1151.455 | NA |
| GC.03.0594 | 1149.724 | NA |
| GC.03.0596 | 1165.0225 | NA |
| GC.03.0598 | 1161.359 | NA |
| GC.03.0599 | 1161.627 | NA |
| GC.03.0603 | 1154.6745 | NA |
| GC.03.0604 | 1168.353 | NA |
| GC.03.0605 | 1170.2275 | NA |
| GC.03.0607 | 1185.3105 | NA |
| GC.03.0609 | 1182.9055 | NA |
| GC.03.0612 | 1225.5135 | NA |
| GC.03.0613 | 1226.596 | NA |
| GC.03.0616 | 1203.4205 | NA |
| GC.03.0621 | 1245.6655 | NA |
| GC.03.0623 | 1280.7505 | NA |
| GC.03.0626 | 1311.237 | NA |
| GC.03.0627 | 1314.4905 | NA |
| GC.03.0628 | 1322.559 | NA |
| GC.03.0629 | 1327.3725 | NA |
| GC.03.0632 | 1302.2585 | NA |
| GC.03.0634 | 1373.1765 | NA |
| GC.03.0635 | 1369.5795 | NA |
| GC.03.0637 | 1339.252 | NA |
| GC.03.0641 | 1064.5965 | NA |
| GC.03.0642 | 1065.136 | NA |

| Compound | Retention Time | Super class |
|---|---|---|
| GC.03.0643 | 1066.8795 | NA |
| GC.03.0644 | 1056.5305 | NA |
| GC.03.0645 | 1051.8145 | Nucleosides, nucleotides, and analogues |
| GC.03.0646 | 1053.913 | NA |
| GC.03.0647 | 1008.8785 | NA |
| GC.03.0648 | 1005.6365 | NA |
| GC.03.0649 | 1006.284 | NA |
| GC.03.0650 | 1006.7795 | NA |
| GC.03.0651 | 1005.3445 | NA |
| GC.03.0653 | 1013.815 | NA |
| GC.03.0658 | 1025.9895 | NA |
| GC.03.0659 | 1026.5455 | NA |
| GC.03.0661 | 1028.216 | NA |
| GC.03.0662 | 1031.3965 | NA |
| GC.03.0663 | 1041.6035 | NA |
| GC.03.0666 | 951.3515 | NA |
| GC.03.0667 | 950.025 | NA |
| GC.03.0670 | 963.062 | NA |
| GC.03.0671 | 956.2795 | NA |
| GC.03.0672 | 956.9945 | NA |
| GC.03.0673 | 966.9 | NA |
| GC.03.0678 | 972.179 | NA |
| GC.03.0680 | 973.3825 | NA |
| GC.03.0682 | 991.6155 | NA |
| GC.03.0683 | 978.3865 | NA |
| GC.03.0684 | 985.1345 | NA |
| GC.03.0686 | 985.4665 | NA |
| GC.03.0688 | 984.259 | NA |
| GC.03.0689 | 984.065 | NA |
| GC.03.0692 | 981.8155 | NA |
| GC.03.0693 | 981.8805 | NA |
| GC.03.0696 | 980.0325 | NA |
| GC.03.0698 | 979.784 | NA |
| GC.03.0699 | 873.5935 | NA |

| Compound | Retention Time | Super class |
|---|---|---|
| GC.03.0702 | 884.332 | NA |
| GC.03.0703 | 883.281 | Phenylpropanoids and polyketides |
| GC.03.0707 | 933.543 | NA |
| GC.03.0708 | 943.0695 | NA |
| GC.03.0709 | 939.6265 | NA |
| GC.03.0713 | 893.4225 | NA |
| GC.03.0714 | 894.408 | NA |
| GC.03.0715 | 895.2145 | NA |
| GC.03.0716 | 900.8085 | NA |
| GC.03.0717 | 898.6835 | NA |
| GC.03.0718 | 909.483 | NA |
| GC.03.0720 | 929.686 | NA |
| GC.03.0722 | 198.202 | NA |
| GC.03.0723 | 181.0715 | NA |
| GC.03.0725 | 232.982 | NA |
| GC.03.0727 | 257.821 | NA |
| GC.03.0728 | 286.482 | NA |
| GC.03.0729 | 680.6805 | Phenylpropanoids and polyketides |
| GC.03.0731 | 660.1215 | NA |
| GC.03.0734 | 614.445 | NA |
| GC.03.0735 | 618.2345 | NA |
| GC.03.0736 | 630.2255 | NA |
| GC.03.0737 | 638.0565 | NA |
| GC.03.0738 | 604.3975 | NA |
| GC.03.0739 | 592.343 | NA |
| GC.03.0740 | 353.5505 | NA |
| GC.03.0741 | 378.55 | NA |
| GC.03.0743 | 389.1205 | NA |
| GC.03.0745 | 400.76 | NA |
| GC.03.0746 | 406.86 | NA |
| GC.03.0747 | 413.1325 | NA |
| GC.03.0748 | 415.691 | NA |
| GC.03.0749 | 429.02 | NA |
| GC.03.0751 | 499.4625 | NA |

| Compound | Retention Time | Super class |
|---|---|---|
| GC.03.0753 | 438.068 | NA |
| GC.03.0754 | 527.18 | NA |
| GC.03.0755 | 523.557 | NA |
| GC.03.0756 | 536.612 | NA |
| GC.03.0757 | 569.6815 | NA |
| GC.03.0762 | 700.75 | NA |
| GC.03.0763 | 715.4665 | NA |
| GC.03.0764 | 715.4805 | NA |
| GC.03.0765 | 714.951 | NA |
| GC.03.0766 | 725.451 | NA |
| GC.03.0768 | 747.856 | NA |
| GC.03.0769 | 748.942 | NA |
| GC.03.0771 | 740.136 | NA |
| GC.03.0774 | 762.131 | NA |
| GC.03.0776 | 753.5765 | NA |
| GC.03.0779 | 831.594 | NA |
| GC.03.0780 | 833.7625 | NA |
| GC.03.0781 | 828.3125 | NA |
| GC.03.0782 | 812.4275 | NA |
| GC.03.0783 | 814.0175 | NA |
| GC.03.0784 | 818.905 | NA |
| GC.03.0786 | 865.552 | NA |
| GC.03.0787 | 864.103 | NA |
| GC.03.0788 | 842.8385 | NA |
| GC.03.0790 | 853.7805 | NA |
| GC.03.0791 | 856.4775 | NA |
| GC.03.0792 | 847.2415 | NA |
| LC.03.0001 | 342.2560634 | Lipids and lipid-like molecules |
| LC.03.0002 | 473.8421209 | NA |
| LC.03.0003 | 749.0235172 | Lipids and lipid-like molecules |
| LC.03.0004 | 188.265942 | Lipids and lipid-like molecules |
| LC.03.0005 | 342.285 | NA |
| LC.03.0006 | 689.0814667 | NA |
| LC.03.0007 | 680.6349322 | Lipids and lipid-like molecules |

| Compound | Retention Time | Super class |
|---|---|---|
| LC.03.0008 | 498.8051897 | NA |
| LC.03.0009 | 550.7769649 | Lipids and lipid-like molecules |
| LC.03.0010 | 18.09253571 | NA |
| LC.03.0011 | 476.60474 | Lipids and lipid-like molecules |
| LC.03.0012 | 484.5197755 | Lipids and lipid-like molecules |
| LC.03.0013 | 200.7148333 | NA |
| LC.03.0014 | 521.5479333 | Lipids and lipid-like molecules |
| LC.03.0015 | 23.47621951 | Organic oxygen compounds |
| LC.03.0016 | 704.3701795 | Lipids and lipid-like molecules |
| LC.03.0017 | 711.3491795 | Lipids and lipid-like molecules |
| LC.03.0018 | 224.5752105 | NA |
| LC.03.0019 | 311.2088421 | Lipids and lipid-like molecules |
| LC.03.0020 | 588.0414865 | Benzenoids |
| LC.03.0022 | 696.0413243 | NA |
| LC.03.0023 | 187.6288108 | Organic acids and derivatives |
| LC.03.0024 | 338.1327222 | Lipids and lipid-like molecules |
| LC.03.0025 | 17.98617143 | NA |
| LC.03.0026 | 618.6593714 | NA |
| LC.03.0027 | 857.9161143 | Lipids and lipid-like molecules |
| LC.03.0028 | 761.6957714 | Lipids and lipid-like molecules |
| LC.03.0031 | 195.5017143 | NA |
| LC.03.0032 | 608.8271765 | Organoheterocyclic compounds |
| LC.03.0033 | 863.9952059 | Lipids and lipid-like molecules |
| LC.03.0034 | 388.1878529 | Lipids and lipid-like molecules |
| LC.03.0036 | 659.5264545 | NA |
| LC.03.0037 | 383.9499688 | NA |
| LC.03.0038 | 612.4354516 | Organoheterocyclic compounds |
| LC.03.0039 | 443.2282581 | Lipids and lipid-like molecules |
| LC.03.0040 | 496.2920968 | Lipids and lipid-like molecules |
| LC.03.0041 | 18.04603333 | NA |
| LC.03.0043 | 727.6321 | Phenylpropanoids and polyketides |
| LC.03.0044 | 756.8296 | Lipids and lipid-like molecules |
| LC.03.0045 | 452.6413 | Lipids and lipid-like molecules |
| LC.03.0046 | 23.55168966 | Organic oxygen compounds |

| Compound | Retention Time | Super class |
|---|---|---|
| LC.03.0047 | 1005.00969 | NA |
| LC.03.0048 | 23.62496296 | Organoheterocyclic compounds |
| LC.03.0049 | 773.9751481 | Lipids and lipid-like molecules |
| LC.03.0050 | 397.0408519 | Lipids and lipid-like molecules |
| LC.03.0051 | 394.4412222 | Lipids and lipid-like molecules |
| LC.03.0052 | 349.1564815 | Lipids and lipid-like molecules |
| LC.03.0053 | 840.1081154 | Lipids and lipid-like molecules |
| LC.03.0054 | 764.8521923 | Lipids and lipid-like molecules |
| LC.03.0055 | 286.7376154 | Lignans, neolignans and related compounds |
| LC.03.0056 | 840.90832 | Lipids and lipid-like molecules |
| LC.03.0057 | 846.69404 | Lipids and lipid-like molecules |
| LC.03.0058 | 855.2888 | Lipids and lipid-like molecules |
| LC.03.0059 | 749.00476 | Lipids and lipid-like molecules |
| LC.03.0060 | 250.90144 | Phenylpropanoids and polyketides |
| LC.03.0061 | 338.16 | Lipids and lipid-like molecules |
| LC.03.0062 | 587.6154167 | Lipids and lipid-like molecules |
| LC.03.0063 | 578.7604167 | Organoheterocyclic compounds |
| LC.03.0064 | 830.2118333 | NA |
| LC.03.0066 | 422.3746522 | Lipids and lipid-like molecules |
| LC.03.0067 | 457.368913 | Lipids and lipid-like molecules |
| LC.03.0068 | 578.943 | Hydrocarbons |
| LC.03.0069 | 709.7594091 | Lipids and lipid-like molecules |
| LC.03.0070 | 170.8471818 | NA |
| LC.03.0071 | 583.1299524 | Lipids and lipid-like molecules |
| LC.03.0072 | 642.0361429 | NA |
| LC.03.0073 | 849.1078571 | Lipids and lipid-like molecules |
| LC.03.0075 | 755.4849524 | Lipids and lipid-like molecules |
| LC.03.0076 | 774.0224286 | Lipids and lipid-like molecules |
| LC.03.0077 | 777.0487619 | Lipids and lipid-like molecules |
| LC.03.0078 | 469.4495714 | Lipids and lipid-like molecules |
| LC.03.0079 | 482.123381 | Lipids and lipid-like molecules |
| LC.03.0080 | 716.7721 | Lipids and lipid-like molecules |
| LC.03.0081 | 748.8587 | Lipids and lipid-like molecules |
| LC.03.0082 | 466.45205 | Lipids and lipid-like molecules |

| Compound | Retention Time | Super class |
|---|---|---|
| LC.03.0083 | 22.64789474 | Lipids and lipid-like molecules |
| LC.03.0084 | 618.0843158 | Organic oxygen compounds |
| LC.03.0085 | 666.2212632 | Lipids and lipid-like molecules |
| LC.03.0087 | 744.8166842 | Lipids and lipid-like molecules |
| LC.03.0088 | 749.2202632 | Organic oxygen compounds |
| LC.03.0089 | 737.9352632 | Lipids and lipid-like molecules |
| LC.03.0090 | 768.1645789 | Lipids and lipid-like molecules |
| LC.03.0092 | 188.4445263 | NA |
| LC.03.0093 | 170.1322632 | NA |
| LC.03.0094 | 23.52938889 | Organic oxygen compounds |
| LC.03.0095 | 23.52188889 | NA |
| LC.03.0096 | 18.11633333 | Phenylpropanoids and polyketides |
| LC.03.0097 | 618.3228333 | NA |
| LC.03.0098 | 834.5708333 | Lipids and lipid-like molecules |
| LC.03.0099 | 788.1256667 | NA |
| LC.03.0100 | 774.9932778 | Lipids and lipid-like molecules |
| LC.03.0101 | 770.8487778 | NA |
| LC.03.0102 | 474.7701667 | NA |
| LC.03.0103 | 522.9615556 | Lipids and lipid-like molecules |
| LC.03.0104 | 319.1828889 | Lipids and lipid-like molecules |
| LC.03.0105 | 339.8824444 | Organic acids and derivatives |
| LC.03.0106 | 867.3052941 | NA |
| LC.03.0107 | 702.1449412 | Lipids and lipid-like molecules |
| LC.03.0109 | 726.1416471 | Phenylpropanoids and polyketides |
| LC.03.0110 | 737.1485294 | Lipids and lipid-like molecules |
| LC.03.0111 | 761.3968235 | Lipids and lipid-like molecules |
| LC.03.0112 | 476.599 | Lipids and lipid-like molecules |
| LC.03.0113 | 250.8772941 | Lignans, neolignans and related compounds |
| LC.03.0114 | 302.3903529 | Phenylpropanoids and polyketides |
| LC.03.0115 | 59.9169375 | Phenylpropanoids and polyketides |
| LC.03.0116 | 582.3154375 | NA |
| LC.03.0117 | 667.545875 | NA |
| LC.03.0118 | 675.0288125 | Lipids and lipid-like molecules |
| LC.03.0119 | 791.113375 | NA |

| Compound | Retention Time | Super class |
| --- | --- | --- |
| LC.03.0120 | 389.929625 | Lipids and lipid-like molecules |
| LC.03.0121 | 476.9790625 | Lipids and lipid-like molecules |
| LC.03.0122 | 482.27875 | Lipids and lipid-like molecules |
| LC.03.0123 | 200.9040625 | NA |
| LC.03.0124 | 307.28825 | NA |
| LC.03.0126 | 22.8896 | Organic acids and derivatives |
| LC.03.0127 | 23.24886667 | Lipids and lipid-like molecules |
| LC.03.0128 | 604.8748667 | Lipids and lipid-like molecules |
| LC.03.0129 | 823.9682 | NA |
| LC.03.0130 | 823.7772 | Lipids and lipid-like molecules |
| LC.03.0132 | 728.1604 | Lipids and lipid-like molecules |
| LC.03.0133 | 805.7139333 | Lipids and lipid-like molecules |
| LC.03.0134 | 758.4136667 | Organic oxygen compounds |
| LC.03.0135 | 756.7864667 | Hydrocarbons |
| LC.03.0136 | 771.1774667 | Lipids and lipid-like molecules |
| LC.03.0137 | 770.5926667 | Organic oxygen compounds |
| LC.03.0138 | 780.9278667 | NA |
| LC.03.0139 | 474.3232667 | Lipids and lipid-like molecules |
| LC.03.0140 | 550.0128 | Lipids and lipid-like molecules |
| LC.03.0141 | 186.1378667 | Phenylpropanoids and polyketides |
| LC.03.0142 | 630.1544286 | Organic acids and derivatives |
| LC.03.0143 | 878.5515 | Lipids and lipid-like molecules |
| LC.03.0145 | 837.2661429 | Lipids and lipid-like molecules |
| LC.03.0146 | 867.7555714 | Lipids and lipid-like molecules |
| LC.03.0147 | 852.7403571 | NA |
| LC.03.0148 | 724.0943571 | Lipids and lipid-like molecules |
| LC.03.0149 | 735.0747857 | Lipids and lipid-like molecules |
| LC.03.0150 | 759.1358571 | Lipids and lipid-like molecules |
| LC.03.0151 | 756.8590714 | Lipids and lipid-like molecules |
| LC.03.0152 | 769.183 | Lipids and lipid-like molecules |
| LC.03.0153 | 386.1592857 | Lipids and lipid-like molecules |
| LC.03.0154 | 395.2845714 | Lipids and lipid-like molecules |
| LC.03.0155 | 449.8193571 | Lipids and lipid-like molecules |
| LC.03.0156 | 454.2207857 | Alkaloids and derivatives |

| Compound | Retention Time | Super class |
|---|---|---|
| LC.03.0157 | 468.9525 | Lipids and lipid-like molecules |
| LC.03.0158 | 536.6536429 | Lipids and lipid-like molecules |
| LC.03.0159 | 192.8697143 | NA |
| LC.03.0160 | 204.811 | NA |
| LC.03.0161 | 18.19107692 | Organic acids and derivatives |
| LC.03.0162 | 584.9566154 | Lipids and lipid-like molecules |
| LC.03.0163 | 572.924 | Lipids and lipid-like molecules |
| LC.03.0164 | 572.2059231 | Lipids and lipid-like molecules |
| LC.03.0165 | 618.4393846 | Organoheterocyclic compounds |
| LC.03.0167 | 830.4668462 | Lipids and lipid-like molecules |
| LC.03.0169 | 862.1353846 | Lipids and lipid-like molecules |
| LC.03.0170 | 869.2729231 | Lipids and lipid-like molecules |
| LC.03.0171 | 716.7555385 | Lipids and lipid-like molecules |
| LC.03.0172 | 727.6268462 | NA |
| LC.03.0173 | 732.3497692 | Lipids and lipid-like molecules |
| LC.03.0174 | 751.8183077 | Organic oxygen compounds |
| LC.03.0175 | 756.4976923 | NA |
| LC.03.0176 | 493.9808462 | Phenylpropanoids and polyketides |
| LC.03.0177 | 554.8421538 | Organoheterocyclic compounds |
| LC.03.0178 | 186.7520769 | NA |
| LC.03.0179 | 23.56991667 | Organic oxygen compounds |
| LC.03.0180 | 23.19658333 | NA |
| LC.03.0181 | 22.43183333 | Lipids and lipid-like molecules |
| LC.03.0182 | 23.51608333 | Organic oxygen compounds |
| LC.03.0183 | 16.91475 | NA |
| LC.03.0184 | 18.10175 | NA |
| LC.03.0185 | 596.5398333 | Lipids and lipid-like molecules |
| LC.03.0186 | 572.1825833 | Lipids and lipid-like molecules |
| LC.03.0187 | 572.17425 | Lipids and lipid-like molecules |
| LC.03.0189 | 611.8813333 | Organic acids and derivatives |
| LC.03.0190 | 614.5425833 | NA |
| LC.03.0191 | 614.1424167 | Organoheterocyclic compounds |
| LC.03.0192 | 873.58725 | Organic oxygen compounds |
| LC.03.0193 | 844.99725 | Lipids and lipid-like molecules |

| Compound | Retention Time | Super class |
|---|---|---|
| LC.03.0194 | 675.6195833 | Organic oxygen compounds |
| LC.03.0195 | 804.6174167 | NA |
| LC.03.0196 | 762.8588333 | NA |
| LC.03.0197 | 767.4145833 | NA |
| LC.03.0198 | 786.3258333 | Organic oxygen compounds |
| LC.03.0199 | 780.8195 | NA |
| LC.03.0200 | 402.65675 | Lipids and lipid-like molecules |
| LC.03.0201 | 447.5168333 | NA |
| LC.03.0202 | 457.0085833 | Phenylpropanoids and polyketides |
| LC.03.0203 | 517.6128333 | Organic nitrogen compounds |
| LC.03.0204 | 188.7555833 | Organic acids and derivatives |
| LC.03.0205 | 1000.34575 | Lipids and lipid-like molecules |
| LC.03.0206 | 586.9548182 | Organic Polymers |
| LC.03.0207 | 578.4074545 | NA |
| LC.03.0208 | 631.8280909 | Lipids and lipid-like molecules |
| LC.03.0209 | 630.5431818 | Organoheterocyclic compounds |
| LC.03.0210 | 617.3987273 | Lipids and lipid-like molecules |
| LC.03.0211 | 651.9716364 | Lipids and lipid-like molecules |
| LC.03.0212 | 647.063 | Lipids and lipid-like molecules |
| LC.03.0213 | 646.178 | NA |
| LC.03.0215 | 829.3309091 | NA |
| LC.03.0217 | 713.4532727 | Lipids and lipid-like molecules |
| LC.03.0218 | 677.6634545 | Lipids and lipid-like molecules |
| LC.03.0219 | 691.7984545 | NA |
| LC.03.0220 | 724.8190909 | Lipids and lipid-like molecules |
| LC.03.0221 | 727.4273636 | Lipids and lipid-like molecules |
| LC.03.0222 | 738.7501818 | Organic oxygen compounds |
| LC.03.0223 | 735.3040909 | Lipids and lipid-like molecules |
| LC.03.0227 | 544.0159091 | Lipids and lipid-like molecules |
| LC.03.0228 | 186.254 | Organic oxygen compounds |
| LC.03.0229 | 34.6456 | NA |
| LC.03.0230 | 589.5901 | NA |
| LC.03.0232 | 575.3978 | Phenylpropanoids and polyketides |
| LC.03.0233 | 632.0601 | Lipids and lipid-like molecules |

| Compound | Retention Time | Super class |
| --- | --- | --- |
| LC.03.0234 | 666.1024 | Lipids and lipid-like molecules |
| LC.03.0235 | 645.1089 | Organic acids and derivatives |
| LC.03.0236 | 817.5262 | NA |
| LC.03.0237 | 826.7764 | Organic oxygen compounds |
| LC.03.0238 | 834.3775 | Lipids and lipid-like molecules |
| LC.03.0239 | 711.9875 | Lipids and lipid-like molecules |
| LC.03.0240 | 717.5192 | Lipids and lipid-like molecules |
| LC.03.0241 | 685.3476 | Lipids and lipid-like molecules |
| LC.03.0243 | 724.6249 | Lipids and lipid-like molecules |
| LC.03.0244 | 727.8339 | NA |
| LC.03.0245 | 805.9303 | Lipids and lipid-like molecules |
| LC.03.0246 | 799.0817 | NA |
| LC.03.0247 | 791.9629 | Organic oxygen compounds |
| LC.03.0248 | 772.0325 | Lipids and lipid-like molecules |
| LC.03.0249 | 778.0164 | NA |
| LC.03.0250 | 447.2739 | Lipids and lipid-like molecules |
| LC.03.0251 | 488.9638 | Lipids and lipid-like molecules |
| LC.03.0252 | 187.4836 | NA |
| LC.03.0253 | 185.0866 | NA |
| LC.03.0254 | 239.0817 | Organic acids and derivatives |
| LC.03.0255 | 293.6246 | Phenylpropanoids and polyketides |
| LC.03.0256 | 358.0024 | Organoheterocyclic compounds |
| LC.03.0257 | 23.84077778 | Benzenoids |
| LC.03.0258 | 22.92044444 | Organic acids and derivatives |
| LC.03.0259 | 52.77844444 | Phenylpropanoids and polyketides |
| LC.03.0260 | 588.8681111 | Lipids and lipid-like molecules |
| LC.03.0261 | 583.4883333 | Organic acids and derivatives |
| LC.03.0262 | 636.9371111 | Lipids and lipid-like molecules |
| LC.03.0263 | 635.4705556 | Benzenoids |
| LC.03.0264 | 620.1863333 | Lipids and lipid-like molecules |
| LC.03.0265 | 625.0436667 | NA |
| LC.03.0266 | 625.1093333 | Organic acids and derivatives |
| LC.03.0267 | 610.9373333 | Lipids and lipid-like molecules |
| LC.03.0268 | 662.2994444 | NA |

| Compound | Retention Time | Super class |
|---|---|---|
| LC.03.0269 | 667.4981111 | Lipids and lipid-like molecules |
| LC.03.0270 | 876.4415556 | Lipids and lipid-like molecules |
| LC.03.0271 | 881.0627778 | Lipids and lipid-like molecules |
| LC.03.0272 | 891.0606667 | NA |
| LC.03.0273 | 837.526 | Lipids and lipid-like molecules |
| LC.03.0274 | 844.2912222 | NA |
| LC.03.0275 | 847.5341111 | NA |
| LC.03.0276 | 858.8853333 | NA |
| LC.03.0277 | 854.1113333 | NA |
| LC.03.0280 | 725.6763333 | Lipids and lipid-like molecules |
| LC.03.0281 | 740.4885556 | NA |
| LC.03.0282 | 742.6215556 | Lipids and lipid-like molecules |
| LC.03.0283 | 742.3885556 | NA |
| LC.03.0284 | 740.0711111 | NA |
| LC.03.0285 | 737.6944444 | Lipids and lipid-like molecules |
| LC.03.0286 | 798.7516667 | Lipids and lipid-like molecules |
| LC.03.0287 | 761.4001111 | NA |
| LC.03.0288 | 764.9797778 | NA |
| LC.03.0289 | 787.2145556 | Lipids and lipid-like molecules |
| LC.03.0291 | 789.7938889 | NA |
| LC.03.0292 | 772.1188889 | NA |
| LC.03.0293 | 769.198 | Lipids and lipid-like molecules |
| LC.03.0295 | 784.8697778 | NA |
| LC.03.0296 | 778.2196667 | Lipids and lipid-like molecules |
| LC.03.0297 | 778.5271111 | Lipids and lipid-like molecules |
| LC.03.0298 | 401.025 | NA |
| LC.03.0299 | 409.4957778 | NA |
| LC.03.0300 | 409.4 | Lipids and lipid-like molecules |
| LC.03.0301 | 500.1615556 | NA |
| LC.03.0302 | 474.843 | NA |
| LC.03.0303 | 482.8468889 | Lipids and lipid-like molecules |
| LC.03.0304 | 186.721 | NA |
| LC.03.0305 | 188.8614444 | NA |
| LC.03.0306 | 193.4805556 | NA |

| Compound | Retention Time | Super class |
|---|---|---|
| LC.03.0307 | 195.4077778 | NA |
| LC.03.0308 | 178.2732222 | NA |
| LC.03.0309 | 260.7042222 | NA |
| LC.03.0310 | 342.4417778 | Lipids and lipid-like molecules |
| LC.03.0311 | 1003.346 | NA |
| LC.03.0312 | 31.269375 | NA |
| LC.03.0313 | 23.68525 | NA |
| LC.03.0314 | 23.628375 | Lipids and lipid-like molecules |
| LC.03.0315 | 23.086125 | Alkaloids and derivatives |
| LC.03.0316 | 17.063375 | NA |
| LC.03.0317 | 18.045625 | NA |
| LC.03.0318 | 44.0005 | NA |
| LC.03.0319 | 579.70025 | Lipids and lipid-like molecules |
| LC.03.0320 | 578.35275 | NA |
| LC.03.0321 | 629.95125 | NA |
| LC.03.0322 | 617.200875 | NA |
| LC.03.0323 | 623.798875 | NA |
| LC.03.0324 | 604.82875 | NA |
| LC.03.0325 | 610.9585 | NA |
| LC.03.0326 | 659.30625 | NA |
| LC.03.0327 | 653.464125 | NA |
| LC.03.0328 | 645.15775 | NA |
| LC.03.0329 | 641.238625 | NA |
| LC.03.0330 | 819.352125 | NA |
| LC.03.0332 | 828.511875 | NA |
| LC.03.0333 | 823.946375 | NA |
| LC.03.0335 | 842.074125 | NA |
| LC.03.0336 | 833.45425 | Lipids and lipid-like molecules |
| LC.03.0339 | 696.6665 | NA |
| LC.03.0340 | 730.9925 | NA |
| LC.03.0341 | 746.408875 | NA |
| LC.03.0342 | 751.85175 | Lipids and lipid-like molecules |
| LC.03.0343 | 805.675 | NA |
| LC.03.0344 | 764.796875 | Lipids and lipid-like molecules |

| Compound | Retention Time | Super class |
|---|---|---|
| LC.03.0345 | 756.435375 | NA |
| LC.03.0346 | 756.716625 | Lipids and lipid-like molecules |
| LC.03.0347 | 790.313 | NA |
| LC.03.0348 | 770.76975 | NA |
| LC.03.0349 | 772.26325 | NA |
| LC.03.0351 | 781.567125 | Lipids and lipid-like molecules |
| LC.03.0352 | 780.097375 | NA |
| LC.03.0353 | 778.159 | NA |
| LC.03.0354 | 777.33775 | Lipids and lipid-like molecules |
| LC.03.0355 | 777.36775 | NA |
| LC.03.0356 | 380.171375 | NA |
| LC.03.0357 | 397.628125 | Lipids and lipid-like molecules |
| LC.03.0358 | 456.90625 | Lipids and lipid-like molecules |
| LC.03.0359 | 502.271625 | Lipids and lipid-like molecules |
| LC.03.0360 | 467.973375 | NA |
| LC.03.0361 | 551.159125 | Organoheterocyclic compounds |
| LC.03.0362 | 184.440375 | NA |
| LC.03.0363 | 199.23175 | NA |
| LC.03.0364 | 199.83275 | NA |
| LC.03.0365 | 180.8735 | NA |
| LC.03.0369 | 272.71225 | NA |
| LC.03.0370 | 286.82 | Phenylpropanoids and polyketides |
| LC.03.0372 | 347.814625 | NA |
| LC.03.0373 | 22.79942857 | NA |
| LC.03.0374 | 23.14685714 | NA |
| LC.03.0375 | 23.13985714 | Lipids and lipid-like molecules |
| LC.03.0376 | 23.32542857 | NA |
| LC.03.0377 | 22.75528571 | Organoheterocyclic compounds |
| LC.03.0378 | 17.943 | Organic oxygen compounds |
| LC.03.0379 | 46.71114286 | NA |
| LC.03.0380 | 596.7442857 | Lipids and lipid-like molecules |
| LC.03.0381 | 577.3551429 | Lipids and lipid-like molecules |
| LC.03.0382 | 631.6404286 | NA |
| LC.03.0383 | 630.8931429 | Phenylpropanoids and polyketides |

| Compound | Retention Time | Super class |
|---|---|---|
| LC.03.0384 | 631.9091429 | NA |
| LC.03.0385 | 628.1182857 | Lipids and lipid-like molecules |
| LC.03.0386 | 624.766 | NA |
| LC.03.0387 | 600.9855714 | NA |
| LC.03.0388 | 608.69 | NA |
| LC.03.0389 | 664.5968571 | Lipids and lipid-like molecules |
| LC.03.0390 | 848.9338571 | NA |
| LC.03.0391 | 861.9571429 | Lipids and lipid-like molecules |
| LC.03.0392 | 857.6711429 | NA |
| LC.03.0393 | 859.2584286 | Lipids and lipid-like molecules |
| LC.03.0394 | 711.2688571 | NA |
| LC.03.0395 | 712.813 | NA |
| LC.03.0396 | 721.7582857 | Lipids and lipid-like molecules |
| LC.03.0397 | 674.4045714 | Benzenoids |
| LC.03.0398 | 681.2037143 | Lipids and lipid-like molecules |
| LC.03.0400 | 697.0745714 | NA |
| LC.03.0402 | 692.469 | Lipids and lipid-like molecules |
| LC.03.0403 | 726.8544286 | Lipids and lipid-like molecules |
| LC.03.0404 | 746.3941429 | Lipids and lipid-like molecules |
| LC.03.0405 | 746.0428571 | Organic acids and derivatives |
| LC.03.0406 | 735.6632857 | Lipids and lipid-like molecules |
| LC.03.0408 | 796.1501429 | NA |
| LC.03.0409 | 796.7842857 | NA |
| LC.03.0410 | 757.8942857 | Phenylpropanoids and polyketides |
| LC.03.0411 | 755.6172857 | Lipids and lipid-like molecules |
| LC.03.0412 | 770.853 | Lipids and lipid-like molecules |
| LC.03.0413 | 784.2265714 | NA |
| LC.03.0414 | 785.573 | Lipids and lipid-like molecules |
| LC.03.0415 | 780.9108571 | Lipids and lipid-like molecules |
| LC.03.0416 | 779.2102857 | Organoheterocyclic compounds |
| LC.03.0417 | 373.7687143 | Organoheterocyclic compounds |
| LC.03.0418 | 396.1874286 | NA |
| LC.03.0419 | 395.0228571 | NA |
| LC.03.0420 | 404.5392857 | NA |

| Compound | Retention Time | Super class |
|---|---|---|
| LC.03.0421 | 413.1582857 | Lipids and lipid-like molecules |
| LC.03.0422 | 497.0507143 | Lipids and lipid-like molecules |
| LC.03.0423 | 500.34 | Lipids and lipid-like molecules |
| LC.03.0424 | 541.6601429 | Lipids and lipid-like molecules |
| LC.03.0426 | 186.4548571 | NA |
| LC.03.0427 | 186.0041429 | NA |
| LC.03.0428 | 189.554 | NA |
| LC.03.0429 | 193.3357143 | NA |
| LC.03.0430 | 196.2865714 | NA |
| LC.03.0431 | 200.6652857 | NA |
| LC.03.0432 | 197.6215714 | NA |
| LC.03.0434 | 261.365 | NA |
| LC.03.0435 | 229.0274286 | NA |
| LC.03.0436 | 219.3641429 | NA |
| LC.03.0437 | 315.2612857 | Phenylpropanoids and polyketides |
| LC.03.0438 | 298.4435714 | Lipids and lipid-like molecules |
| LC.03.0439 | 335.6967143 | NA |
| LC.03.0440 | 343.3422857 | Lipids and lipid-like molecules |
| LC.03.0441 | 32.142 | Phenylpropanoids and polyketides |
| LC.03.0442 | 34.78916667 | NA |
| LC.03.0443 | 21.70466667 | Phenylpropanoids and polyketides |
| LC.03.0444 | 22.98566667 | NA |
| LC.03.0445 | 22.98466667 | NA |
| LC.03.0446 | 17.04016667 | NA |
| LC.03.0447 | 18.14533333 | Organoheterocyclic compounds |
| LC.03.0448 | 60.046 | NA |
| LC.03.0449 | 44.28066667 | NA |
| LC.03.0450 | 595.7781667 | NA |
| LC.03.0451 | 583.288 | Lipids and lipid-like molecules |
| LC.03.0452 | 577.3768333 | Organoheterocyclic compounds |
| LC.03.0453 | 574.7131667 | NA |
| LC.03.0454 | 632.6543333 | NA |
| LC.03.0456 | 626.937 | NA |
| LC.03.0457 | 623.8405 | NA |

| Compound | Retention Time | Super class |
|---|---|---|
| LC.03.0458 | 623.5641667 | Lipids and lipid-like molecules |
| LC.03.0459 | 623.7191667 | NA |
| LC.03.0460 | 604.8513333 | Lipids and lipid-like molecules |
| LC.03.0461 | 645.6496667 | NA |
| LC.03.0462 | 645.5255 | NA |
| LC.03.0463 | 870.9551667 | NA |
| LC.03.0464 | 896.0383333 | Lipids and lipid-like molecules |
| LC.03.0465 | 821.4775 | Lipids and lipid-like molecules |
| LC.03.0466 | 815.0978333 | NA |
| LC.03.0469 | 848.8095 | NA |
| LC.03.0471 | 866.3981667 | NA |
| LC.03.0472 | 855.2743333 | Lipids and lipid-like molecules |
| LC.03.0473 | 700.7153333 | NA |
| LC.03.0474 | 710.3183333 | Lipids and lipid-like molecules |
| LC.03.0475 | 720.51 | Lipids and lipid-like molecules |
| LC.03.0476 | 717.5561667 | Lipids and lipid-like molecules |
| LC.03.0477 | 718.9123333 | Lipids and lipid-like molecules |
| LC.03.0478 | 674.19 | Lipids and lipid-like molecules |
| LC.03.0479 | 679.9371667 | NA |
| LC.03.0480 | 695.9811667 | NA |
| LC.03.0481 | 692.927 | NA |
| LC.03.0482 | 690.1338333 | NA |
| LC.03.0483 | 722.8616667 | NA |
| LC.03.0484 | 731.6405 | NA |
| LC.03.0485 | 745.6671667 | NA |
| LC.03.0486 | 744.8145 | NA |
| LC.03.0487 | 744.2225 | NA |
| LC.03.0488 | 748.4273333 | Lipids and lipid-like molecules |
| LC.03.0489 | 748.9246667 | Organic oxygen compounds |
| LC.03.0490 | 738.5878333 | Organic oxygen compounds |
| LC.03.0491 | 802.4348333 | NA |
| LC.03.0492 | 800.7626667 | NA |
| LC.03.0493 | 765.187 | NA |
| LC.03.0494 | 764.4735 | NA |

| Compound | Retention Time | Super class |
|---|---|---|
| LC.03.0495 | 790.4328333 | Lipids and lipid-like molecules |
| LC.03.0496 | 787.5468333 | NA |
| LC.03.0497 | 789.3603333 | Lipids and lipid-like molecules |
| LC.03.0498 | 767.7416667 | NA |
| LC.03.0499 | 768.9703333 | Lipids and lipid-like molecules |
| LC.03.0500 | 784.3068333 | Organic oxygen compounds |
| LC.03.0501 | 785.5315 | Lipids and lipid-like molecules |
| LC.03.0502 | 781.832 | Lipids and lipid-like molecules |
| LC.03.0503 | 781.1201667 | Lipids and lipid-like molecules |
| LC.03.0504 | 779.9603333 | Lipids and lipid-like molecules |
| LC.03.0505 | 779.3933333 | Lipids and lipid-like molecules |
| LC.03.0506 | 780.9278333 | Lipids and lipid-like molecules |
| LC.03.0507 | 387.5458333 | Lipids and lipid-like molecules |
| LC.03.0508 | 409.483 | Phenylpropanoids and polyketides |
| LC.03.0509 | 429.8105 | Lipids and lipid-like molecules |
| LC.03.0510 | 444.7783333 | NA |
| LC.03.0511 | 459.3028333 | Lipids and lipid-like molecules |
| LC.03.0512 | 494.0485 | Phenylpropanoids and polyketides |
| LC.03.0513 | 493.433 | NA |
| LC.03.0515 | 479.1803333 | NA |
| LC.03.0516 | 481.7378333 | Lipids and lipid-like molecules |
| LC.03.0517 | 509.58 | NA |
| LC.03.0518 | 523.6296667 | NA |
| LC.03.0519 | 531.8875 | NA |
| LC.03.0520 | 561.1721667 | NA |
| LC.03.0521 | 558.5926667 | Lipids and lipid-like molecules |
| LC.03.0522 | 563.2918333 | Lipids and lipid-like molecules |
| LC.03.0523 | 194.3438333 | NA |
| LC.03.0524 | 178.4958333 | NA |
| LC.03.0525 | 175.4705 | NA |
| LC.03.0526 | 170.0255 | Organoheterocyclic compounds |
| LC.03.0527 | 161.8415 | NA |
| LC.03.0529 | 257.4003333 | Benzenoids |
| LC.03.0530 | 269.1216667 | Phenylpropanoids and polyketides |

| Compound | Retention Time | Super class |
|---|---|---|
| LC.03.0531 | 276.028 | Benzenoids |
| LC.03.0532 | 240.3298333 | NA |
| LC.03.0533 | 246.9325 | NA |
| LC.03.0534 | 311.6365 | Organic acids and derivatives |
| LC.03.0535 | 357.369 | NA |
| LC.03.0536 | 333.5505 | Benzenoids |
| LC.03.0537 | 333.9526667 | Lipids and lipid-like molecules |
| LC.03.0538 | 331.3646667 | Organoheterocyclic compounds |
| LC.03.0539 | 1002.354 | Phenylpropanoids and polyketides |
| LC.03.0540 | 71.0132 | Organoheterocyclic compounds |
| LC.03.0541 | 70.7738 | Organoheterocyclic compounds |
| LC.03.0542 | 32.7344 | NA |
| LC.03.0543 | 33.6744 | Nucleosides, nucleotides, and analogues |
| LC.03.0544 | 23.5142 | Organoheterocyclic compounds |
| LC.03.0545 | 23.589 | NA |
| LC.03.0546 | 25.3556 | NA |
| LC.03.0547 | 23.223 | NA |
| LC.03.0548 | 21.7178 | Benzenoids |
| LC.03.0549 | 52.1256 | NA |
| LC.03.0550 | 46.8744 | Organic acids and derivatives |
| LC.03.0551 | 588.2316 | NA |
| LC.03.0552 | 575.0152 | NA |
| LC.03.0553 | 568.7996 | Lipids and lipid-like molecules |
| LC.03.0555 | 617.4708 | NA |
| LC.03.0556 | 623.815 | NA |
| LC.03.0557 | 668.2492 | NA |
| LC.03.0558 | 649.0706 | NA |
| LC.03.0559 | 883.1052 | NA |
| LC.03.0560 | 881.3402 | NA |
| LC.03.0561 | 882.3758 | Lipids and lipid-like molecules |
| LC.03.0562 | 872.2074 | NA |
| LC.03.0564 | 901.0276 | Organic oxygen compounds |
| LC.03.0565 | 815.6816 | NA |
| LC.03.0566 | 817.3788 | NA |

| Compound | Retention Time | Super class |
|---|---|---|
| LC.03.0567 | 814.2602 | Lipids and lipid-like molecules |
| LC.03.0569 | 842.9766 | NA |
| LC.03.0570 | 846.5042 | NA |
| LC.03.0571 | 849.53 | NA |
| LC.03.0573 | 860.5482 | Organic oxygen compounds |
| LC.03.0574 | 868.9736 | Lipids and lipid-like molecules |
| LC.03.0575 | 857.815 | NA |
| LC.03.0576 | 856.7972 | NA |
| LC.03.0577 | 859.3522 | NA |
| LC.03.0578 | 852.7774 | NA |
| LC.03.0579 | 851.9562 | NA |
| LC.03.0580 | 704.5222 | NA |
| LC.03.0581 | 716.9544 | NA |
| LC.03.0582 | 718.3982 | NA |
| LC.03.0583 | 681.6978 | Lipids and lipid-like molecules |
| LC.03.0585 | 683.4006 | NA |
| LC.03.0586 | 698.133 | NA |
| LC.03.0587 | 694.997 | Lipids and lipid-like molecules |
| LC.03.0589 | 725.2816 | Lipids and lipid-like molecules |
| LC.03.0590 | 732.664 | NA |
| LC.03.0591 | 746.8584 | Lipids and lipid-like molecules |
| LC.03.0592 | 751.517 | Lipids and lipid-like molecules |
| LC.03.0593 | 750.2308 | NA |
| LC.03.0594 | 738.1194 | Organic acids and derivatives |
| LC.03.0595 | 796.4674 | NA |
| LC.03.0596 | 799.2546 | Lipids and lipid-like molecules |
| LC.03.0597 | 755.2304 | NA |
| LC.03.0598 | 788.9484 | Lipids and lipid-like molecules |
| LC.03.0599 | 789.6946 | NA |
| LC.03.0600 | 791.474 | Lipids and lipid-like molecules |
| LC.03.0601 | 793.116 | NA |
| LC.03.0602 | 774.2984 | NA |
| LC.03.0603 | 782.634 | Lipids and lipid-like molecules |
| LC.03.0604 | 382.7472 | NA |

| Compound | Retention Time | Super class |
|---|---|---|
| LC.03.0605 | 388.1226 | NA |
| LC.03.0606 | 372.9346 | NA |
| LC.03.0607 | 396.643 | NA |
| LC.03.0608 | 394.1066 | Organoheterocyclic compounds |
| LC.03.0609 | 402.1658 | NA |
| LC.03.0610 | 438.1418 | NA |
| LC.03.0611 | 446.3924 | NA |
| LC.03.0612 | 444.1152 | Lipids and lipid-like molecules |
| LC.03.0613 | 456.3678 | Organic acids and derivatives |
| LC.03.0614 | 486.3364 | NA |
| LC.03.0615 | 490.3114 | Organoheterocyclic compounds |
| LC.03.0616 | 488.1836 | NA |
| LC.03.0617 | 467.128 | Lipids and lipid-like molecules |
| LC.03.0618 | 467.8018 | NA |
| LC.03.0619 | 473.7856 | NA |
| LC.03.0621 | 480.5684 | NA |
| LC.03.0622 | 508.581 | NA |
| LC.03.0623 | 545 | Lipids and lipid-like molecules |
| LC.03.0624 | 533.911 | Lipids and lipid-like molecules |
| LC.03.0625 | 532.4618 | Lipids and lipid-like molecules |
| LC.03.0626 | 555.338 | NA |
| LC.03.0627 | 551.055 | Alkaloids and derivatives |
| LC.03.0628 | 187.5498 | NA |
| LC.03.0629 | 185.7932 | NA |
| LC.03.0630 | 185.646 | NA |
| LC.03.0631 | 190.4132 | NA |
| LC.03.0632 | 200.5026 | NA |
| LC.03.0633 | 202.6474 | NA |
| LC.03.0634 | 179.0494 | NA |
| LC.03.0635 | 170.8746 | NA |
| LC.03.0636 | 268.9506 | NA |
| LC.03.0637 | 226.9856 | Benzenoids |
| LC.03.0638 | 217.8296 | NA |
| LC.03.0639 | 235.098 | NA |

| Compound | Retention Time | Super class |
|---|---|---|
| LC.03.0640 | 291.7698 | Phenylpropanoids and polyketides |
| LC.03.0641 | 292.1006 | NA |
| LC.03.0642 | 349.0844 | Organoheterocyclic compounds |
| LC.03.0643 | 339.354 | NA |
| LC.03.0644 | 1001.6088 | NA |
| LC.03.0645 | 1000.3658 | NA |
| LC.03.0646 | 36.74925 | NA |
| LC.03.0647 | 25.151 | Benzenoids |
| LC.03.0648 | 26.99225 | Organic acids and derivatives |
| LC.03.0649 | 20.6485 | Organoheterocyclic compounds |
| LC.03.0650 | 23.3565 | NA |
| LC.03.0651 | 23.39425 | NA |
| LC.03.0652 | 22.884 | NA |
| LC.03.0653 | 22.60875 | Organic acids and derivatives |
| LC.03.0654 | 22.64025 | NA |
| LC.03.0655 | 23.60125 | NA |
| LC.03.0656 | 18.57275 | Organic nitrogen compounds |
| LC.03.0657 | 20.92475 | NA |
| LC.03.0658 | 45.06525 | NA |
| LC.03.0659 | 910.24525 | NA |
| LC.03.0660 | 915.9385 | Organic oxygen compounds |
| LC.03.0661 | 916.394 | NA |
| LC.03.0662 | 594.7685 | Lipids and lipid-like molecules |
| LC.03.0663 | 587.92825 | NA |
| LC.03.0664 | 574.606 | Lipids and lipid-like molecules |
| LC.03.0665 | 572.772 | NA |
| LC.03.0666 | 618.418 | Phenylpropanoids and polyketides |
| LC.03.0667 | 618.06475 | Lipids and lipid-like molecules |
| LC.03.0669 | 666.8545 | NA |
| LC.03.0670 | 664.69075 | NA |
| LC.03.0671 | 652.0005 | Lipids and lipid-like molecules |
| LC.03.0672 | 890.0625 | Lipids and lipid-like molecules |
| LC.03.0673 | 895.731 | NA |
| LC.03.0675 | 818.897 | Lipids and lipid-like molecules |

| Compound | Retention Time | Super class |
|---|---|---|
| LC.03.0676 | 818.78225 | NA |
| LC.03.0677 | 821.4725 | NA |
| LC.03.0678 | 819.73925 | Lipids and lipid-like molecules |
| LC.03.0679 | 820.60325 | Lipids and lipid-like molecules |
| LC.03.0680 | 813.4305 | NA |
| LC.03.0682 | 827.429 | Lipids and lipid-like molecules |
| LC.03.0684 | 826.81125 | NA |
| LC.03.0685 | 829.36475 | NA |
| LC.03.0688 | 840.9385 | NA |
| LC.03.0689 | 836.59 | NA |
| LC.03.0690 | 836.316 | Lipids and lipid-like molecules |
| LC.03.0691 | 849.03875 | NA |
| LC.03.0692 | 846.6175 | NA |
| LC.03.0699 | 856.6255 | NA |
| LC.03.0700 | 852.8615 | Lipids and lipid-like molecules |
| LC.03.0701 | 854.6445 | NA |
| LC.03.0703 | 703.11775 | Lipids and lipid-like molecules |
| LC.03.0704 | 704.6665 | Lipids and lipid-like molecules |
| LC.03.0705 | 714.4545 | NA |
| LC.03.0706 | 708.07375 | NA |
| LC.03.0707 | 681.17075 | NA |
| LC.03.0710 | 695.799 | NA |
| LC.03.0711 | 695.96425 | Lipids and lipid-like molecules |
| LC.03.0712 | 689.43725 | Lipids and lipid-like molecules |
| LC.03.0713 | 687.1315 | NA |
| LC.03.0715 | 731.0715 | NA |
| LC.03.0716 | 731.852 | Lipids and lipid-like molecules |
| LC.03.0717 | 741.95875 | NA |
| LC.03.0718 | 807.289 | NA |
| LC.03.0719 | 804.0045 | Lipids and lipid-like molecules |
| LC.03.0720 | 807.5665 | NA |
| LC.03.0721 | 796.05675 | NA |
| LC.03.0722 | 801.93475 | NA |
| LC.03.0723 | 798.68475 | NA |

| Compound | Retention Time | Super class |
|---|---|---|
| LC.03.0724 | 797.99 | NA |
| LC.03.0726 | 763.931 | NA |
| LC.03.0727 | 764.292 | NA |
| LC.03.0729 | 758.3795 | Lipids and lipid-like molecules |
| LC.03.0731 | 794.6735 | Lipids and lipid-like molecules |
| LC.03.0733 | 792.84125 | NA |
| LC.03.0735 | 771.36175 | Lipids and lipid-like molecules |
| LC.03.0736 | 769.40525 | Lipids and lipid-like molecules |
| LC.03.0737 | 785.6155 | NA |
| LC.03.0738 | 784.18475 | Organoheterocyclic compounds |
| LC.03.0739 | 784.632 | NA |
| LC.03.0740 | 783.08175 | NA |
| LC.03.0741 | 783.249 | NA |
| LC.03.0742 | 783.004 | Lipids and lipid-like molecules |
| LC.03.0743 | 780.70525 | Lipids and lipid-like molecules |
| LC.03.0744 | 781.16775 | Lipids and lipid-like molecules |
| LC.03.0745 | 781.58325 | NA |
| LC.03.0747 | 776.23 | NA |
| LC.03.0748 | 776.16775 | NA |
| LC.03.0749 | 386.251 | NA |
| LC.03.0750 | 380.299 | NA |
| LC.03.0751 | 436.605 | NA |
| LC.03.0752 | 430.3315 | Phenylpropanoids and polyketides |
| LC.03.0753 | 441.33325 | NA |
| LC.03.0754 | 452.07675 | NA |
| LC.03.0755 | 491.713 | NA |
| LC.03.0756 | 503.49675 | NA |
| LC.03.0757 | 497.13825 | Organosulfur compounds |
| LC.03.0758 | 496.22625 | Lipids and lipid-like molecules |
| LC.03.0759 | 464.767 | Lipids and lipid-like molecules |
| LC.03.0760 | 466.53175 | Organic acids and derivatives |
| LC.03.0761 | 468.223 | NA |
| LC.03.0763 | 474.64 | NA |
| LC.03.0764 | 483.3625 | NA |

| Compound | Retention Time | Super class |
|---|---|---|
| LC.03.0765 | 481.10175 | NA |
| LC.03.0766 | 507.1525 | Lipids and lipid-like molecules |
| LC.03.0767 | 527.91425 | Lipids and lipid-like molecules |
| LC.03.0768 | 518.288 | NA |
| LC.03.0769 | 515.4305 | NA |
| LC.03.0770 | 545.04175 | NA |
| LC.03.0771 | 542.52775 | Organic nitrogen compounds |
| LC.03.0772 | 533.27375 | Lipids and lipid-like molecules |
| LC.03.0773 | 555.11625 | NA |
| LC.03.0774 | 188.82575 | NA |
| LC.03.0775 | 190.82975 | NA |
| LC.03.0776 | 183.00325 | Organic oxygen compounds |
| LC.03.0777 | 196.9085 | NA |
| LC.03.0778 | 195.75125 | NA |
| LC.03.0779 | 199.632 | NA |
| LC.03.0780 | 204.1685 | Phenylpropanoids and polyketides |
| LC.03.0781 | 172.03625 | NA |
| LC.03.0782 | 173.3625 | NA |
| LC.03.0783 | 173.3055 | NA |
| LC.03.0784 | 158.83275 | NA |
| LC.03.0785 | 262.905 | NA |
| LC.03.0786 | 209.588 | NA |
| LC.03.0787 | 212.72075 | NA |
| LC.03.0788 | 232.749 | NA |
| LC.03.0789 | 247.705 | NA |
| LC.03.0790 | 248.1525 | NA |
| LC.03.0791 | 285.29625 | NA |
| LC.03.0792 | 325.7125 | NA |
| LC.03.0793 | 318.903 | NA |
| LC.03.0794 | 322.2685 | NA |
| LC.03.0795 | 359.8215 | NA |
| LC.03.0796 | 358.236 | Phenylpropanoids and polyketides |
| LC.03.0797 | 357.68325 | NA |
| LC.03.0798 | 355.8645 | NA |

| Compound | Retention Time | Super class |
|---|---|---|
| LC.03.0799 | 341.9315 | NA |
| LC.03.0800 | 29.20033333 | Organoheterocyclic compounds |
| LC.03.0802 | 38.388 | NA |
| LC.03.0803 | 22.476 | NA |
| LC.03.0804 | 23.244 | Lipids and lipid-like molecules |
| LC.03.0805 | 22.74366667 | NA |
| LC.03.0807 | 25.69366667 | NA |
| LC.03.0808 | 15.82533333 | Organic nitrogen compounds |
| LC.03.0809 | 20.841 | NA |
| LC.03.0810 | 51.952 | Organoheterocyclic compounds |
| LC.03.0811 | 47.90366667 | Organoheterocyclic compounds |
| LC.03.0812 | 939.398 | NA |
| LC.03.0813 | 597.176 | NA |
| LC.03.0814 | 592.7416667 | NA |
| LC.03.0815 | 598.4526667 | NA |
| LC.03.0816 | 596.6913333 | Lipids and lipid-like molecules |
| LC.03.0817 | 582.369 | NA |
| LC.03.0819 | 635.6103333 | Organoheterocyclic compounds |
| LC.03.0820 | 621.5803333 | NA |
| LC.03.0821 | 600.3233333 | NA |
| LC.03.0822 | 611.7276667 | NA |
| LC.03.0824 | 670.207 | NA |
| LC.03.0825 | 667.2703333 | Organic acids and derivatives |
| LC.03.0826 | 665.2546667 | Lipids and lipid-like molecules |
| LC.03.0827 | 654.3936667 | NA |
| LC.03.0828 | 653.9963333 | Lipids and lipid-like molecules |
| LC.03.0829 | 654.5133333 | Organic nitrogen compounds |
| LC.03.0830 | 880.5906667 | NA |
| LC.03.0831 | 885.658 | NA |
| LC.03.0832 | 902.959 | NA |
| LC.03.0835 | 839.4 | Lipids and lipid-like molecules |
| LC.03.0837 | 843.5086667 | Lipids and lipid-like molecules |
| LC.03.0838 | 832.0226667 | NA |
| LC.03.0839 | 844.1853333 | NA |

| Compound | Retention Time | Super class |
|---|---|---|
| LC.03.0841 | 865.908 | NA |
| LC.03.0843 | 850.8576667 | Lipids and lipid-like molecules |
| LC.03.0844 | 702.0336667 | Lipids and lipid-like molecules |
| LC.03.0845 | 701.864 | NA |
| LC.03.0846 | 700.8206667 | Lipids and lipid-like molecules |
| LC.03.0847 | 706.5383333 | NA |
| LC.03.0848 | 704.296 | Lipids and lipid-like molecules |
| LC.03.0851 | 713.0363333 | NA |
| LC.03.0852 | 709.203 | NA |
| LC.03.0853 | 678.146 | NA |
| LC.03.0855 | 683.4766667 | NA |
| LC.03.0856 | 697.5783333 | NA |
| LC.03.0857 | 696.1626667 | NA |
| LC.03.0858 | 724.6746667 | NA |
| LC.03.0859 | 746.0456667 | NA |
| LC.03.0860 | 752.4106667 | NA |
| LC.03.0861 | 738.0886667 | NA |
| LC.03.0862 | 737.2293333 | Lipids and lipid-like molecules |
| LC.03.0864 | 806.9423333 | Lipids and lipid-like molecules |
| LC.03.0865 | 799.6286667 | NA |
| LC.03.0866 | 765.8376667 | Lipids and lipid-like molecules |
| LC.03.0867 | 764.788 | NA |
| LC.03.0868 | 754.721 | Lipids and lipid-like molecules |
| LC.03.0869 | 793.18 | NA |
| LC.03.0871 | 789.5896667 | NA |
| LC.03.0873 | 791.66 | Lipids and lipid-like molecules |
| LC.03.0874 | 767.6676667 | Lipids and lipid-like molecules |
| LC.03.0875 | 778.2546667 | Organic oxygen compounds |
| LC.03.0876 | 397.3666667 | NA |
| LC.03.0877 | 403.396 | NA |
| LC.03.0879 | 420.6233333 | NA |
| LC.03.0880 | 434.657 | Organic nitrogen compounds |
| LC.03.0882 | 447.7456667 | NA |
| LC.03.0883 | 441.0543333 | NA |

| Compound | Retention Time | Super class |
|---|---|---|
| LC.03.0884 | 453.5506667 | NA |
| LC.03.0885 | 457.691 | NA |
| LC.03.0887 | 504.0563333 | Lipids and lipid-like molecules |
| LC.03.0888 | 464.4466667 | Lipids and lipid-like molecules |
| LC.03.0889 | 468.8346667 | Organoheterocyclic compounds |
| LC.03.0890 | 476.6913333 | NA |
| LC.03.0891 | 472.991 | Lipids and lipid-like molecules |
| LC.03.0893 | 527.4226667 | NA |
| LC.03.0895 | 513.3593333 | NA |
| LC.03.0896 | 520.702 | Organic acids and derivatives |
| LC.03.0898 | 542.8566667 | Lipids and lipid-like molecules |
| LC.03.0899 | 537.4096667 | NA |
| LC.03.0900 | 536.7043333 | NA |
| LC.03.0901 | 553.9076667 | NA |
| LC.03.0903 | 188.2733333 | NA |
| LC.03.0904 | 183.0856667 | NA |
| LC.03.0906 | 194.424 | NA |
| LC.03.0907 | 195.6446667 | Organoheterocyclic compounds |
| LC.03.0908 | 190.2623333 | NA |
| LC.03.0909 | 192.8403333 | Lipids and lipid-like molecules |
| LC.03.0910 | 200.6213333 | NA |
| LC.03.0911 | 199.1 | Organic acids and derivatives |
| LC.03.0913 | 206.1103333 | Organoheterocyclic compounds |
| LC.03.0914 | 152.6073333 | NA |
| LC.03.0915 | 175.4146667 | NA |
| LC.03.0916 | 168.749 | NA |
| LC.03.0917 | 169.5416667 | Phenylpropanoids and polyketides |
| LC.03.0918 | 172.9333333 | NA |
| LC.03.0919 | 166.023 | NA |
| LC.03.0921 | 162.2853333 | Benzenoids |
| LC.03.0922 | 262.89 | NA |
| LC.03.0923 | 279.3156667 | NA |
| LC.03.0925 | 276.969 | NA |
| LC.03.0926 | 223.5236667 | Phenylpropanoids and polyketides |

| Compound | Retention Time | Super class |
|---|---|---|
| LC.03.0928 | 210.005 | NA |
| LC.03.0929 | 216.242 | NA |
| LC.03.0930 | 216.8366667 | NA |
| LC.03.0931 | 291.8236667 | Phenylpropanoids and polyketides |
| LC.03.0932 | 314.179 | NA |
| LC.03.0933 | 317.783 | Lipids and lipid-like molecules |
| LC.03.0934 | 301.6493333 | Lipids and lipid-like molecules |
| LC.03.0935 | 334.0303333 | Lipids and lipid-like molecules |
| LC.03.0936 | 348.1033333 | NA |
| LC.03.0937 | 336.8936667 | Lipids and lipid-like molecules |
| LC.03.0939 | 1005.009333 | NA |
| LC.03.0940 | 1005.115333 | NA |
| LC.03.0941 | 87.8145 | NA |
| LC.03.0942 | 30.4685 | Lipids and lipid-like molecules |
| LC.03.0943 | 30.5075 | Organic acids and derivatives |
| LC.03.0944 | 25.8475 | Organoheterocyclic compounds |
| LC.03.0945 | 21.355 | Organic acids and derivatives |
| LC.03.0946 | 22.587 | Organic acids and derivatives |
| LC.03.0948 | 21.666 | Alkaloids and derivatives |
| LC.03.0950 | 23.2375 | NA |
| LC.03.0951 | 23.4905 | Organic oxygen compounds |
| LC.03.0952 | 23.727 | Organic acids and derivatives |
| LC.03.0953 | 21.776 | NA |
| LC.03.0954 | 23.127 | NA |
| LC.03.0955 | 23.159 | NA |
| LC.03.0956 | 23.488 | NA |
| LC.03.0957 | 23.197 | Benzenoids |
| LC.03.0958 | 23.4935 | NA |
| LC.03.0960 | 23.549 | NA |
| LC.03.0961 | 23.3635 | NA |
| LC.03.0962 | 23.625 | NA |
| LC.03.0963 | 23.604 | NA |
| LC.03.0964 | 23.5915 | NA |
| LC.03.0965 | 22.776 | Lipids and lipid-like molecules |

| Compound | Retention Time | Super class |
|---|---|---|
| LC.03.0966 | 15.773 | Organic oxygen compounds |
| LC.03.0967 | 17.9655 | NA |
| LC.03.0968 | 19.952 | NA |
| LC.03.0969 | 20.343 | Organic oxygen compounds |
| LC.03.0970 | 60.2475 | Organosulfur compounds |
| LC.03.0971 | 60.45 | NA |
| LC.03.0972 | 51.2545 | Organic oxygen compounds |
| LC.03.0973 | 50.6215 | NA |
| LC.03.0974 | 50.9185 | NA |
| LC.03.0975 | 47.123 | NA |
| LC.03.0976 | 43.055 | NA |
| LC.03.0977 | 44.047 | NA |
| LC.03.0978 | 960.656 | Organic acids and derivatives |
| LC.03.0979 | 932.5805 | NA |
| LC.03.0980 | 595.092 | NA |
| LC.03.0981 | 588.077 | Alkaloids and derivatives |
| LC.03.0982 | 585.4445 | NA |
| LC.03.0983 | 583.235 | NA |
| LC.03.0984 | 579.135 | Lipids and lipid-like molecules |
| LC.03.0987 | 578.3475 | NA |
| LC.03.0988 | 574.6485 | NA |
| LC.03.0989 | 573.823 | NA |
| LC.03.0990 | 568.549 | NA |
| LC.03.0993 | 636.085 | NA |
| LC.03.0994 | 632.076 | NA |
| LC.03.0995 | 633.92 | NA |
| LC.03.0996 | 630.461 | NA |
| LC.03.0997 | 631.955 | NA |
| LC.03.0998 | 628.0945 | Lipids and lipid-like molecules |
| LC.03.0999 | 615.298 | NA |
| LC.03.1000 | 620.8625 | NA |
| LC.03.1001 | 620.2925 | NA |
| LC.03.1002 | 626.7115 | NA |
| LC.03.1003 | 624.8945 | Lipids and lipid-like molecules |

| Compound | Retention Time | Super class |
|---|---|---|
| LC.03.1004 | 606.8615 | NA |
| LC.03.1005 | 612.1665 | Lipids and lipid-like molecules |
| LC.03.1006 | 610.2285 | Organoheterocyclic compounds |
| LC.03.1007 | 659.4325 | NA |
| LC.03.1008 | 658.3675 | Lipids and lipid-like molecules |
| LC.03.1009 | 654.331 | NA |
| LC.03.1011 | 648.772 | Lipids and lipid-like molecules |
| LC.03.1012 | 884.8855 | NA |
| LC.03.1013 | 904.425 | NA |
| LC.03.1014 | 901.3335 | NA |
| LC.03.1015 | 818.45 | NA |
| LC.03.1017 | 849.408 | NA |
| LC.03.1020 | 866.262 | Lipids and lipid-like molecules |
| LC.03.1024 | 704.296 | Lipids and lipid-like molecules |
| LC.03.1025 | 698.941 | NA |
| LC.03.1026 | 699.5205 | Lipids and lipid-like molecules |
| LC.03.1027 | 701.2495 | Lipids and lipid-like molecules |
| LC.03.1028 | 712.2095 | Lipids and lipid-like molecules |
| LC.03.1029 | 714.617 | NA |
| LC.03.1030 | 714.54 | Lipids and lipid-like molecules |
| LC.03.1031 | 713.3365 | NA |
| LC.03.1032 | 706.3025 | Lipids and lipid-like molecules |
| LC.03.1033 | 708.1225 | NA |
| LC.03.1034 | 715.1735 | NA |
| LC.03.1035 | 716.5895 | NA |
| LC.03.1036 | 719.108 | NA |
| LC.03.1037 | 718.661 | NA |
| LC.03.1038 | 675.865 | NA |
| LC.03.1039 | 675.1325 | NA |
| LC.03.1040 | 675.802 | Phenylpropanoids and polyketides |
| LC.03.1041 | 679.296 | NA |
| LC.03.1042 | 680.976 | NA |
| LC.03.1043 | 682.1705 | Lipids and lipid-like molecules |
| LC.03.1044 | 679.2565 | NA |

| Compound | Retention Time | Super class |
|---|---|---|
| LC.03.1045 | 682.962 | NA |
| LC.03.1046 | 684.125 | NA |
| LC.03.1047 | 686.423 | Lipids and lipid-like molecules |
| LC.03.1048 | 697.518 | NA |
| LC.03.1049 | 697.678 | Benzenoids |
| LC.03.1050 | 695.439 | Organic oxygen compounds |
| LC.03.1051 | 687.247 | NA |
| LC.03.1052 | 690.985 | NA |
| LC.03.1054 | 693.5605 | NA |
| LC.03.1055 | 724.5825 | NA |
| LC.03.1056 | 722.2435 | NA |
| LC.03.1057 | 723.1745 | NA |
| LC.03.1058 | 729.526 | NA |
| LC.03.1059 | 727.241 | NA |
| LC.03.1061 | 754.6835 | Lipids and lipid-like molecules |
| LC.03.1062 | 753.267 | Lipids and lipid-like molecules |
| LC.03.1063 | 743.3815 | Lipids and lipid-like molecules |
| LC.03.1064 | 741.4755 | Lipids and lipid-like molecules |
| LC.03.1067 | 737.257 | NA |
| LC.03.1068 | 737.5755 | NA |
| LC.03.1069 | 738.0475 | NA |
| LC.03.1070 | 738.2635 | Lipids and lipid-like molecules |
| LC.03.1072 | 808.806 | Organoheterocyclic compounds |
| LC.03.1073 | 801.1915 | NA |
| LC.03.1074 | 800.6985 | NA |
| LC.03.1075 | 799.5075 | NA |
| LC.03.1077 | 762.9835 | NA |
| LC.03.1078 | 763.957 | NA |
| LC.03.1080 | 763.9875 | NA |
| LC.03.1081 | 766.304 | NA |
| LC.03.1082 | 768.338 | NA |
| LC.03.1083 | 759.5585 | NA |
| LC.03.1084 | 758.181 | NA |
| LC.03.1085 | 756.71 | NA |

| Compound | Retention Time | Super class |
|---|---|---|
| LC.03.1088 | 789.8765 | Lipids and lipid-like molecules |
| LC.03.1089 | 788.3565 | NA |
| LC.03.1091 | 774.8045 | NA |
| LC.03.1092 | 773.4965 | NA |
| LC.03.1093 | 768.6365 | NA |
| LC.03.1095 | 768.7545 | Lipids and lipid-like molecules |
| LC.03.1096 | 787.108 | NA |
| LC.03.1097 | 784.7615 | NA |
| LC.03.1098 | 782.7555 | Lipids and lipid-like molecules |
| LC.03.1099 | 783.193 | Lipids and lipid-like molecules |
| LC.03.1100 | 776.4685 | NA |
| LC.03.1101 | 779.817 | Lipids and lipid-like molecules |
| LC.03.1102 | 781.38 | NA |
| LC.03.1104 | 775.7515 | Lipids and lipid-like molecules |
| LC.03.1105 | 382.6835 | Lipids and lipid-like molecules |
| LC.03.1106 | 387.0005 | NA |
| LC.03.1107 | 373.8905 | NA |
| LC.03.1108 | 374.1725 | NA |
| LC.03.1109 | 370.0775 | Lipids and lipid-like molecules |
| LC.03.1110 | 379.9695 | NA |
| LC.03.1111 | 379.4375 | Lipids and lipid-like molecules |
| LC.03.1112 | 376.5455 | NA |
| LC.03.1113 | 393.3525 | NA |
| LC.03.1114 | 396.628 | NA |
| LC.03.1115 | 397.3265 | Lipids and lipid-like molecules |
| LC.03.1117 | 398.876 | Organic acids and derivatives |
| LC.03.1118 | 398.865 | Lipids and lipid-like molecules |
| LC.03.1119 | 401.5995 | Benzenoids |
| LC.03.1120 | 402.1595 | Lipids and lipid-like molecules |
| LC.03.1121 | 403.427 | NA |
| LC.03.1122 | 405.708 | NA |
| LC.03.1123 | 409.453 | NA |
| LC.03.1124 | 408.466 | NA |
| LC.03.1125 | 409.388 | NA |

| Compound | Retention Time | Super class |
|---|---|---|
| LC.03.1126 | 408.0895 | Lipids and lipid-like molecules |
| LC.03.1127 | 418.1325 | NA |
| LC.03.1128 | 437.623 | Organic oxygen compounds |
| LC.03.1129 | 436.6755 | NA |
| LC.03.1131 | 446.842 | NA |
| LC.03.1132 | 450.8115 | NA |
| LC.03.1134 | 459.2015 | Lipids and lipid-like molecules |
| LC.03.1135 | 494.8975 | Organic nitrogen compounds |
| LC.03.1136 | 490.564 | Lipids and lipid-like molecules |
| LC.03.1137 | 500.7425 | NA |
| LC.03.1138 | 502.1015 | NA |
| LC.03.1139 | 498.7805 | NA |
| LC.03.1140 | 495.7435 | NA |
| LC.03.1141 | 499.441 | NA |
| LC.03.1142 | 464.966 | Lipids and lipid-like molecules |
| LC.03.1143 | 464.7515 | NA |
| LC.03.1144 | 465.8115 | NA |
| LC.03.1145 | 468.91 | Benzenoids |
| LC.03.1146 | 468.734 | NA |
| LC.03.1147 | 477.128 | Lipids and lipid-like molecules |
| LC.03.1149 | 472.3785 | NA |
| LC.03.1150 | 473.3805 | NA |
| LC.03.1151 | 473.9825 | NA |
| LC.03.1152 | 484.5315 | NA |
| LC.03.1153 | 484.812 | NA |
| LC.03.1154 | 482.7455 | NA |
| LC.03.1155 | 479.96 | Lipids and lipid-like molecules |
| LC.03.1156 | 481.5715 | Lipids and lipid-like molecules |
| LC.03.1157 | 480.912 | Phenylpropanoids and polyketides |
| LC.03.1158 | 481.5995 | NA |
| LC.03.1159 | 526.109 | Organic oxygen compounds |
| LC.03.1160 | 521.448 | Organic oxygen compounds |
| LC.03.1161 | 522.685 | NA |
| LC.03.1162 | 522.3185 | NA |

| Compound | Retention Time | Super class |
|---|---|---|
| LC.03.1163 | 542.7035 | NA |
| LC.03.1164 | 537.8945 | NA |
| LC.03.1165 | 560.0515 | NA |
| LC.03.1166 | 562.251 | NA |
| LC.03.1167 | 550.838 | Organoheterocyclic compounds |
| LC.03.1168 | 549.9465 | NA |
| LC.03.1170 | 184.6535 | NA |
| LC.03.1171 | 194.419 | NA |
| LC.03.1173 | 191.5155 | Lipids and lipid-like molecules |
| LC.03.1174 | 201.1505 | NA |
| LC.03.1175 | 204.392 | NA |
| LC.03.1176 | 143.5615 | Organoheterocyclic compounds |
| LC.03.1177 | 179.9045 | NA |
| LC.03.1178 | 179.8385 | NA |
| LC.03.1179 | 174.2925 | NA |
| LC.03.1180 | 177.136 | Phenylpropanoids and polyketides |
| LC.03.1181 | 176.0565 | NA |
| LC.03.1183 | 168.849 | Lipids and lipid-like molecules |
| LC.03.1184 | 159.6255 | Phenylpropanoids and polyketides |
| LC.03.1185 | 261.0755 | NA |
| LC.03.1186 | 272.1685 | NA |
| LC.03.1187 | 271.541 | Phenylpropanoids and polyketides |
| LC.03.1188 | 225.381 | NA |
| LC.03.1189 | 226.758 | NA |
| LC.03.1190 | 207.137 | NA |
| LC.03.1191 | 209.72 | Organoheterocyclic compounds |
| LC.03.1192 | 210.135 | NA |
| LC.03.1193 | 211.5015 | NA |
| LC.03.1194 | 211.052 | Phenylpropanoids and polyketides |
| LC.03.1195 | 218.947 | NA |
| LC.03.1196 | 219.192 | NA |
| LC.03.1197 | 217.202 | NA |
| LC.03.1198 | 241.5335 | NA |
| LC.03.1199 | 238.747 | NA |

| Compound | Retention Time | Super class |
|---|---|---|
| LC.03.1200 | 236.442 | Lipids and lipid-like molecules |
| LC.03.1201 | 234.4845 | Phenylpropanoids and polyketides |
| LC.03.1202 | 251.2975 | NA |
| LC.03.1205 | 283.7015 | NA |
| LC.03.1206 | 287.576 | Organoheterocyclic compounds |
| LC.03.1207 | 328.0685 | Organic oxygen compounds |
| LC.03.1208 | 327.295 | NA |
| LC.03.1209 | 318.109 | Organoheterocyclic compounds |
| LC.03.1210 | 321.9625 | NA |
| LC.03.1211 | 323.8605 | Phenylpropanoids and polyketides |
| LC.03.1212 | 307.442 | Alkaloids and derivatives |
| LC.03.1213 | 304.673 | NA |
| LC.03.1214 | 307.0595 | NA |
| LC.03.1215 | 309.7015 | Benzenoids |
| LC.03.1216 | 296.4415 | NA |
| LC.03.1217 | 298.375 | NA |
| LC.03.1219 | 300.225 | NA |
| LC.03.1221 | 357.251 | NA |
| LC.03.1222 | 355.1905 | NA |
| LC.03.1223 | 362.9445 | NA |
| LC.03.1224 | 352.9475 | NA |
| LC.03.1225 | 338.739 | NA |
| LC.03.1226 | 336.582 | NA |
| LC.03.1227 | 340.461 | Organic nitrogen compounds |
| LC.03.1228 | 342.3705 | Hydrocarbons |
| LC.03.1229 | 1005.149 | Organic acids and derivatives |
| LC.03.1231 | 1002.738 | NA |
| LC.03.1232 | 1001.9225 | NA |
| LC.03.1233 | 985.176 | NA |

**Supplemental Table 5** Lipids and lipid-like molecules enrichment test in 26 network modules identified by WGCNA based on metabolite annotation

| | Network Module | | All metabolites | | | |
|---|---|---|---|---|---|---|
| Module | Lipids-like | Non-Lipids-like | Lipids-like | Non-Lipids-like | Pvalue | Pvalue.adj |
| red | 37 | 44 | 321 | 1373 | $9.51\times10^{-8}$ | $2.00\times10^{-6}$ |
| tan | 27 | 29 | 321 | 1373 | $1.16\times10^{-6}$ | $1.50\times10^{-5}$ |
| pink | 30 | 37 | 321 | 1373 | $2.21\times10^{-6}$ | $1.90\times10^{-5}$ |
| orange | 14 | 8 | 321 | 1373 | $6.37\times10^{-6}$ | $3.30\times10^{-5}$ |
| green | 34 | 49 | 321 | 1373 | $5.72\times10^{-6}$ | $3.30\times10^{-5}$ |
| brown | 39 | 73 | 321 | 1373 | $1.00\times10^{-4}$ | $4.33\times10^{-4}$ |
| darkred | 14 | 18 | 321 | 1373 | $1.30\times10^{-3}$ | $4.82\times10^{-3}$ |
| yellow | 31 | 75 | 321 | 1373 | $8.80\times10^{-3}$ | $2.86\times10^{-2}$ |
| lightyellow | 12 | 24 | 321 | 1373 | 0.03 | 0.09 |
| midnightblue | 14 | 32 | 321 | 1373 | 0.04 | 0.12 |
| lightcyan | 12 | 34 | 321 | 1373 | 0.15 | 0.34 |
| darkgrey | 7 | 17 | 321 | 1373 | 0.16 | 0.34 |
| greenyellow | 15 | 48 | 321 | 1373 | 0.21 | 0.42 |
| royalblue | 7 | 27 | 321 | 1373 | 0.47 | 0.88 |
| black | 0 | 76 | 321 | 1373 | 1.00 | 1.00 |
| magenta | 1 | 63 | 321 | 1373 | 1.00 | 1.00 |
| grey | 4 | 132 | 321 | 1373 | 1.00 | 1.00 |
| darkgreen | 0 | 31 | 321 | 1373 | 1.00 | 1.00 |
| lightgreen | 1 | 36 | 321 | 1373 | 1.00 | 1.00 |
| purple | 2 | 62 | 321 | 1373 | 1.00 | 1.00 |
| turquoise | 1 | 181 | 321 | 1373 | 1.00 | 1.00 |
| blue | 4 | 119 | 321 | 1373 | 1.00 | 1.00 |
| grey60 | 5 | 37 | 321 | 1373 | 0.92 | 1.00 |
| darkturquoise | 1 | 29 | 321 | 1373 | 1.00 | 1.00 |
| salmon | 2 | 50 | 321 | 1373 | 1.00 | 1.00 |
| cyan | 7 | 42 | 321 | 1373 | 0.84 | 1.00 |

Lipids-like=Lipids and lipid-like molecules;

Non-Lipids-like= Non-Lipids and non-lipid-like molecules

**Supplemental Table 6.** Percent changes in prediction accuracy of G+M over GBLUP(G) and metabolomic BLUP (M) models for the 17 traits in the Diversity panel. The first three columns are median values of prediction accuracies across 50 re-sampling runs for G, M and G+M models, respectively; the last two columns are percent changes in prediction accuracy of G+M over G and M models, respectively.

| TraitName | G | M | G+M | GM_to_G (%)[a] | GM_to_M (%)[b] |
|---|---|---|---|---|---|
| Plant Height | 0.598 | 0.539 | 0.660 | 10.4 | 11.5 |
| Days to Heading | 0.615 | 0.652 | 0.688 | 11.9 | 11.2 |
| Seed Length | 0.457 | 0.465 | 0.672 | 46.9 | 46.2 |
| Seed Width | 0.694 | 0.692 | 0.722 | 4.1 | 4.1 |
| Seed Height | 0.779 | 0.705 | 0.799 | 2.5 | 2.8 |
| Hundred Kernel Weight | 0.715 | 0.700 | 0.801 | 12.0 | 12.2 |
| Hundred Hull Weight | 0.395 | 0.436 | 0.560 | 41.7 | 37.8 |
| Groat Percentage | 0.545 | 0.534 | 0.622 | 14.2 | 14.5 |
| C14:0 | 0.421 | 0.405 | 0.628 | 49.2 | 51.1 |
| C16:0 | 0.524 | 0.803 | 0.840 | 60.4 | 39.4 |
| C16:1 | 0.612 | 0.566 | 0.651 | 6.4 | 6.9 |
| C18:0 | 0.501 | 0.784 | 0.839 | 67.3 | 43.1 |
| C18:1 | 0.550 | 0.855 | 0.878 | 59.7 | 38.4 |
| C18:2 | 0.580 | 0.805 | 0.844 | 45.4 | 32.7 |
| C18:3 | 0.358 | 0.430 | 0.520 | 45.1 | 37.5 |
| C20:0 | 0.525 | 0.843 | 0.848 | 61.4 | 38.3 |
| C20:1 | 0.609 | 0.707 | 0.768 | 26.2 | 22.5 |

[a]percent changes in prediction accuracy of G+M over G model

[b]percent changes in prediction accuracy of G+M over M model

**Supplemental Table 7** Description of the multi-trait models

| | | Covariance structure[a] | |
|---|---|---|---|
| ID | Code | G | R |
| 1 | D-D | D | D |
| 2 | D-UN | D | UN |
| 3 | UN-D | UN | UN |
| 4 | UN-UN | UN | UN |
| 5 | FA-D | FA | D |
| 6 | FA-UN | FA | UN |

[a]when fitted a multi-trait models with two kernels (G+M), G kernel and M kernel had the same genetic and residual covariance structures.
D=diagonal, UN=unstructured, FA=factor-analytic.
G=genetic covariance structure, R= residual covariance structure
